# Capillary interactions drive the self-organization of bacterial colonies

**DOI:** 10.1101/2024.05.28.596252

**Authors:** Matthew E. Black, Chenyi Fei, Ricard Alert, Ned S. Wingreen, Joshua W. Shaevitz

## Abstract

Many bacteria inhabit thin layers of water on solid surfaces both naturally in soils or on hosts or textiles and in the lab on agar hydrogels. In these environments, cells experience capillary forces, yet an understanding of how these forces shape bacterial collective behaviors remains elusive. Here, we show that the water menisci formed around bacteria lead to capillary attraction between cells while still allowing them to slide past one another. We develop an experimental apparatus that allows us to control bacterial collective behaviors by varying the strength and range of capillary forces. Combining 3D imaging and cell tracking with agent-based modeling, we demonstrate that capillary attraction organizes rod-shaped bacteria into densely packed, nematic groups, and profoundly influences their collective dynamics and morphologies. Our results suggest that capillary forces may be a ubiquitous physical ingredient in shaping microbial communities in partially hydrated environments.

## Introduction

Many bacteria live in thin layers of water wetted to solid surfaces. In partially saturated soils, cells inhabit the liquid bridges that bind soil particles together [1, 2, 3, 4]. In plants, bacteria colonize the surfaces of leaves and flowers, where thin water films and threads can form [5]. In textiles, cells adhere to wetted fibers, with the thickness of the surrounding liquid film changing as the material hydrates or dries [6]. Laboratory experiments typically use hydrogels containing liquid media to study growth and motility of microorganisms.

For nearly a century, substrate hydration has been recognized as a necessary and important factor in structuring the swarming behavior of flagellated bacteria [7, 8, 9, 10, 11, 12, 13]. For such cells, local water availability sets an allowable range within which the flagella can operate and cells can thus move. Non-flagellated, surface-adherent bacteria have demonstrated a variety of other motility mechanisms, *e*.*g*. through pilus-mediated twitching or focal-adhesion mediated gliding [14, 15, 16, 17]. Motility through these mechanisms also depends on water availability. For example, bulk twitching and gliding assays often need to be conducted in humid environments [15, 19, 21]. Recently, this link was more firmly established in *Myxococcus xanthus* where it was demonstrated that the wetting of surface heterogeneities affects cellular motility and aggregation [22]. Despite long-standing, albeit often tacit, knowledge that hydration affects the motility of surface-adherent bacteria, the mechanisms by which forces from the liquid affect these populations remain unknown.

Previous work on colloidal particles has shown that interfacial forces from water can have substantial effects on the micron scales typical of bacteria. Particles partially immersed in a thin layer of liquid experience capillary forces due to the deformation of the liquid-air interface [23, 24]. For a single hydrophilic particle, capillary forces push it into the hydrated substrate and may increase adhesion to or friction with the surface [25, 26]. When the menisci of two particles coalesce, capillary forces typically produce an attractive interaction between the particles [23]. Here, we show that this liquid-mediated interaction strongly affects the motility of gliding bacteria and governs their collective behaviors.

## Results

### Water menisci form around bacterial cells on hydrated substrates

We first sought to measure the wetting meniscus around a single cell. We spotted bacteria onto 1.5% v/v agarose hydrogels and imaged the resulting liquid-air interface around single cells with a laser-scanning confocal microscope (Keyence VK-X1000). This instrument provides both an image of the reflectance of the sample, from which cells can be identified, and the corresponding height field of the sample’s surface. We imaged the gliding bacteria *M. xanthus* and *Flavobacterium johnsoniae*, non-gliding *Escherichia coli*, and 1-µm-diameter polystyrene colloidal beads (Supplementary Table 1). In all cases, we observed a liquid meniscus surrounding the objects (Fig. 1a).

**Figure 1.**
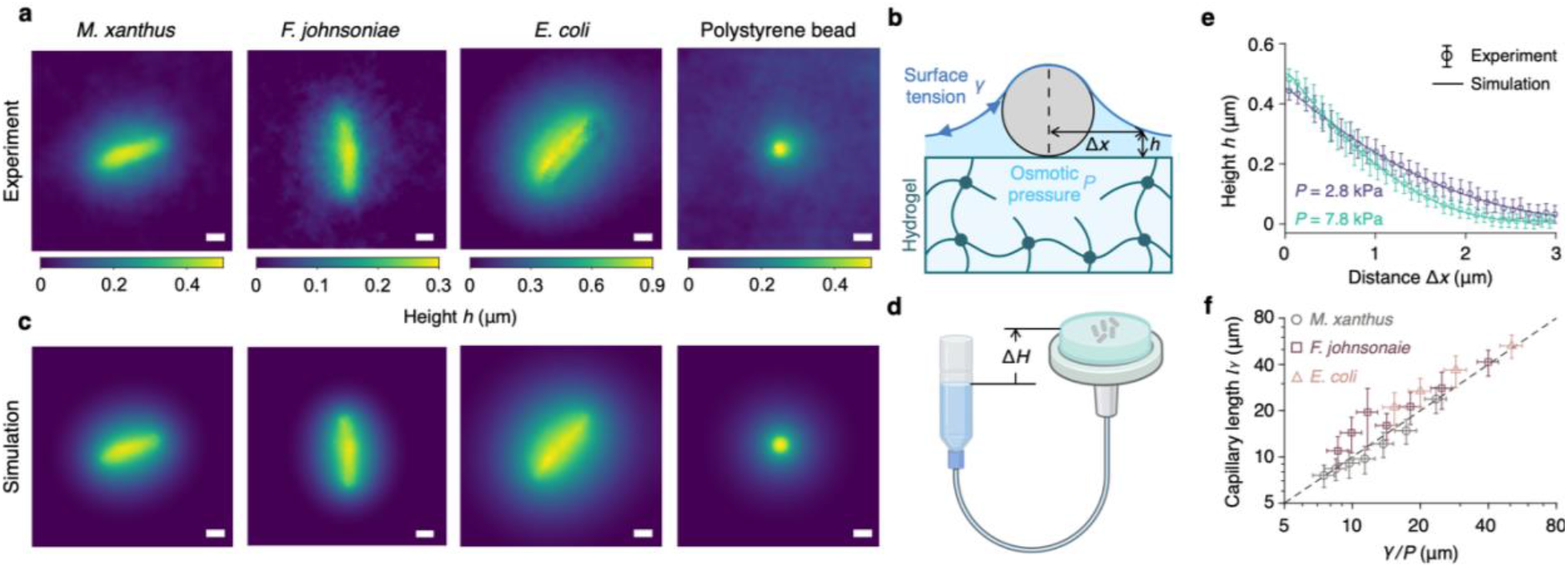
Water menisci around bacteria and colloidal particles on a hydrated substrate. **(a)** Measured height profiles around single bacteria and a 1 μm diameter polystyrene bead deposited onto an agarose hydrogel. **(b)** Schematic of the modeled water meniscus that results from the balance of surface tension (dark blue) and osmotic pressure difference P (light blue) across the surface of the hydrogel substrate (green meshwork). The model assumes that the water meniscus completely wets the surface of a cell (gray circle with black outline; cross-section). h denotes the height of water above the substrate surface and Δx denotes the horizontal distance to the midline of the cell. See Methods for details. **(c)** Height profiles of the modeled wetting menisci. Scale bar is the same for all images and represents 1 μm. **(d)** Illustration of the experimental device used to control water pressure. A reservoir of media is connected to the bottom of the agarose hydrogel onto which cells are deposited for each experiment. The height difference, ΔH, between the top of the hydrogel surface and the top of the reservoir sets an additional hydrostatic pressure that tunes the net osmotic pressure difference across the hydrogel. **(e)** Water height profiles h(Δx), measured in the midplane perpendicular to the long axis of the cell, at the designated values of P. The reference value P_0_ is estimated by fitting the height profile at ΔH = 0, and P is inferred using the relation P = P_0_ + *ρ*_w_gΔH. Circles with error bars indicate experimental measurements (mean +/-standard deviation). Solid curves indicate fits of simulation results. **(f)** Capillary lengths inferred from the height profiles of the wetting menisci at varying values of P. Different colors and symbols represent the designated bacterial species. Black dashed line indicates the line y = x. See Supplementary Information for details.

For a hydrated substrate such as agarose, we modeled the meniscus profile *h*(*r*) = *h*(*x, y*) by minimizing the energy of the wetting liquid with surface tension *γ* and an energy penalty for extracting water from the substrate due to the osmotic pressure difference *P* across the substrate surface (Fig. 1b). We computed the equilibrium height profile of the modeled water meniscus via numerical minimization of a total free energy 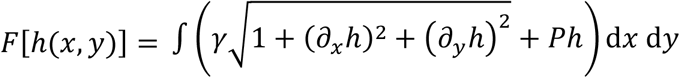, assuming that the water meniscus completely wets the cell surface (Supplementary Sec. I.A). These results are in agreement with the measured height profiles around all samples tested (Fig. 1c).

### The shape of a water meniscus varies with osmotic pressure

Our model predicts that the width of the water meniscus around the cell is controlled by the capillary length 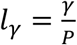. Specifically, for an infinitely long cylindrical cell of radius *R*_c,_ the non-zero principal curvature of the surrounding water meniscus is 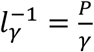, and thus its width scales with 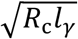, which arises from the geometry of two tangent circles of radii *R*_c_ and *l*_*γ*_ (Supplementary Figs. 1 and 2, and Sec. I.B).

To test this prediction for the meniscus width, we developed an apparatus to control the osmotic pressure difference *P*. Similar to devices for controlling water availability in model soils [27, 28], the apparatus works by connecting the agarose gel directly to a liquid reservoir that, like the gel surface, is open to the atmosphere (Supplementary Fig. 3 and Sec. VI.B). The height of the gel surface (*H*_gel_) relative to the height of the reservoir (*H*_res_), Δ*H* = *H*_gel_ − *H*_res_, can then be used to apply a hydrostatic pressure to the gel Δ*P* = *ρ*_w_*g*Δ*H*, where *ρ*_w_*g* = 9.8 kPa/m is the specific weight of water (Fig. 1d, and Supplementary Fig. 4). We set Δ*H* = 0 as the reference state for each condition and obtained the osmotic pressure, *P*_0_ ≈ 2 kPa, of this state by fitting the modeled water height profiles to those measured in experiment (Supplementary Fig. 5, Sec. I.C,D, and Table 2). The osmotic pressure difference *P* across the substrate surface is given by *P* = *P*_0_ + *ρ*_w_*g*Δ*H*, (Supplementary Fig. 6 and Sec. I.E). We varied Δ*H* and measured the meniscus shape around cells of variable size, shape, and aspect ratio. The surface tension of all media used in the experiments was measured by the pendant drop method (Supplementary Sec. VI.C and Table 3). We set *γ* = 66 mN/m when fitting the experimental profiles in simulation. As our model predicts, the width of the wetting meniscus, which correlates with the capillary length 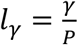, becomes smaller when *P* is increased due to the larger free-energy cost of extracting water from the substrate surface (Fig. 1e,f).

### Capillary forces facilitate the formation of cell groups

For an isolated cell, the lateral capillary forces around the cell balance one another. However, when the menisci of adjacent cells overlap with each other, the water profile around each cell becomes asymmetric. This gives rise to a lateral attractive capillary force between the cells (Fig. 2a). Higher osmotic pressures, *P*, corresponding to lower water availability, should lead to higher lateral forces but also smaller menisci and thus a shorter interaction range. We confirmed this intuition by coupling the free-energy model of water to an agent-based model of bacterial cells. We model the cells as horizontally oriented spherocylinders with radius *R*_c_ and cylindrical length *l*_c_ and compute the forces on the cells exerted by the simulated water menisci (Supplementary Figs. 7 and 8, and Sec. I.F,G). Using this model, we simulated the merging of two adjacent cells being pulled together by capillary forces for different values of the osmotic pressure (Fig. 2b,c). We next asked whether our model was sufficient to capture the merging dynamics of non-motile *M. xanthus* cells. We consider the balance of forces and torques exerted on each cell, including those from cell-water interaction, cell-cell steric repulsion, and cell-substrate friction (Supplementary Sec. II). Following Copenhagen et al. [29], we model the friction between *M. xanthus* cells and the underlying substrate as anisotropic with smaller friction coefficient along the cell-body axis (*ξ*_∥_) than perpendicular to it (*ξ*_⊥_). We determined the values of these coefficients (Supplementary Table 2) by fitting the agent-based simulations to the dynamics of an experiment in which non-motile *M. xanthus* cells deposited onto an agarose gel were pulled together, into a side-by-side configuration, by their overlapping menisci (Fig. 2d, and Supplementary Video 1). The fitted model recapitulates both the translational (Fig. 2e) and rotational (Fig. 2f) dynamics of these merger events, and supports the anisotropic friction proposed in ref. [29] (Supplementary Fig. 9).

**Figure 2.**
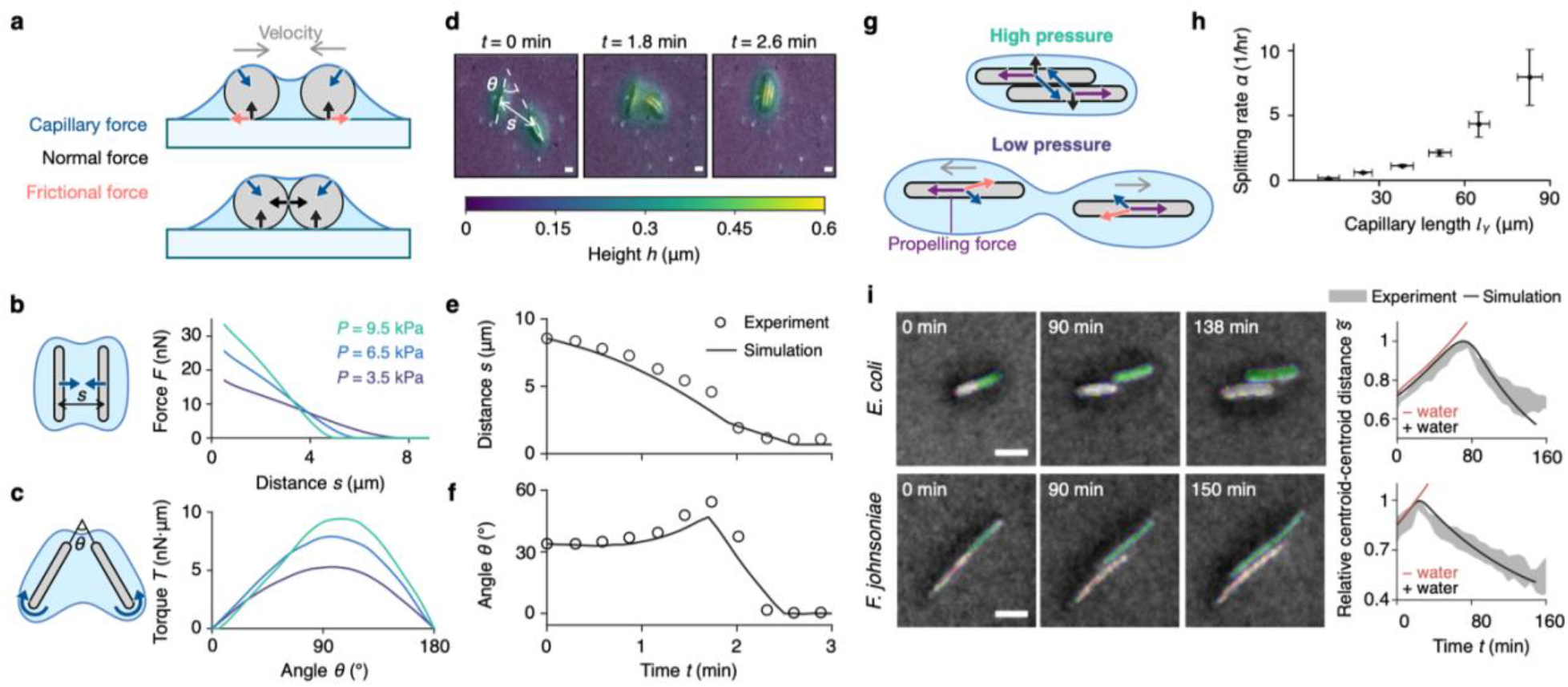
Capillary forces promote mergers and hinder separation of bacterial cells. (**a**) Cross-section schematics showing the forces exerted on non-motile cells. Color code as in Fig. 1**b**. Black arrows indicate normal forces exerted by either the substrate or the cells. Dark blue arrows indicate capillary forces. Pink arrows indicate frictional forces between the cells and the substrate. Gray arrows indicate cell velocities. Two nearby cells with overlapping water menisci (top) are pulled together by capillary forces (bottom). (**b**) Left: top-view schematic of two parallel cells at a distance, s. Color code as in **a**. Right: magnitude of the capillary force at varying distances s for the designated values of osmotic pressure, P. (**c**) Left: top-view schematic of two nearby cells with a fixed center-to-center distance and forming an angle θ. Color code as in **a**. Right: magnitude of the capillary torque at varying angles for the designated values of osmotic pressure, P. The center-to-center distance is set to 5.1 μm. In **b** and **c**, forces and torques are computed for cells with radius R_c_ = 0.3 μm and length l_c_ = 4.5 μm. (**d**) Water height profiles, overlaid on reflectance images of M. xanthus cells (ΔcglBΩpilA) that merge due to overlapping wetting menisci. Scale bars: 1 μm. (**e, f**) Time evolution of (**e**) center-to-center distance s and (**f**) relative orientation angle θ, as indicated for the two cell groups in **d**. Circles indicate experimental measurements and solid curves indicate fitted results of agent-based simulations. (**g**) Top-view schematics of the forces exerted on self-propelling cells for high osmotic pressure (top) and low osmotic pressure (bottom). Color code as in **a**. Purple arrows indicate self-propelling forces along cell’s long axis. (**h**) The rate at which pairs of motile wild-type M. xanthus cells separate apart, as schematized in **g**, increases with the measured capillary length. Error bars are boot-strapped standard deviation. (**i**) Stereotyped dynamics of non-motile, dividing cells due to capillary forces. Post-division pairs of cells undergo a period of end-on-end growth before capillary forces trigger them to slide relative to each other. Example pairs are shown for E. coli and F. johnsonaie (ΔsprB) and the centroid-to-centroid distance, relative to its largest value for each pair, is plotted on the right. Relative centroid-centroid distances are averaged over N=5 and N=10 cells for E. coli and F. johnsonaie, respectively. Shaded regions represent mean +/-the standard deviation of the experimental data. Black curves and red curves represent the simulation results with and without water, respectively. Model parameters used in **e, f**, and **i** are listed in Supplementary Table 2.

Similar rearrangements also occur for motile bacteria, giving rise to formation of pairs and larger groups when adjacent cells move near one another (Supplementary Video 2). Given the attractive capillary forces between adjacent cells (Supplementary Fig. 10), we reasoned that the strength of such forces would anti-correlate with the rate at which two motile cells would split apart from one another. Measurements of the splitting rate (number of events/total time tracked) of wild-type *M. xanthus* cell pairs across a range of capillary lengths confirmed this hypothesis (Fig. 2g,h). Similarly, for large groups, the magnitude of group shape fluctuations is inversely related to the strength of capillary forces (Supplementary Figs. 11 and 12).

For non-motile but growing bacteria, we hypothesized that similar dynamics driven by capillary forces could account for the side-by-side alignment of daughter cells after division. To test this, we deposited exponentially growing but non-motile *E. coli* and *F. johnsoniae* on agarose pads and collected timelapse micrographs of pairs of daughter cells formed by the division of an isolated, single cell (Fig. 2i, left). Subsequently, we segmented images using a convolutional neural network and tracked individual daughter cells over time. As hypothesized, pairs of cells of both species exhibited similar dynamics: cells that were initially arranged end-to-end first slid sideways and then longitudinally to align in a side-to-side configuration. To quantify this process, we tracked the centroid-to-centroid distance between pairs of cells, which first increased due to growth as the cells slid sideways, and then decreased as the cells were pulled toward each other by the capillary force (Fig. 2i, right, and Supplementary Figs. 13 and 14).

### Capillary forces and cell motility lead to different phases of collective motion

We used the above data to calibrate an extended version of our agent-based model where cells self-propel by generating in-plane forces, ***F***_prop_, along their long axis, and reverse directions stochastically with mean reversal time *τ*_r_. Since the capillary force acting laterally on a cell can be on the order of 10 nN (Fig. 2b), much stronger than the forces generated by bacterial motility motors (10∼100 pN) [30, 31, 32], we explored how capillary forces might affect the collective dynamics of motile cells. We performed a series of simulations for motile but non-growing cells, fixing their density and aspect ratio while varying three dimensionless variables: a normalized reversal period 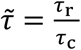 where 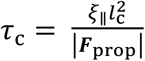 is the time it takes for a self-propelled cell to travel its body length, a normalized self-propulsion strength 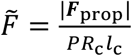 where *PRc l*_c_ is the scale of the horizontal capillary force acting on a cell, and a normalized capillary length 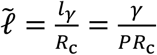.

For the simulation parameters we investigated, the collective dynamics are primarily determined by the cell-motility variables 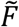 and 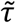, with only a minor dependency on 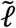. When the self-propulsion force is much stronger than the capillary force, 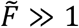, the cells mainly interact with one another sterically and thus behave like a nematic gas (Fig. 3a, gas, and Supplementary Video 3). In the opposite limit 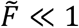, when capillary forces dominate the dynamics, groups of cells cannot split apart. The population thus rapidly coarsens into “droplets” of cells (Fig. 3a, droplets, and Supplementary Video 4). However, for intermediate regimes 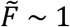, we observe dynamic cell groups which are capable of both merging and splitting. In this regime, the structure and dynamics of these groups are determined by the cell’s reversal period. For very long reversal times 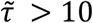, polar clusters form (Fig. 3a, polar clusters, and Supplementary Video 5). At shorter reversal times 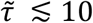, cells propelling in opposite directions can pass by each other but are typically unable to escape the range of capillary attraction before reversing their direction of motion. Here, quasi-one-dimensional streams form (Fig. 3a, streams, and Supplementary Video 6). When the reversal time becomes shorter than the time for a cell to move the length of its body, the droplet phase is recovered (Fig. 3a, droplets). We identified these different phases by computing the cell-axis-aligned, equal-time spatial correlation 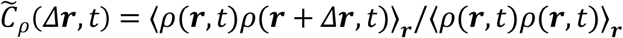 of cell density *ρ* (Fig. 3c) and using the mean and variance of these correlations to classify the dynamics on a phase diagram (Fig. 3e, and Supplementary Fig. 15 and Sec. III).

**Figure 3.**
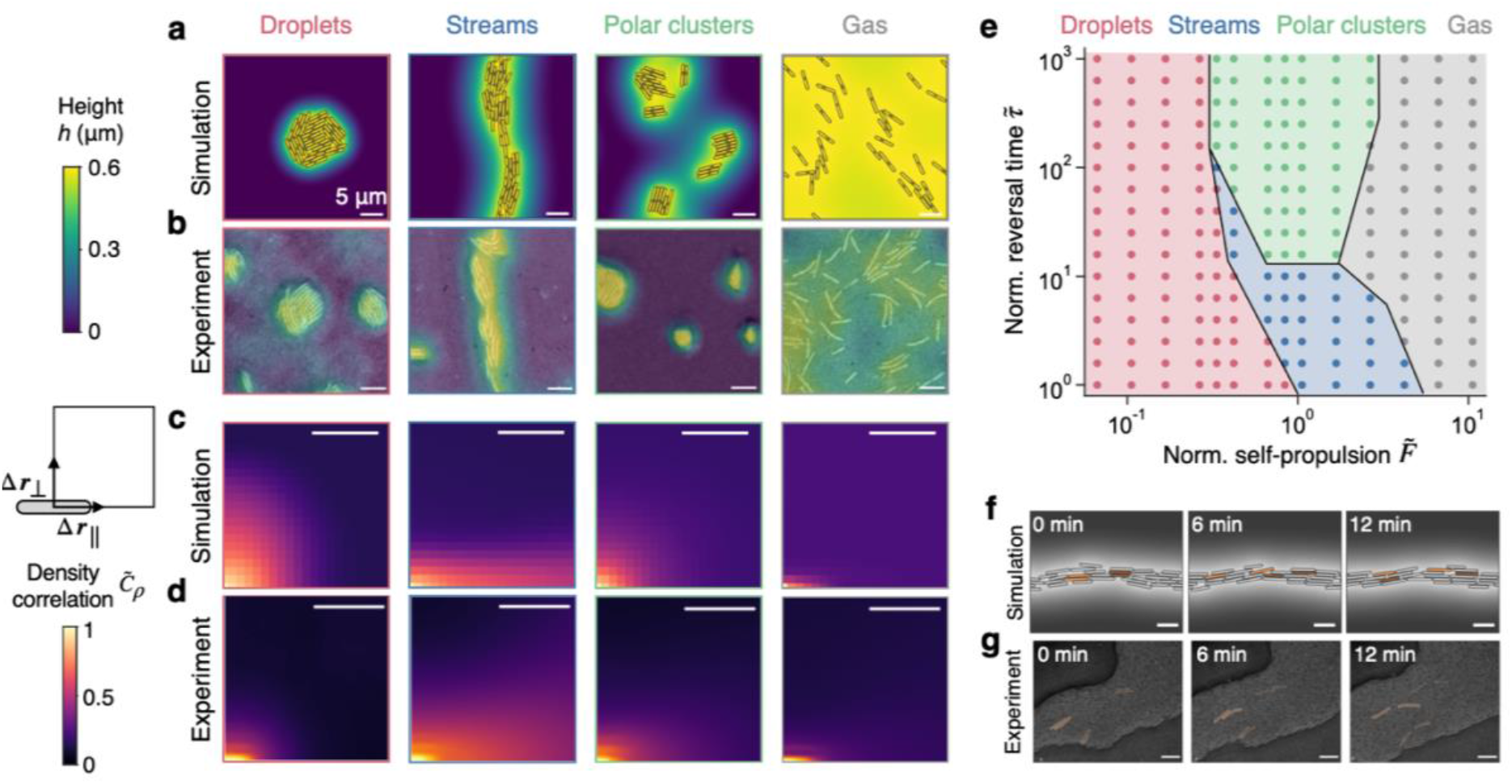
Capillary attraction and cell motility lead to multiple phases of collective cell organization. (**a**,**b**) Heatmaps of water-height profiles for typical snapshots of the designated phases in (**a**) simulations and (**b**) experiments. (**c, d**) Time-averaged steady-state spatial density autocorrelation for the designated phases in (**c**) simulations and (**d**) experiments. (**e**) A section of the phase diagram for simulated cells. Color code as in **a**. Phase boundaries are drawn by hand. The normalized capillary length is set to 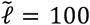. **(f, g)** Snapshots of (**f**) simulation and (**g**) experimental timelapse in which adjacent cells in a stream are labeled by the same color at time t=0 min and tracked over time. Adjacent cells can slide relative to one another, allowing the pairs to move apart and exchange neighbors. In all panels, scale bars: 5 μm.

To explore the phase diagram experimentally, we captured time-lapse micrographs of various strains of *M. xanthus* on agarose hydrogels filled with rich media (CTT) and maintained at constant osmotic pressure with the previously described device. We segmented the resulting images using a convolutional neural network, and used the resulting cell masks to compute the mean and variance of 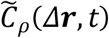 (Fig. 3b,d, and Supplementary Fig. 16) over 3-hour windows. To achieve 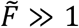, we used our device to reduce the osmotic pressure *P* and flood the hydrogel surface. In this case, wild-type populations show qualitative similarity to the gas phase (Fig. 3b,d, and Supplementary Video 7). In the opposite limit, a non-motile strain of *M. xanthus*, corresponding to 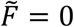, forms cellular droplets as predicted by our simulations (Fig. 3b,d, and Supplementary Video 8). Also consistent with the simulation results, wild-type cells form stable, elongated streams, while the hypo-reversing Δ*frzZ* cells form polar clusters (Fig. 3b,d, and Supplementary Videos 9 and 10). These phases of nematic streams and polar clusters have also been reported in previous studies of *M. xanthus* [33, 34, 35]. Overall, our experiments showed all the phases predicted by the simulations.

Additional experiments using a mutant strain of *M. xanthus* lacking pili (Δ*pilA*) did not reveal stable streams, suggesting that additional interactions may be necessary to maintain these structures in the wild-type [36, 37, 38]. Similar to the results of Anderson and Ordal [20] wherein gliding of flavobacteria was dependent on a high degree of water availability, experiments in *F. johnsoniae* revealed only “gas” and “droplet” phases with the population rapidly transitioning between the two as water was moved into or out of the gel (Supplementary Fig. 16).

As would be expected for a “gas,” gliding cells on saturated hydrogels can move freely relative to their nearest neighbors and appear to do so in an unstructured manner (Supplementary Figs. 15 and 16). Conversely, cells in “droplets” are unable to readily pass each other because they are bound together by strong capillary forces (Supplementary Figs. 17 and 18). Interestingly, both the streams and polar clusters allow the cells to be tightly packed into spatially segregated groups while also moving relative to their neighbors (Fig. 3f,g, and Supplementary Video 11). These features might be crucial for the large-scale collective cell movement involved in the development of *M. xanthus* colonies (see Discussion).

### Capillary forces influence the organization of bacterial populations

To demonstrate how capillary forces influence the organization of bacterial populations, we used our device to impose changes in water availability on populations of *M. xanthus* and *F. johnsoniae* (Fig. 4a, and Supplementary Fig.19, and Videos 12 and 13). We tracked changes in population structure by computing the Delaunay triangulation between cell centroids (Fig. 4b, schematic). The distribution of edge lengths reveals both the relevant length scale(s) in the population – typically the approximate intercellular and intergroup spacings – as well as their heterogeneity across the population. Due to the initially low water availability in the substrate, cells are packed into small, dense groups. We then used our device to gradually saturate the substrate with water. This caused the surface to flood and the groups to break up, giving rise to a gaseous phase with a single apparent peak in the distribution of Delaunay edge lengths (Fig. 4b). Subsequently lowering the water reservoir below the substrate surface reimposed low water availability as in the start of the experiment. This change in water availability forced the cells back into densely packed groups (Fig. 4a,b).

**Figure 4.**
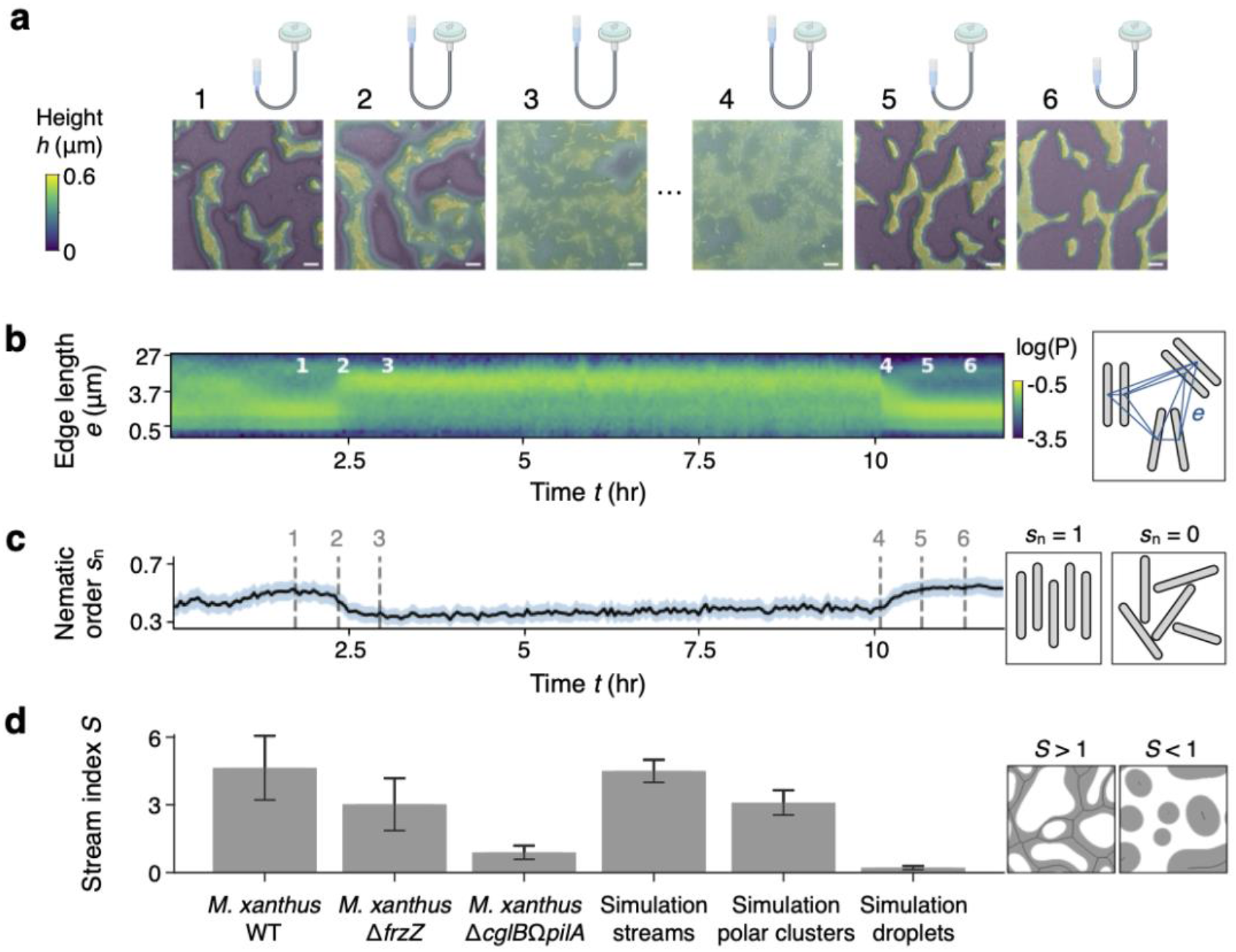
Water availability controls the organization of colonies of M. xanthus. (**a**) Experimental time series in which cells initially deposited onto an agarose pad are subjected first to increasing water availability until a gas phase is reached (snapshots 1-3). Hours later, water is drained from the substrate, and the colony goes back to forming tightly packed, nematically ordered groups and streams (snapshots 4-6). Time points (1-6) are marked in **b** and **c**. (**b**) Kymograph of the probability density function (P) of Delaunay edge lengths over time for the time series shown in **a**. Schematic on the right demonstrates how edges (blue lines) are calculated for a small population of cells (gray). At early times and late times in the experiment, low water availability at the surface constrains cells into tightly packed, well-separated groups. At intermediate times, high water availability permits cells to be further spaced apart and homogenizes the population. (**c**) Strength of nematic order s_n_, calculated within groups of 8 nearest-neighboring cells and averaged over the population at each time. Shaded regions indicate variance. Right: schematics showing a small population of cells with perfect nematic order (s_n_=1) or no order (s_n_=0). (**d**) Stream indices for wild-type M. xanthus, a hypo-reversing mutant of M. xanthus (ΔfrzZ), a non-motile mutant of M. xanthus (ΔcglBΩpilA), and simulated streams, polar clusters, and droplets. Error bars for the experimental results are standard deviations for each corresponding mean calculated over N=25, 9, and 7 frames taken from 2, 3, and 1 separate timelapses, respectively. Error bars for the simulation results are standard deviations for each corresponding mean calculated over N=50 frames during the steady state of 1 timelapse. Right: schematics showing populations that form streams (S > 1) or droplets (S < 1).

Given the rod shape of individual cells, the packing into dense groups induces local nematic ordering (Fig. 4c). In the gas phase, nematic order is reduced as cells move in non-straight paths and only interact sterically for brief periods of time [39]. Our data shows that capillary attraction promotes dense cell packing and hence the emergence of nematic order in colonies of rod-shaped bacteria.

To further explore this idea, we asked how the morphology of both motile and non-motile cell groups was affected by capillary forces. To quantify the degree of elongation of cell groups, we calculated a stream index, 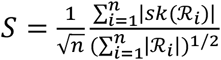 where *n* is the number of groups, |ℛ_*i*_| is the area of the *i*^th^ group, and |*sk*(ℛ_*i*_)| is the length of the morphological skeleton of the *i*^th^ group (Fig. 4d, schematic, and Supplementary Fig. 20 and Sec. IV). For both *M. xanthus* and *F. johnsoniae*, motile cell groups had higher stream indices than their non-motile counterparts, *M. xanthus* Δ*cglB*Ω*pilA* and *F. johnsoniae* Δ*sprB* (Fig. 4d, Supplementary Fig. 19). However, *M. xanthus*, whose cells exhibit periodic direction reversals, formed even more elongated structures than non-reversing *F. johnsoniae*, whose stream index was similar to that of the hypo-reversing Δ*frzZ* mutant of *M. xanthus* (Supplementary Fig. 19). The stream indices of the reversing, non-reversing, and non-motile cell groups are captured by simulations of a continuum model in the streams, polar clusters, and droplets phases, respectively (Fig. 4d and Supplementary Sec. IV).

## Discussion

In total, our work suggests that lateral capillary forces promote the aggregation of terrestrial bacteria by condensing individual cells into tightly packed groups while allowing for relative motion of neighboring cells. These features allow motile bacterial cell groups to behave as active nematic liquid crystals, whose properties cells can exploit as part of their lifestyle [29, 40]. In *M. xanthus*, cells form collective groups to swarm, predate, and aggregate under different environmental conditions [41]. When nutrients are limited, topological defects of cell alignment in the starving swarm promote the formation of new cell layers [29], which stack on top of one another to form a nascent fruiting body [42]. While we focused on lateral motion in the current work, the capillary force also contains a strong vertical component (∼100 nN on isolated cells) that pushes cells downward and may deform the underlying substrate. The same vertical capillary force opposes the extrusion of cells into new layers above the substrate, such that layer formation requires collective fluctuations in the vicinity of topological defects [40].

Nascent fruiting bodies are connected by quasi-one-dimensional streams [43, 44]. These streams may accelerate the growth of fruiting bodies by providing pathways that guide cells to coalesce [35, 45, 46]. Our results indicate that capillary attraction may play a role in the developmental process by facilitating the population’s condensation into nematic phases and the formation of inter-fruiting body streams. In the future, we will further investigate how vertical capillary forces influence the three-dimensional structure and developmental dynamics of bacterial colonies.

Our combined experimental and modeling results showed that water availability, which tunes the range and strength of capillary forces, controls the organization of bacterial colonies and gives rise to various collective phases formed by gliding bacteria on agarose hydrogels. Bacteria are known to produce surfactants that reduce the surface tension of the surrounding liquid [47, 48, 49], although whether or not this process is regulated and how it could affect the capillary forces we explored in this work remain unknown. More broadly, future work will be needed to understand how capillary forces affect bacterial growth, motion, and population morphology in more complex geometries such as those found in textiles and soil.

## Acknowledgments

We thank members of the Shaevitz and Wingreen groups for helpful discussions. We thank Dr. Mark McBride for the gift of *F. johnsoniae* strains and for helpful discussions. We thank Dr. Lotte Søgaard-Andersen for the gift of *M. xanthus* strain SA3985. We thank John D. McEnany, Sebastian González-LaCorte, and Siddhartha Jena for their valuable contributions to the early stage of this project. This work was supported in part by the National Science Foundation through the Center for the Physics of Biological Function (PHY-1734030) and award PHY-1806501, and in part by NIH grant GM082938 (N.S.W.).

## Supplementary Information for

### I MODELING WATER MENISCI AROUND CELLS

#### A A free-energy model for the water meniscus

Laser-scanning microscopy reveals that there are intensity halos around *M. xanthus* cells grown on agar and these halos display continuously decreasing height from the centerlines of the cells toward the agar surface far from the cells (main Fig. 1). Our main hypothesis is that these halos correspond to water menisci formed around the cells. To account for the shape of the water meniscus, we propose a minimal model based on a free energy that takes into account the surface energy of the air-water interface and the energy penalty of extracting water from the agar substrate. Denoting by *h*(*x, y*) the height of water above the surface of the agar substrate, the total free energy of the water is given by

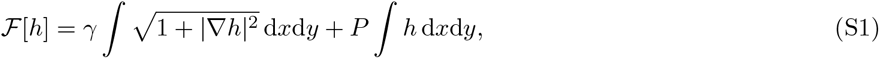

where *∇* = (*∂*_*x*_, *∂*_*y*_), *γ* denotes the surface tension of water, and *P* denotes the per-volume energy penalty of extracting water out of the substrate, i.e., the osmotic pressure difference across the substrate surface (see Sec. I E for details). Note that *γ* and *P* together yield a capillary length *l*_*γ*_ = *γ/P* that controls the range of capillary interactions (see Sec. I G). We assume that the equilibrium shape of the water meniscus *h*^*∗*^(*x, y*) minimizes the free-energy functional Eq. (S1), subject to the constraint that the height of water cannot be lower than the upper cell surfaces *z*_upper_ or the substrate surface *z* = 0. To express this constraint mathematically, we next specify the geometry of the modeled cells.

We treat each cell as a spherocylinder with cap radius *R*_c_ and cylindrical length *l*_c_. The position of cell *i* is determined by its center-of-mass coordinates 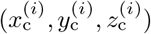 and its orientation 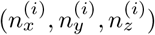. For simplicity, we assumed that cells are parallel to the substrate surface and the surface is tangent to the cylindrical part of the cell, i.e.,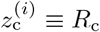 and 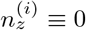. Thus, the position of each cell *i* is determined by its center-of-mass coordinates 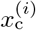 and 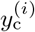, and its orientation angle 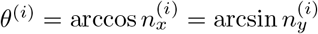. The height profile of cell *i*’s upper surface is given by

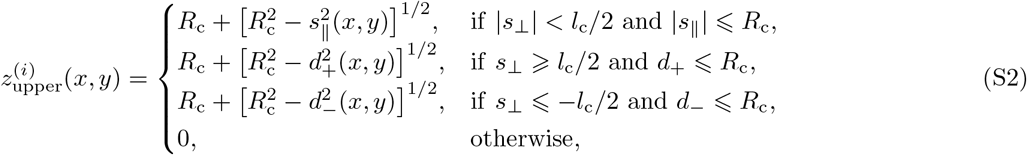

where 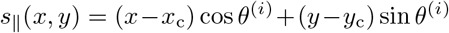 and 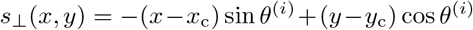 denote, respectively, the coordinates of (*x, y*) along or perpendicular to the cell’s long axis, and 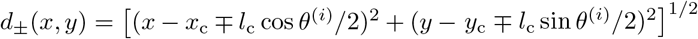 denotes the distance from (*x, y*) to the ± ends of the cell. Therefore, the constraint on *h* can be expressed as

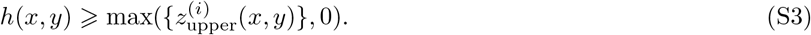

#### B Examples of equilibrium height profiles

We perform the constrained energy minimization described in Sec. I A numerically to solve for the equilibrium height profiles. In the simulations, we vary the normalized capillary length 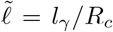 to change the width of the water meniscus. To verify our simulation results, we consider two special cases below where analytical solutions can be obtained.

##### Water meniscus around a 2D circle

We start by considering a case of reduced dimensionality in the plane *y* = 0. We consider a 2D circular “cell” of radius *R*_c_, whose center is located at *x* = 0, *z* = *R*_c_. In this case, the total free energy in Eq. (S1) is given by 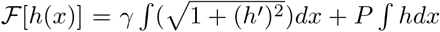 where we define *f*^*′*^ ≡ *df /dx* for an arbitrary function *f* (*x*). The free-energy minimum is achieved when the functional derivative *δ*ℱ*/δh* = 0, which yields

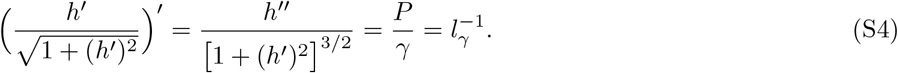

The solution of Eq. (S4) is a circular arc of radius *l*_*γ*_. Taking into account that the height profile of water must be continuous, we obtain^1^

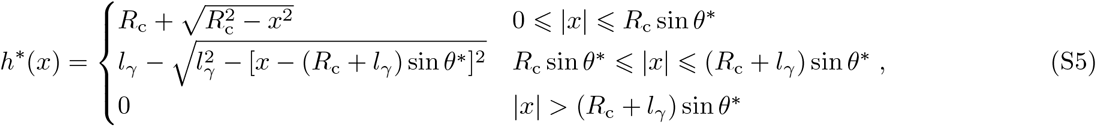

where 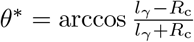. Our simulation results match closely with the analytical solution (Supplementary Fig. 1).

##### Water meniscus around a 3D sphere

A similar procedure as above can be performed to compute the shape of the water meniscus around a 3D spherical “cell” of radius *R*_c_ whose center is located at *x* = 0, *y* = 0, *z* = *R*_c_. Due to polar symmetry, the water height profile *h* = *h*(*r*) is only a function of *r* = (*x*^2^ + *y*^2^)^1*/*2^. The free-energy functional is given by 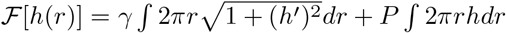, and the optimization *δ*ℱ*/δh* = 0 yields

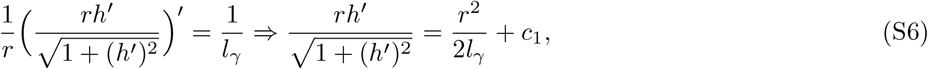

where *c*_1_ is an integration constant. The explicit expression for the solution of Eq. (S6) is cumbersome and contains multiple elliptical functions. Nevertheless, *c*_1_ can be determined from *h*^*′*^ = 0 at *r* = *r*_max_ ≡ argmin[*h*(*r*) = 0], and Eq. (S6) can thus be rewritten as

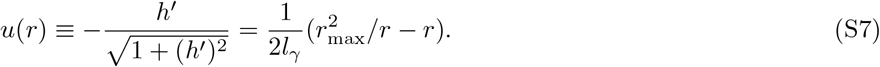

Indeed, the simulation results are in good agreement with this analytical expression (Supplementary Fig. 1).

#### C Estimating the capillary length from the equilibrium height profiles

In this section, we discuss how to estimate the capillary length *l*_*γ*_ from the measured height profile of the water meniscus in experiment (Fig. 1). We measured the height profile *h*^*∗*^(*d*) in the midplane perpendicular to the long axis of the cell, where *d* denotes the horizontal distance to the long axis. Note that for a spherocylindrical cell that is located at *x*_c_ = 0, *y*_c_ = 0 and whose orientation angle is *θ* = *π/*2, we have *h*^*∗*^(*d*) ≡ *h*^*∗*^(*x* = ±*d, y* = 0).

As discussed in Sec. I B **Water meniscus around a 2D circle**, for an infinitely long cell, *h*^*∗*^(*d*) corresponds a circular arc of radius *l*_*γ*_, and thus *l*_*γ*_ can in theory be estimated from the radius of curvature *R*_*κ*_ of *h*^*∗*^(*d*), i.e., 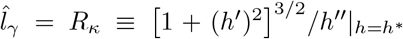 In practice, there are two problems: (1) calculating derivatives of a noisy measurement *h*^*∗*^(*d*) can be unreliable, and (2) the experimental cells are not infinitely long. To address the first problem, we measure the distance *d*_1*/*2_ at which *h*^*∗*^ drops below *R*_c_*/*2 and estimate the radius of curvature *R*_*κ*_ to be 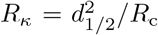 assuming that *R*_*κ*_ ≫ *R*_c_ (see Supplementary Fig. 2a for the geometry). We further verified that, for all the bacterial cells tested, the condition *R*_*κ*_ ≫ *R*_c_ is satisfied (main Fig. 1). To address the second problem, we simulate the equilibrium height profiles for varying values of cell aspect ratio *l*_c_*/*(2*R*_c_) and capillary strength (*PR*_c_)*/γ*, and compute 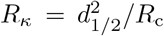 for the simulated profiles (Supplementary Fig. 2b). We find that, indeed, *R*_*κ*_ = *l*_*γ*_ when *l*_c_*/*(2*R*_c_) =∞ and *R*_*κ*_ = *l*_*γ*_*/α*_corr_ (*α*_corr_ *>* 1) for finite *l*_c_*/*(2*R*_c_). Thus, we infer the capillary length from the experimental measurements using 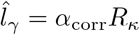 (with *α*_corr_ obtained from Supplementary Fig. 2b) and ignore the uncertainty of the correction factor *α*_corr_ for simplicity.

#### D Model parameters

We estimate the model parameters in our model of water meniscus directly from experiment:

##### Cell length and radius

We estimate the cap radius *R*_c_ by half the maximal height measured in experiment. The cylindrical length *l*_c_ is determined from the bright-field images of bacterial cells. We choose *R*_c_ = 0.3 µm and *l*_c_ = 4.5 µm for *M. xanthus* cells, *R*_c_ = 0.4 µm and *l*_c_ = 2.4 µm for *E. coli* cells, and *R*_c_ = 0.15 µm and *l*_c_ = 2.7 µm for *F. johnsoniae* cells. For the full simulation of *M. xanthus* cells, we set *R*_c_ = 0.3 µm and we draw *l*_c_ from a uniform distribution U[3.6 µm, 5.4 µm].

##### Water surface tension

The surface tension *γ* of the liquid media is measured experimentally. See Sec. VI C for details. The values of *γ* are reported in Table 3.

##### Osmotic pressure

To estimate the values of *P* under different conditions, we assume that *P* = *P*_0_ + **ρ**_w_*g*Δ*H*, where **ρ**_w_ denotes the mass density of water, *g* denotes the gravitational acceleration constant, Δ*H* denotes the height difference between the gel surface and the water level of the syringe connected to the gel (Fig. 1d), and *P*_0_ denotes the osmotic pressure difference across the gel surface at Δ*H* = 0. The value of *P*_0_ is determined by fitting the height profiles of the modeled water menisci to those measured in experiments with Δ*H* = 0. We consider two fitting procedures:

1. We compare the 3D height profiles *h*(*x, y*) of the water menisci around *N* individual cells between experiment and theory. For each individual cell *i*, we denote by 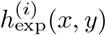 and 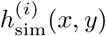 the experimental measurements and the simulation results, respectively. Since the experimental cells are not perfect spherocylinders, we minimize the free energy Eq. (S1) using the constraint

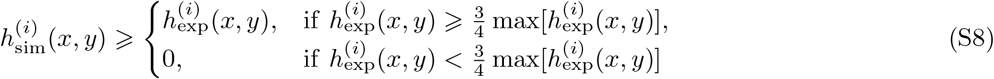

to obtain the simulated equilibrium water meniscus 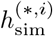. The fitting error is measured by the root-mean-square height difference 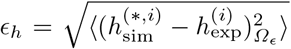 on the region 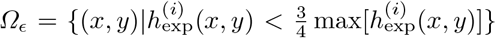. We vary the value of *P*_0_ and obtain the optimal fitting value 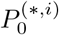 that minimizes *ϵ*_*h*_. Finally, the value of *P*_0_ is determined by 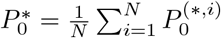 and the uncertainty of *P*_0_ is given by 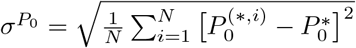
2. Alternatively, we compare the modeled and measured water height profiles in the midplane perpendicular to the long axis of the cell (Fig. 1e). We average the measured height profiles over multiple cells at various distances *d* from the cell midline to obtain 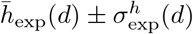. We set 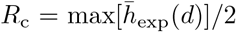, and simulate the equilibrium height profile 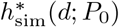 with varying values of *P*_0_. To quantify the fitting error, we compute the negative log-likelihood function 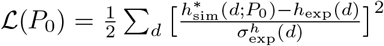. Subsequently, the value of *P*_0_ is determined by 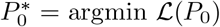 and the uncertainty 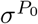 satisfies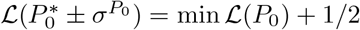.

As shown in Supplementary Fig. 5, the two fitting procedures lead to similar estimations of *P*_0_. Specifically, fitting the full 3D height profile yields *P*_0_ = 2.69 ± 0.61 kPa and fitting the midplane 2D height profile yields *P*_0_ = 2.81 ± 0.29 kPa.

#### E Additional notes on the osmotic pressure difference across the gel surface

In Sec. I D, we find that there is a nonzero osmotic pressure difference *P*_0_ across the gel surface even when Δ*H* = 0. We hypothesize that there is a vertical osmotic pressure gradient across the gel due to evaporation –– the top of the gel has a higher polymer volume fraction, and hence a higher osmotic pressure, than the bottom of the gel (Supplementary Fig. 6a). In this subsection, we explore whether this gradient can account for a nonzero *P*_0_. Since the gel is able to swell or shrink, we consider two configurations of the gel, one that represents the reference configuration and one that corresponds to the current configuration. We denote by *Z*_0_ and *z* the vertical coordinates in the reference configuration and the current configuration, respectively.

##### Governing equations

Consider an infinitesimal horizontal slice of the gel that is located at [*Z*_0_, *Z*_0_ + d*Z*_0_] in the reference configuration and at [*z, z* + d*z*] in the current configuration. Since the volume of polymers in this slice of the gel is conserved, we obtain *ϕ*_0_d*Z*_0_ = *ϕ*d*z*, where *ϕ*_0_ and *ϕ* denote, respectively, the volume fraction of polymers in the reference and current configurations, and we assume *ϕ*_0_ to be independent of *Z*_0_ for simplicity. In addition, the mass conservation of water yields *∂*(Δ*V*_w_)*/∂t* = (*J*|_*Z*_0 − *J*|_*Z*_0 +d*Z*0)*S*, where the volume of water in this gel slice can be expressed as Δ*V*_w_ = *S*d*Z*_0_(*ϕ*_0_*/ϕ − ϕ*_0_), *S* denotes the surface area of the gel, and *J* denotes the flux of water. In the current configuration, *J* is given by 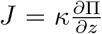 where *κ* denotes the permeability of the gel and Π denotes the osmotic pressure of the gel. Introducing d*z* = *ϕ*_0_d*Z*_0_*/ϕ* into the expression for *J*, we obtain 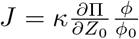. Using this result, the continuity equation for the aqueous phase of the gel is given by

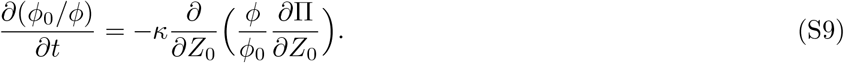

To close the equation, we use the Flory-Rehner theory to model the osmotic pressure of the hydrogel [1, 2], i.e., 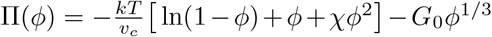, where the first term results from the free energy of mixing and the second term results from the elastic energy of stretching the polymers. Here, 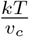 denotes the thermal free energy density, *χ* denotes the solvent-polymer interaction paremeter, and *G*_0_ is the shear modulus of the gel in the reference state. For agarose gels, we assume *χ* ≈ 1*/*2 and obtain 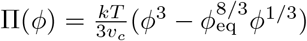 where 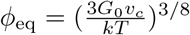 denotes the polymer volume fraction of a freely swollen gel. Finally, we set the reference state *ϕ*_0_ = *ϕ*_eq_ for simplicity.

##### Boundary conditions

In experiment, the bottom of the gel is connected with a water reservoir, and evaporation of water occurs at the top surface of the gel. To describe this experimental set up, we choose the following boundary conditions: Π = *C* = **ρ**_w_*g*Δ*H* at the bottom of the gel *Z*_0_ = 0, and *J* = *J*_ev_ at the top of the gel *Z*_0_ = *L*_0_. Here, *C* can be changed by tuning Δ*H* in experiment, and *J*_ev_ has been measured previously to be *J*_ev_ = 20 nm*/*s [3].

##### Nondimensionalization

We define dimensionless variables 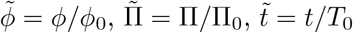, and 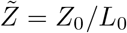, where Π_0_ = 1 kPa, *T*_0_ = 1 min, and *L*_0_ = 6 mm are the relevant scales for the experimental system. Upon non-dimensionalize Eq. (S9), we derive

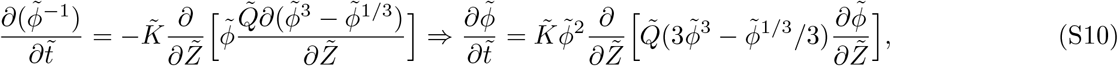

where 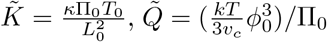, and we have used 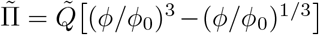. Similarly, the boundary condition for 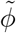 can be derived to be: 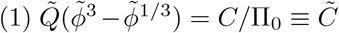 at 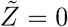, and 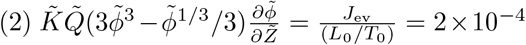 at 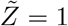.

##### Comparing with the experiment

In an experiment where the water reservoir is lowered by Δ*H* = 0.51 m, corresponding to a change in *C* from *C* = 0 kPa to *C* = 5 kPa (similar to the procedure in main Fig. 4), the system reaches the new steady state in 10–15 min and the gel shrinks by about 1%. To mimic this experimental procedure, we started 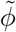at 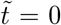 with the steady state profile 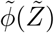 at 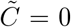, and simulated Eq. (S10) with 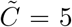 to track the system’s dynamics of approaching the new steady state. We define 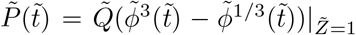 as the osmotic pressure difference across the gel surface, and we fit 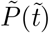 with 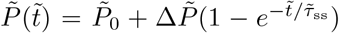 to obtain the relaxation time 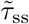. In addition, we also compute the steady-state gel thickness at 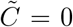 and 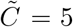 using the expression 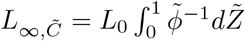. We use the relative gel thickness *α* = *L*_*∞*_ */L∞*,0 and the relaxation time 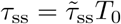 to compare the model to the experiment, yielding 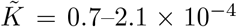 and 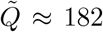 (Supplementary Fig. 6b and c). Finally, we use the fitted values of 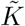 and 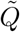 to estimate the steady state 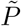 (or *p*) for varying values of 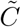 (or *C*), corresponding to different values of Δ*H* in experiments. Our analysis leads to a nonzero *P*_0_ ≈ 2 kPa, which is consistent with the value of *P*_0_ obtained from fitting the height profile. We also verify that *P* varies (approximately) linearly with the height difference Δ*H* between the water level of the reservoir and the gel surface (Supplementary Fig. 6d).

#### F Calculating the capillary force exerted on cells

We next consider the force exerted by the water meniscus on each cell, which we term the “capillary force”. We consider a water meniscus that is in mechanical equilibrium with the cells and has a height profile of *h*^*∗*^(*x, y*). For an infinitesimal surface element of the water meniscus that is in contact with a cell, we assume that the per-area force exerted on the cell by this element is ***f*** (*x, y*) ≡ −*f* (*x, y*)***N*** (*x, y*), where *f* denotes the magnitude of the force and 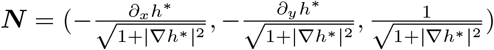 defines the local normal.

To compute the magnitude *f* (*x, y*), we consider a perturbation to the water meniscus ***u*** = *δ*(*x, y*)***N*** (*x, y*) and compute the free-energy change Δℱ associated with this perturbation. First, according to the virtual work principle, 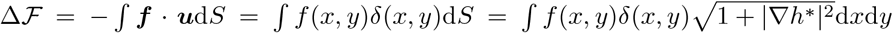, where *dS* denotes the infinitesimal area in 3D and the integral runs over the surface where the water meniscus is in contact with the upper surface of the cell. Additionally, the free-energy change can also be calculated directly using the free energy functional 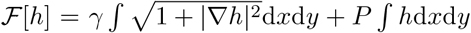. The perturbed water height profile is given by 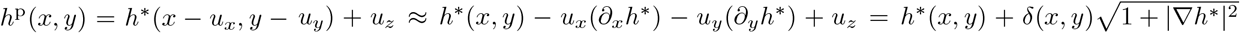 Thus,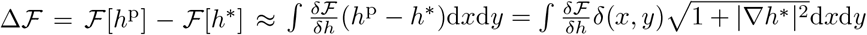 Comparing the two expression for Δℱ, we obtain

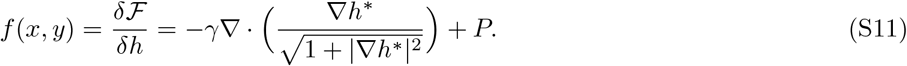

Introducing Eq. (S11) into the expression for ***f***, we obtain the capillary force acting on the cell per 3D surface area. If we define ***f***_cap_ to be the force per projected area d*x*d*y*, i.e., 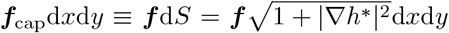, then ***f***cap is given by

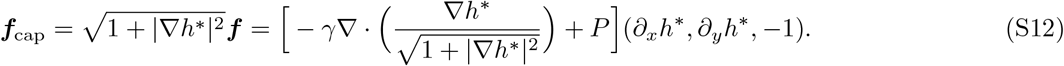

The *z* component of Eq. (S12) indicates that the water meniscus exerts a downward force on the cells. However, in the present work, we ignore the *z* component, assuming this is balanced by an upward force from the substrate, and focus on the horizontal capillary force and torque acting on each cell.

Thus, the total horizontal capillary force and torque exerted on cell *i* can be calculated as

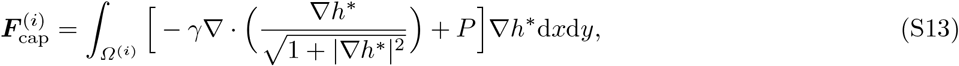

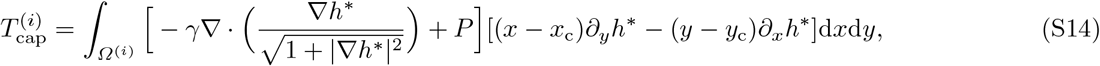

where *Ω*^(*i*)^ denotes the region where the water meniscus is in contact with cell *i*, i.e., 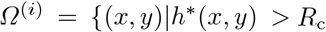 and 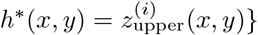 (see Eq. (S2)). In the simulations, we performed numerical integrals of Eqs. (S13) and (S14) to compute the capillary force and torque. As a cross-check, we also perturbed the position and orientation of the cell and calculated the force and torque from the change in the free energy of water, i.e.,

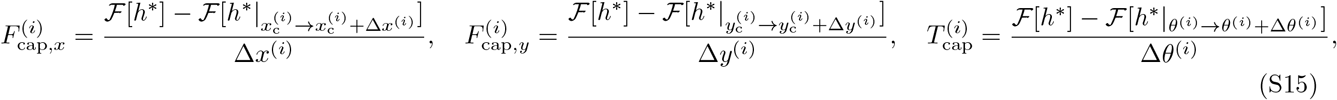

where *h*^*∗*^|_*C*_ denotes the new equilibrium height profile upon a change C in the position of the cell. The two methods yield the same computational results (Supplementary Fig. 7). For the rest of our simulations, we used the first integral-based method to calculate the capillary force and torque because it is more time efficient than the second method.

#### G Additional notes on the magnitude and range of capillary force

To gain more insights into the magnitude and range of capillary forces, we consider a group of *N* cells and calculate its per-cell capillary attraction ⟨*F*_cap_⟩_*N*_ by an infinitely large colony located a distance Δ away (see Supplementary Fig. 8). For a fixed number *N* of cells, we find that maximum ⟨*F*_cap_⟩_*N*_ occurs at Δ = 0, and that the magnitude of capillary attraction increases with the osmotic pressure of the gel *P* while the range of attraction decreases with *P* as ∼ *P*^*−*1*/*2^. This scaling relation can be understood by considering the simple case of a water meniscus around a 2D circle (Sec. I B). Equation (S5) shows that the half width of the water meniscus is *d*_wm_ = (*R*_c_ + *l*_*γ*_) sin *θ*^*∗*^ where 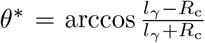. When *l*_*γ*_ ≫ *R*_c_, we derive that 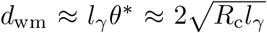, and thus *d*_wm_ ∼ *P*^*−*1*/*2^ since *l*_*γ*_ = *γ/P*. Next, we set Δ = 0 and look at how the maximum capillary attraction ⟨*F*_cap_⟩_*N*_ varies with the number of cells *N*, finding that it decreases with *N* as 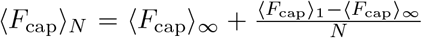. In particular, we find that ⟨*F*_cap_⟩_*∞*_ scales linearly with *P*. To gain some physical intuition for this relationship, we again turn to the simple case of the water meniscus around a circular cell. We consider a chain of closely packed circular cells extending from *x* = −∞ to *x* = 0. The water height profile follows Eq. (S5) for *x* ⩾; 0 and *h* ≈ 2*R*_c_ for *x <* 0 where cells are filled up. The capillary force acting on the cell at *x* = 0 can thus be calculated from Eq. (S13) to be 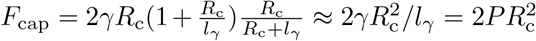 when *l*_*γ*_ ≫ *R*_c_.

## II. AGENT-BASED MODEL

### A Equations of motion

In this subsection, we summarize the different forces that we consider in the agent-based model, and list the equations of motion for each cell.

#### Cell shape

As noted in Sec. I A, we consider spherocylindrical cells oriented horizontally. For simplicity, we only consider the in-plane motion of the cells. Thus, unless otherwise specified, the vectors discussed below will be two dimensional in the *x* − *y* plane. We denote by 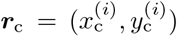 the center-of-mass position of cell *i*, and we define 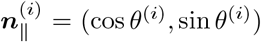 and 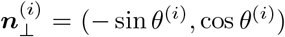 to denote, respectively, the unit vectors parallel to and perpendicular to the long axis of cell *i*.

#### Steric interaction between cells

For each pair of cells *i* and *j*, we compute the smallest distance *d*_*ij*_ between the centerlines of the two cell cylinders. Denote by 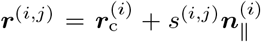 the coordinates of the point on cell *i*’s centerline that is the closest to cell *j*’s centerline. Thus, the distance *d*_ij_ can be calculated by 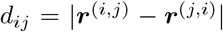. The magnitude of the repulsive force between cells *i* and *j* is modeled as 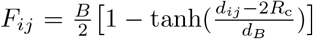, where *B* and *d*_*B*_ denote the magnitude and range of steric interaction, respectively. *F*_*ij*_ ≈ 0 if *d*_*ij*_ ≫ 2*R*_c_ + *d*_*B*_, i.e., when the cells are not in contact with each other. The total steric force and torque acting on cell *i* are given by

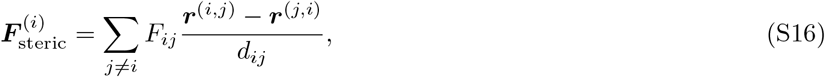

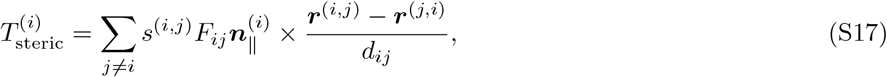

where the cross product of two-dimensional vectors ***f*** = (*f*_*x*_, *f*_*y*_) and ***g*** = (*g*_*x*_, *g*_*y*_) is defined by ***f*** × ***g*** = *f*_*x*_*g*_*y*_ − *f*_*y*_*g*_*x*_.

#### Friction between cells and substrate

Cell motion is opposed by friction from the substrate surface. We model the friction as a viscous drag that opposes the motion of each infinitesimal segment of the cell cylinder’s centerline. Denote by 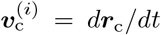 and 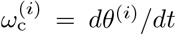 the translational and rotational velocity of cell *i*. The velocity of the point 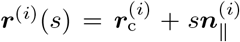 on cell *i* is 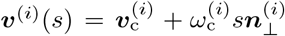. Assuming anisotropic friction coefficients in the directions parallel to and perpendicular to the long axis of cell *i*, we derive the infinitesimal frictional force *d****F***_fric_ acting on ***r***^(*i*)^(*s*) to be given by 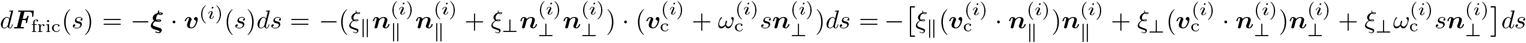. Thus, the total frictional force and torque acting on cell *i* are given by

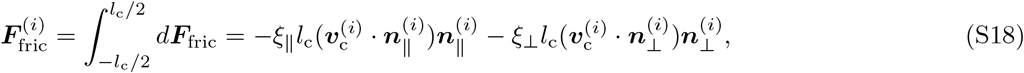

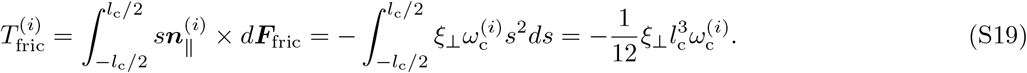

#### Self-propelling force of each cell

The self-propelling force is assumed to be along the long axis of each cell, i.e.,

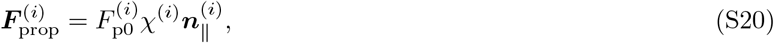

where *F*_p0_ denotes the magnitude of self propulsion and *χ* = ±1 follows a dichotomous Poisson process with rate 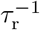. The cells do not generate self-propelling torques, i.e., *T*_prop_ = 0.

#### Capillary force acting on the cells

The capillary force and torque exerted by the water meniscus are discussed in Sec. I F and are given by Eqs. (S13) and (S14).

#### Equations of motion

Taken together, the balance of the above forces determine the equations of motion for the modeled cells,

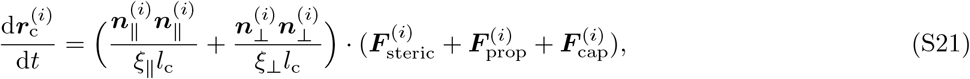

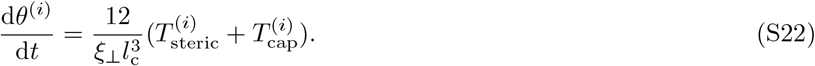

Note that the dichotomous noise for reversal is the only source of noise in the system.

#### Growth of non-motile cells

To study how capillary forces influences the development of growing colonies, we follow previous work [4] and model bacterial growth as elongation of modeled spherocylinderical cells, whose projected area is given by 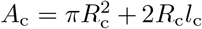. Denoting by *α*_g_ the areal growth rate, and introducing the expression for *A*_c_ into d*A*_c_*/*d*t* = *α*_g_*S*, we obtain

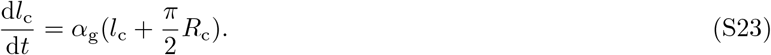

Each modeled cell has an initial length of *l*_0_, elongates according to Eq. (S23) until it reaches a final length of *l*m = 2*l* + 2*R*, and divides into two daughter cells. Cell division is modeled by replacing the mother cell at 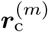 with two spherocylinders of length *l*_0_ at positions 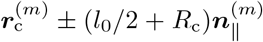 with the same orientation 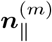 as the mother cell.

### B Simulations and parameters

The cell positions and the equilibrium water height profile are updated iteratively. The equations of cell motion are simulated using a forward Euler method with an adaptive time increment such that for every time step the maximum displacement of any cell segment does not exceed 0.1*R*_c_. The simulation parameters are chosen or determined as follows:

#### Capillary force

As discussed in Sec. I D, the surface tension *γ* is measured to be *γ* = 66 mN*/*m. The osmotic pressure *P* is varied between *P* = 2.8 kPa and *P* = 8.8 kPa.

#### Steric interaction

To capture the steric interactions between cells, we choose *B* = 8*γR*_c_, which is much larger than typical capillary force and self-propelling force, to ensure that the modeled cells cannot penetrate into each other. We set *d*_*B*_ = 0.1*R*_c_ to ensure that the interaction is very short-ranged.

#### Friction coefficients

We estimated the coefficients of the anisotropic friction between the cells and the substrate from an experiment in which two nearby non-motile cells/cell clusters are pulled together by capillary attraction (see main Fig. 2). We fit the modeled merger dynamics to the experiment. The fitting error was calculated by comparing the center-to-center distance 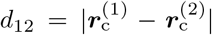 and the angle *ϕ*_12_ = *θ*^(1)^ − *θ*^(2)^ of the two cell clusters between the model and the experiment. For each time point *t*, the fitting error is given by the root-mean-square-deviation 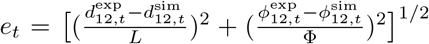 where the superscripts exp and sim denote the quantities in experiment and simulation, respectively, and we set *L* = 2.5 µm and Φ = 1 rad to be the characteristic scales for length and angle. The total fitting error was calculated as ∑_*t*_*e*_*t*_. Finally, we varied the friction coefficient *ξ*_*⊥*_ and friction anisotropy 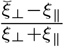 to determine the optimal fitting parameters that minimize ∑_*t*_ *e*_*t*_ (Supplementary Fig. 9), which yielded *ξ*_*∥*_ = 80 Pa · min and 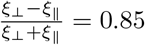.

#### Phase diagrams

To explore the collective dynamics in the model, we varied three dimensionless model parameters: a normalized reversal period 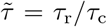 where 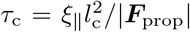 is the time it takes for a non-reversing free-running cell to travel its cell body length, a normalized self-propulsion strength 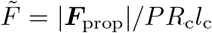 where *PR*_c_*l*_c_ is the scale of the longitudinal capillary force acting on isolated cells, and a normalized capillary length 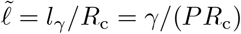.

#### Cell growth

The areal growth rate *α*_g_ of bacteria was determined by tracking the total area covered by the cells over time (Supplementary Fig. 13).

### C A toy model for the splitting of cell groups

In the presence of a water meniscus, a group of cells can split into two subgroups if they propel sufficiently strong in opposite directions to overcome the opposing capillary force. We wondered how this capillary attraction depends on the partition of the subgroups, which describes the number of cells in each subgroup. To explore this question, we consider two subgroups with *N*_1_ and *N*_2_ cells that are aligned in the same direction but propel in opposite directions (Supplementary Fig. 10). Since the equilibrium water meniscus experiences zero net force exerted by the cells, the total capillary force exerted on the two groups must have the same magnitude 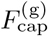 but opposite directions. Assuming that each cell has the same self-propelling force *F*_p0_ and the cells in the same subgroup move at the same speeds *v*_1_ and *v*_2_, we derive the force-balance conditions for the two groups

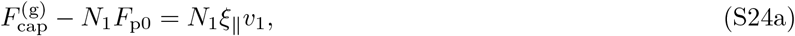

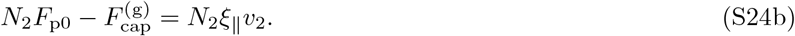

Splitting occurs when *v*_2_ *> v*_1_, i.e.,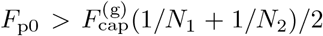. As shown in Supplementary Fig. 10, for a fixed number of total cells *N*_*1*_ + *N*_*2*_, the effective per-cell capillary attraction 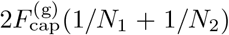 is the smallest when *N*_1_ ≈ *N*_2_, suggesting that symmetric splitting of cell groups is favored. In addition, for a fixed number *N*_2_, the effective per-cell capillary attraction on cells in group 1 decreases with increasing *N*_1_, as noted in Sec. I G.

### D Quantifying the elongation and splitting of cell groups

As illustrated in Supplementary Fig. 10 and discussed in Sec. II C, cells within a group can propel in the opposite directions, which creates a force dipole that tends to either elongate the cell group when the capillary force is strong or split apart the group when the capillary force is small.

To quantify the elongation of the cell group, we first consider a group of *N* = 10 cells with relatively small values of 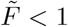 The contour of the cell group is determined by the contour line of the water height profile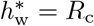. We follow previous literature [5] to analyze the deformation modes of the contour. Briefly, we first compute the center of all points on the contour, defined as their average position weighted by the local curvilinear distance, and calculate their polar coordinates (*r*_*i*_, *θ*_*i*_) with respect to the center. Subsequently, we decompose the contour into Fourier modes, i.e., 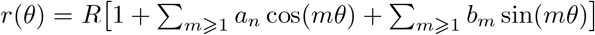 where 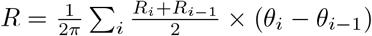 denotes the zero mode component, and compute 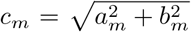 to characterize the mode-*m* deformation of the contour (Supplementary Fig. 11). Mode *m* = 1 corresponds to the translation mode of the contour and thus *a*_1_ = *b*_1_ ≡ 0 by our definition. In our simulations, the dominant deformation mode of cell groups is mode *m* = 2, which corresponds to the elliptical elongation of the contour (Supplementary Fig. 11a). To quantify the shape fluctuation of the cell groups, we compute the fluctuation of mode *m*=2, i.e., 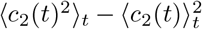, finding that it increases with increasing reversal time 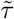 of the cells and increasing self-propelling force 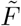 (Supplementary Fig. 11b).

To quantify the splitting of the cell group, we consider a group of *N* = 10 cells with 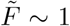. We use the criterion 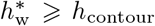 to determine the cell regions G_cell_ and track the number *n*_cc_ of connected components in G_cell_. The splitting events are identified by changes in *n*_cc_ from *n*_cc_ = 1 to *n*_cc_ *>* 1. The splitting rate of cell groups can be determined from the inverse of the average waiting times in the *n*_cc_ = 1 state. As shown in Supplementary Fig. 11c, our simulations show that the splitting rate also increases with increasing reversal time 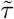 of the cells and increasing self-propelling force 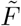.

### E Capillary forces facilitate merger of growing colonies

Since water menisci should generally exist around bacterial cells grown on hydrated substrates, we wondered how capillary forces might affect the development of growing but non-motile colonies. To explore this question, we imaged *E. coli* colonies, which are known to be poorly motile on hard agar [6], and found that two adjacent colonies elongate and expand preferentially toward each other (Supplementary Fig. 14). We hypothesized that the elongation of growing colonies is due to the asymmetric capillary force acting on them. Specifically, the water meniscus between the two colonies has a smaller slope than that in other regions, and thus the in-plane capillary force is smaller in the middle, allowing the colonies to expand more easily toward each other. To test this hypothesis, we also simulated colony growth in the presence of a water meniscus, and our simulation results are in quantitative agreement with the experiment. To further verify our hypothesis, we simulated colony growth in the absence of capillary forces, finding that in this case the adjacent colonies stay roughly circular and do not expand toward each other (Supplementary Fig. 14).

## III. CLASSIFYING AND QUANTIFYING DIFFERENT PHASES OF COLLECTIVE CELL DYNAMICS

### A Phase classification

Simulations of the agent-based model described in Sec. II yielded a range of collective dynamics when the normalized parameters of water and cell motility are varied. Qualitatively, the simulated dynamics can be categorized into four phases:

1. a “gas” phase in which cells are spatially distributed rather than packed together by the capillary force,
2. a “polar clusters” phase in which cells are clustered into small groups that move persistently over long distances and constantly undergo merging and splitting events,
3. a “streams” phase in which cells self-organize into a quasi-one-dimensional structure, and
4. a “droplets” phase in which cells are packed together and trapped in cell groups that do not move persistently.

To quantify the collective dynamics of *N* cells, we compute the equal-time correlation function of cell density **ρ**

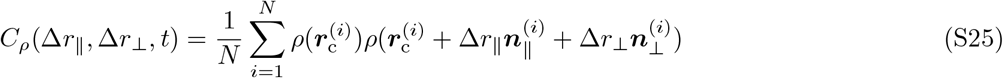

and normalize *C*_**ρ**_ by its maximum *C*_**ρ**_(0, 0, *t*) to obtain 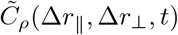. The average and standard deviation of 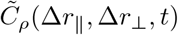 with respect to time, denoted by 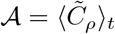 and 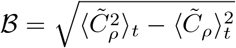 characterize the morphological and dynamical features of 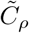, respectively (main Fig. 3 and Supplementary Fig. 15a).

To classify the simulation results with different parameters in an unbiased way, we develop a measure of distance between any pair of simulated dynamics, and use a standard clustering method to group different simulations into four clusters (Supplementary Fig. 15b). We follow previous computer-vision literature [7, 8] and measure the pairwise “distance” between the simulated dynamics using the distances of 𝒜 and B in the Sobolev space. For arbitrary matrices X_*i*_ and X_*j*_, the Sobolev norm of order 1*/*2 is given by

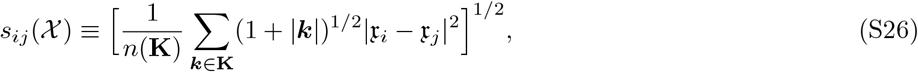

where 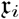 and 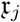 denote, respectively, the discrete Fourier transforms of 𝒳_*i*_ and 𝒳_*j*_ and *n*(**K**) denotes the number of discrete frequency vectors in the frequency domain **K**. We define the distance D_*ij*_ between simulations i and j as 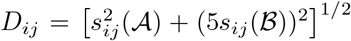 where the factor 5 is chosen such that the terms containing s_*ij*_ (𝒜) and 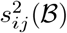 have similar magnitudes. The clustering is performed using the WPGMA (Weighted Pair Group Method with Arithmetic Mean) method and implemented in MATLAB. Finally, we handpick one representative simulation for each phase of collective dynamics to verify and annotate the clustering results.

### B Phase quantification

As shown in main Fig. 3, for a fixed self-propelling force, the collective cell dynamics transition from the gas phase when the cell-cell capillary attraction is low, to the streams and polar clusters phases when the cell-cell capillary attraction is intermediate, and to the droplets phase when the cell-cell capillary attraction is high. The effect of the capillary attraction on the collective cell dynamics manifests itself in a trade-off between cell mobility and cell-cell adjacency. To quantitatively describe this trade-off, we compute the instantaneous cell speeds, normalized by the average free-running speed |**F**_prop_|*/*(*ξ*_∥_*l*_c_), and the cell-cell distances^2^, normalized by the average cell body length *l*_c_, for the different phases. Our simulations show that both the cell speed and the cell-cell distance decreases with increasing magnitude of capillary attraction (Supplementary Fig. 17a,b). The streams and polar clusters phases show similar distribution of cell-cell distances, but their collective dynamics are different – cells moving in quasi-1D streams frequently slip past each other while cells moving in small polar clusters stay together for a relatively long time. To capture this difference, we compute the cell-cell adjacency matrix *A*_*ij*_(*t*), where *A*_*ij*_ is 1 if the distance between cells *i* and *j* are smaller than 4*R*_c_ and 0 otherwise. The average autocorrelation function of this matrix *C*_A_(Δ*t*) ≡ ∑*i*≠ *j* ⟨*A*_*ij*_(*t*)*A*_*ij*_(*t*+Δ*t*)⟩_*t*_/ *i j* ⟨*A*_*ij*_(*t*)*A*_*ij*_(*t*)⟩_*t*_ describes the probability that a pair of adjacent cells remain adjacent after time Δ*t*. The life time of adjacent cell pairs is the longest in the droplets phase, second longest in the polar clusters phase, third longest in the streams phase, and the shortest in the gas phase (Supplementary Fig. 17c). In the droplets phase where the self-propelling force of each cell is much smaller than the capillary force, entire cell groups undergo collective translational motion. Intuitively, over time scales longer than the mean reversal time *τ*_r_ of the cell, such group motion is diffusive, and the group diffusivity *D*_g_ should decrease with increasing number *N* of cells in the group because the overall propelling force of the group 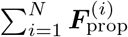 scales with 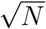 while the overall friction of the group scales (approximately) with *N*. To verify our intuition, we simulate the dynamics of cell groups with varying cell numbers *N* in the droplet phase. We use the same thresholding method as in Sec. II D to determine the cell regions and track the center-of-mass trajectories ***r***_g_(*t*) of the cell groups. The mean squared displacement (MSD) is subsequently computed as MSD(Δ*t*) = ⟨|***r***_g_(*t* +Δ*t*) − ***r***_g_(*t*)|2⟩_*t*_. We obtain the group diffusivity *D*_g_ by fitting a linear function *D*_g_Δ*t* to the MSD with a lag time Δ*t >* 3*τ*_r_. Our analyses show that, when friction is isotropic, i.e., *ξ*_⊥_ = *ξ*_∥_, both the mean squared speed ⟨*V*_g_⟩2 and the diffusivity *D*_g_ of the group scale inversely with *N* and they are related by *D*_g_ = ⟨*V*_g_⟩2*τ*_r_ (Supplementary Fig. 18). For anisotropic friction *ξ*_⊥_ *> ξ*_∥_, the relationship *D*_g_ = ⟨*V*_g_⟩2*τ*_r_ remains true. In this case, *D*_g_ and ⟨*V*_g_⟩2 decrease with *N* more rapidly than *N*^*−*1^ (Supplementary Fig. 18) because cells can align in different orientations in groups with large *N* and get pushed sideways, thus increasing the overall friction and decreasing the group diffusivity and mobility compared to the case of isotropic friction.

## IV. CONTINUUM MODEL

### A Phase-field formulation

To describe the large-scale organization of cell groups in the presence of water, we study a minimal phase-field model. Briefly, we consider groups of cells that self-propel in *M* pairs of opposite directions 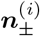 with densities 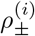 and velocities 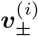. Since we are interested in the large-scale morphology of *M. xanthus* cell groups, we introduce a phase field *ϕ* that tracks the region of cells. *ϕ* = 1 denotes regions filled with cells and *ϕ* = 0 denotes cell-free voids. Thus, the mass conservation equation can be expressed as

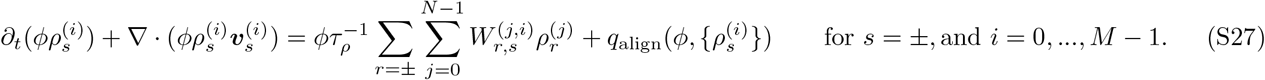

Here, *τ*_**ρ**_ denotes the rate of changing propulsion directions and 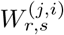 denotes the transition probability matrix. The alignment term *q*_align_ describes the dynamics of cell alignment due to cell-cell and cell-meniscus interactions. To model the nematic cell-cell alignment, we consider the local nematic tensor order parameter 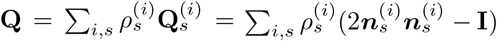, and an associated free energy,

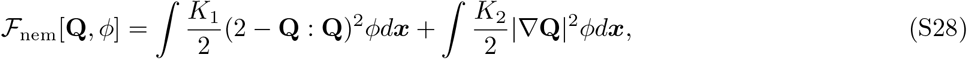

where the first term favors local nematic ordering with a coefficient *K*_1_, and the second term penalizes gradient of nematic order parameter with a coefficient *K*_2_. To capture the alignment with the boundary of water meniscus, we consider an additional surface free-energy term with nematic symmetry 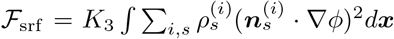. By introducing the expression for **Q** into Eq. (S28) and taking the functional derivative with respect to 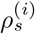, we derive that

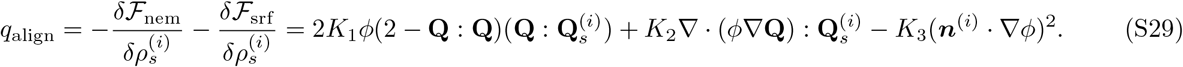

Using the phase field *ϕ*, we derive the force-balance condition

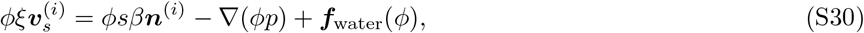

where the effect of the water meniscus is described by ***f***_water_ and is nonzero only at the boundary of cell regions. In our agent-based model, we found that the water meniscus exerts an in-plane pressure on the periphery of a cell group and the pressure is proportional to the curvature of the boundary, similar to the Laplacian pressure due to surface tension (Supplementary Fig. 21). Thus, in the continuum model, we treat the cell population as 2D droplets, and approximate the effect of water menisci by an equivalent liquid surface tension *σ*, which will be described below. Finally, as described in previous works [9–11], the time evolution of *ϕ* follows

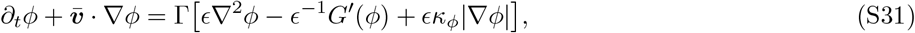

where the phase field *ϕ* is advected by the mean velocity 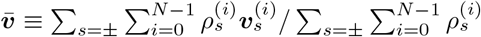. We choose the standard double-well potential *G*(*ϕ*) = 18*ϕ*^2^(1 − *ϕ*^2^) such that the bulk regions have either *ϕ* = 1 or *ϕ* = 0. Here, Γ denotes the relaxation rate of the phase field, *ϵ* denotes the width of the boundary layer, and 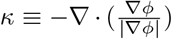 denotes the local curvature. Using the phase field *ϕ*, we express the surface force ***f***_water_ as ***f***_water_ = *σ*(*ϵ*∇2*ϕ* − *G*^*′*^(*ϕ*)*/ϵ*)∇*ϕ* [11].

#### Choice of parameters

We simulate the above equations on a square domain [−10, 10] × [−10, 10]. We set the phase-field parameters *ϵ* = 0.5 and Γ = 1. We set the scale of time, force, and density by setting ξ = 1, *β* = 1, and 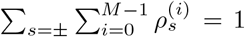, respectively. For simplicity, we study the case *M* = 3. We set the propensity of reversing self-propelling directions to be the highest 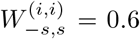, and we set the propensity of changing to other directions to be the same 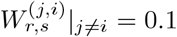. To reproduce the polar clusters, streams, and droplets observed in our agent-based simulations, we set the alignment parameters to be *K*_1_ = 1, *K*_2_ = 0.01, and *K*_3_ = 1. The remaining parameters *σ* and *τ*_**ρ**_ are varied to explore how the strength of capillary force (relative to the self-propelling force *β* = 1) and the cell reversal frequency influence the large-scale dynamics.

### B Numerical scheme

The numerical scheme for solving Eqs. (S27-S31) was described previously in detail [11]. Briefly, we employed an isotropic finite-difference method to compute spatial derivatives on a discretized grid of 256 × 256 points. Equations (S27) and (S31) were solved using an Euler forward scheme with a fixed time increment Δ*t* = 2 × 10^*−*3^. To ensure numerical convergence, a small diffusion term were introduced to Eq. (S27).

To solve the force-balance equation Eq. (S30), we separate the velocities 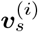 into two terms 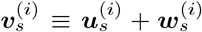, where 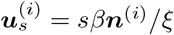. Introducing this expression into Eq. (S30) and the incompressibility condition, we obtain

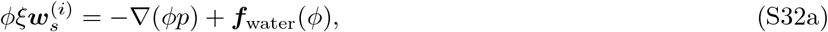

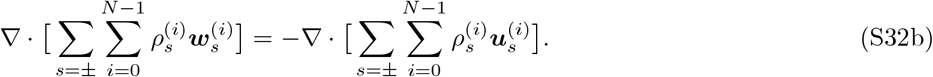

Since Eq. (S32a) is independent of *s* and *i*, we look for solutions 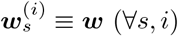. Thus, ***w*** follows

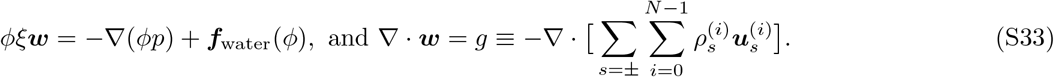

Note that in deriving Eq. (S33), we have used the normalization 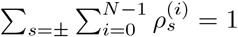

Equation (S33) was solved iteratively using a semi-implicit Fourier-spectral method. In particular, we formally rewrite Eq. (S33) as

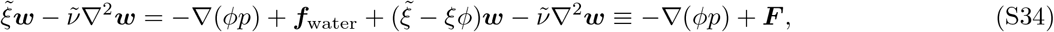

where we have introduced two positive constants 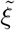 and 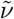 to stabilize the numerical scheme, and we have grouped all the non-pressure terms into ***F***. Taking the divergence of Eq. (S34) and using ∇· ***w*** = *g*, we obtain 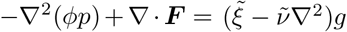. We denote by [*f* ]_***k***_ = ∫ *f* (***x***)*e*^*−i****k****·****x***^*d*^2^***x*** the Fourier transform of an arbitrary function *f* (***x***). The Fourier transform of the simplified equation leads to 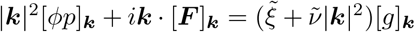. Introducing this relation into the Fourier transform of Eq. (S34), we obtain that 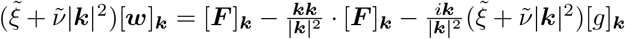. We solve for ***w*** iteratively using the following recursion relation

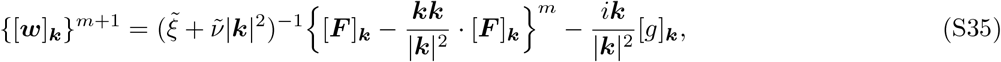

where {}^m^ denotes the expression evaluated at the *m*^th^ iteration step (for a particular time step). We obtain {***w***}^m+1^ from the inverse Fourier transform of {[***w***]_***k***_}^m+1^, which is then used to update {***F*** }^m+1^ in the next iteration step. The iteration ended when max |{***w***}^m+1^ − {***w***}^m^| *<* 10^*−*3^ max{***w***}^m+1^. Finally, the pressure can be obtained by applying inverse Fourier transform to 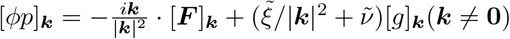.

### C Stream index

Simulations of the continuum model yielded two distinct morphologies of the cell populations: one consisting of stream-like aggregates of cells, and one consisting of droplet-like aggregates of cells. Thus, we sought to develop a metric to quantify the “stream-ness” of the overall morphology, which we term the “stream index” *S*. For simplicity, we define the cell regions as ℛ ≡ {(*x, y*)|*ϕ*(*x, y*) *>* 0.5} using the phase field *ϕ*. To highlight the topological features of ℛ, we extract the skeletons *sk*(ℛ_*i*_) of all the connected components ℛ_*i*_ in ℛ (Supplementary Fig. 20). For an arbitrary shape ℰ, the topological skeleton *sk*(ℰ) is defined as the ridges (i.e., curvature singularities) of the distance transformation of ℰ, which calculates the minimal distance of any point *E* ∈ ℰ to the shape boundary *∂*ℰ. Note that the length of skeleton |*sk*(ℰ)| is zero for a circular shape, and |*sk*(ℰ)| *>* 0 for an elongated shape. Thus, a dimensionless quantity 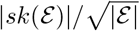 can be used to quantify the “stream-ness” of shape ℰ, where |ℰ| denotes the area of ℰ. Finally, for a structure ℛ that contains *n* disconnected shapes ℛ_*i*_, we define the stream index *S* as 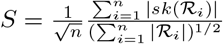 where the factor 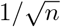 is introduced such that *n* identical shapes have the same *S* as each individual shape. Our analyses show that the *S* index defined as such is able to clearly distinguish between stream-like (*S* ⩾ 1) and droplet-like (*S <* 1) morphologies (Supplementary Fig. 20).

### a) CELL CULTURE AND STRAINS

#### Cell Culture and Growth

All strains used in this study are listed in Supplementary Table 1.

*M. xanthus* was grown in 1% CTT media consisting of 10 g/L Peptone (Ovia), 1% v/v Tris-HCl buffer, and 0.1% v/v 1 M KH2PO4 manually adjusted to pH=7.6. All cells were struck from frozen stocks onto 1.5% v/v agar/CTT plates. The day before each experiment, single colonies were picked from plates and incubated in 20 mL of CTT, shaking at 32^*°*^C overnight. Cells were collected at an optical density, OD600 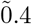. We then concentrated or diluted cells to the desired cell density by centrifugation at 8000rpm for 5 minutes in a tabletop centrifuge (Eppendorf 5415D) and subsequently resuspending the pellet in fresh CTT of the desired volume.

*F. johnsoniae* cells were grown in 1% CYE media, first from frozen stocks on 1.5% agarose/CYE plates. Single colonies were selected and grown overnight in liquid CYE, shaking, at 26^*°*^C. Prior to each experiment, overnight cultures were back-diluted 1:100 in fresh motility media and grown shaking for 2 hours before the cells were spun and resubmerged in fresh motility media. CYE media consists of 10 g/L Peptone (Ovia), 5 g/L yeast extract (BD), 8 mM MgSO_4_, and 10 mM Tris (pH=7.5). Motility media consists of 1 part CYE without MgSO_4_ (see above) and 2 parts water.

*E. coli* was grown first as single colonies on 1.5% v/v agarose/LB plates. Single colonies were selected and grown overnight in liquid LB, shaking at 37^*°*^C. Prior to each experiment, overnight cultures were back-diluted 1:100 in fresh LB and grown shaking at 37^*°*^C for 2 hours before the cells were transferred to an agarose pad for imaging.

## VI. EXPERIMENTAL DEVICE AND SETUP

### A Microscopy

All images presented were obtained on a Keyence VK-X1000 confocal microscope. A custom MatLab program was used for microscope control and image acquisition. At each time, the microscope captures an image of (1) the reflectance of the sample at each pixel and (2) the height of each pixel relative to the image’s minimum. This is done by scanning through a finite axial range and determining, at each pixel, the optimal reflectance and its corresponding height using a proprietary algorithm. Height images were corrected for surface tilt, see VII A. For *F. johnsoniae* and *E. coli*, the cells were not sufficiently discernable in the reflectance images to be labeled and segmented by our neural network. Therefore, we also captured a color image of the cells at each time. This image by a similar scanning process as used to acquire the reflectance and height images, except the light is passed to a CMOS camera chip instead of a photodiode. A final image is constructed by determining, for each pixel, which point in the stack of collected images is the most in focus (as determined by an undisclosed algorithm).

All timelapses were captured at a rate of 1-3 minutes per frame. At each point in time, the microscope was automatically refocused and the appropriate scan range determined by either determining the range used in the previous frame and adding a small “buffer” to ensure a complete scan or by an automatic range-determination algorithm internal to the microscope. Lateral sample drift was corrected by post-processing, see Section VII B for details.

### B Device Description

We designed a custom device to control the osmotic pressure across the hydrogel substrate. The device works by applying a hydrostatic head to the bottom of the gel. This hydrostatic head then serves as an additional pressure on top of the typical osmotic pressure (gradient) that exists in any hydrogel exposed to air. To do this, a reservoir of liquid media (or water) is connected directly to the gel. By moving the reservoir up or down relative to the gel, a hydrostatic head is applied to the gel. This hydrostatic pressure acts in addition to the usual osmotic pressure at the gel/air interface and thus directly modifies the water availability, and thus the cell wetting, at the surface. See Supplementary Figs. 3 and 4.

All custom parts were made by laser cutting acrylic sheets of varying thickness (specified below, per part). Acrylic sheets were purchased from McMaster-Carr and laser cut on a Universal Laser Systems VLC 3.60 laser cutter.

The base of the device consists of an elevated plate with a hole in the middle (Supplementary Fig. 3a). This is cut from a single piece of 0.5”-thick acrylic. This middle hole is cut such that it does not penetrate through the plate, but instead fits around and supports a male Luer lock (McMaster Cat. #) that will penetrate completely through the plate. The base is elevated by four 0.5^3^” legs (0.5^2^” squares cut from 0.5”-thick acrylic) that are glued to the bottom of the base with acrylic glue (SciGrip IPS Weld-On # 3). Halfway along each outer edge, a through hole is cut. Heat-set inserts (McMaster Cat. # 94459A320) for 8-32 screws were then fit into these through holes with a soldering iron.

Placed atop the base is a support plate, consisting of a 0.25”-thick circular piece of acrylic with a through-hole cut such that a male Luer lock can be press-fit into it. This plate should sit flush to the base when constructed properly. A spacer ring that outlines this plate is then placed on top of it. The thickness of this spacer ring will determine the thickness of the gel. For all experiments in this work, a 3 mm thick spacer was used. Finally, a thin plate (cut from 3 mm-thick acrylic) is placed on top. Crucially, this plate contains a 15mm opening in the middle, as well as four holes near its outer edge that are to be lined up with the threaded holes in the base. The middle, 15 mm-diameter opening in the middle of the plate is left exposed to air and is where the sample will be deposited and imaged. As such, its area is a key determinant of how rapidly the gel loses water due to evaporation. We found that 15 mm-diameter was wide enough to comfortably fit the nose of the objective close enough to sample such that it could be imaged while not being so wide that the the additional hydrostatic head from the reservoir would be insufficient to keep the gel hydrated over the course of long (multi-hour) times.

The device is assembled by stacking the components, as previously described (Supplementary Fig. 3). Finally, the pieces are fastened together by pushing four 8-32 screws through the top plate through holes and into the inserts in the base. Relevant dimensions for the device are shown in Supplementary Fig. 3c,d.

After the pieces are fastened together, a syringe filled with media and connected to a fluid line (Masterkleer PVC plastic tubing, McMaster Cat. # 5233K51) is connected to the Luer lock on the plate and filled completely such that no air bubbles are in the line. Molten agarose gel is then poured into the top opening until it is completely filled. The gel is then allowed to cool and thus, solidify. We note that while a small amount of agarose does go into the line, this can be minimized (¡1mm depth into the gel) by pouring the gel at a temperature just above it’s melting temperature. Once the gel has solidified, cells are pipetted onto the gel, the spot allowed to dry, and the device is transferred to the microscope. Here, the syringe is disconnected from the fluid line and the line is reconnected to the reservoir. As a reservoir, we used plungerless 10 mL syringes with female Luer locks screwed onto the end. The reservoir was mounted to a rail connector that then ran along a piece of optical dovetail rail mounted perpendicular to the optical table. After setting the reservoir height, we allowed the system to stabilize for 15 minutes before starting to image.

### C Measurement of Liquid Surface Tension

Surface tension for all media used was measured by the pendant drop tensiometry using a custom-built tensiometer. The tensiometer consists of a Basler Ace acA3088-57uc camera equipped with a Computar MLH-10X close-up manual zoom lens focused on a backlit, blunt-end dispensing syringe. Pendant drops were manually loaded into the syringe and imaged once before being pushed out of the syringe and replaced with a new drop. Each trial consists of a single, imaged drop. Images were processed using OpenDrop [12].

Measured surface tensions for all media used in this study is given in Supplementary Table 3.

## VII. IMAGE AND DATA ANALYSIS

All analysis was done in Python 3.9 [13] using the NumPy [14], SciPy libraries [15], and scikit-image [16] libraries. Plots were generated using Matplotlib [17].

### A Image Height Correction

Height maps for all analyzed images were corrected for surface tilt. To do this, we first automatically identified pixels corresponding to the underlying substrate and then fit them to a first or second order polynomial.

To identify the substrate, the reflectance image was converted to an 8-bit image and a local entropy filter was applied using a diameter=30 pixel disk as a structuring element (Fig. 26c). The resultant image was then flattened and the values histogrammed to generate a distribution of entropy values (Fig. 26d). Because (groups of) cells exhibit greater local entropy due to their exhibiting an edge around the surface, the distribution of entropy values was typically bimodal with one mode representing the typical (lower) entropy value of the surface versus the other being higher and typically corresponding to the cells. We fit the resulting distribution to a 2 component Gaussian mixture model and determined the threshold for calling a pixel part of the surface by multiplying the mean of the smaller component by 1.1.

Heights for the list of resulting (x, y) pixel coordinates that are considered part of the underlying substrate were then fit to a three-dimensional surface. We found that for all experiments, either a 1^st^ (*h*(*x, y*) = *ax* + *by* + *c*) or 2^nd^ (*h*(*x, y*) = *ax*^2^ + *by*^2^ + *cxy* + *d* were sufficient models of the underlying surface (*R*^2^ *>* 0.9, Fig. 26e). This fitted function was then applied to all (x,y) coordinates to generate a representation of the substrate and subtracted off from the original height map to generate a “corrected” map. This corrected map was used for all analyses.

### B Timelapse Drift Correction

To account for global sample drift over the course of long time series, images were aligned into a global coordinate system for each time series. Subsequent analyses were conducted in this coordinate system. To do this, we first calculated a relative shift between all pairs of adjacent frames by calculating the phase difference between the Fourier transforms of these frames and finding the maximum value of the resulting cross power spectrum. See [18] for details. These shifts are then cumulatively summed to yield a “global” shift for each frame, assuming that the first frame is unshifted. The largest global shift also yields a sufficient amount of padding that is applied to all images such that none of them are truncated upon shifting. Shifts are applied to each image by Fourier transforming each image, multiplying the result by a the shift operator 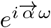where 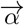 is the desired shift and *ω* is spatial frequency, and then inverse Fourier transforming the result.

### C Convolutional neural network-based cell segmentation

#### 1 Network Architecture

We use a standard U-net architecture [19]. The contracting and expanding paths of the net consist of 5 convolutional blocks, each corresponding to a single spatial scale. Each convolutional block consists of 2 convolutional layers with ReLU activations, 3×3 pixel kernels, and implicit padding such that the output maintains the shape of the input. Spatial downsampling in the contracting path is done by 2-dimensional maximum pooling such that the spatial dimensions of the input are reduced by a factor of 2 after each pooling operation. In the expanding path, we use bilinear interpolation to expand the spatial dimensions by a factor of two between spatial scales. All network weights are initialized with Kaiming initialization [20].

For *M. xanthus*, the network takes as input both the laser intensity image and its corresponding height map, z-scored with respect to the mean and standard deviation of the training set data. For *F. johnsoniae* and *E. coli*, the RGB image collected at each time point is concatenated onto the two-channel laser intensity image and height maps. For tracking single *M. xanthus* cells in groups, an additional dataset was generated similar to that for *F. johnsoniae* and *E. coli* wherein an RGB image of the cells was also collected. This 5 channel image is similarly z-scored with respect to the mean and standard deviation of the training set data. The network outputs a multi-channel confidence map that predicts, independently, the probability that each pixel does/not belong to either a cell or border pixel. All code was written in PyTorch [21].

#### 2 Training set generation

A ground truth dataset of cell masks was generated by manual annotation of 191 images for *M. xanthus*. For *F. johnsoniae*, a ground truth dataset of 46 images was used. For *E. coli*, 19 images were included in the ground truth dataset. For the dataset of *M. xanthus* cells with included RGB images, 52*images* were included. Cell masks were manually annotated using either a custom Matlab GUI, or by manually drawing the masks on an iPad using the ibis Paint app. First, the GUI (or iPad) was used to create an initial dataset consisting of matched pairs of cell masks and their corresponding laser intensity and height images. This dataset was then used to train a preliminary version of the cell segmentation network. The trained network was then used to predict segmentations on new images, which were then manually corrected using a custom Matlab GUI before being added to the ground truth dataset. This ”human in the loop” procedure was repeated until satisfactory network performance was reached.

A “cell border” class was added to the ground truth cell masks following the procedure in [22]. Briefly, cell masks were dilated with a 2 pixel-radius disk and all pixels in the resulting mask that are not labelled as “cell” in the ground truth are marked as “border.” We note that in our labeling, cells by definition must have at least a 1 pixel gap and thus a border region always exists between 2 cells.

During training, this ground truth dataset was split 80/20 into training and validation sets.

#### 3 Training details

Segmentation networks were trained using the Adam optimizer with initial learning rate 0.0005 and moment estimate decay rates, *β*_1_, *β*_2_ set to 0.9, 0.999, respectively. Over the course of training, the learning rate was reduced by a factor of 0.25 after the validation loss had reached a plateau for 100 epochs. To prevent overfitting, the network was trained until the validation loss reached a plateau for 250 epochs (early stopping). We use dice-weighted cross entropy (Eqn. S36) as a loss function where the cross entropy term was annealed (*τ* in the Eqn. S36) over the course of training such that the final network was trained solely on the dice loss.

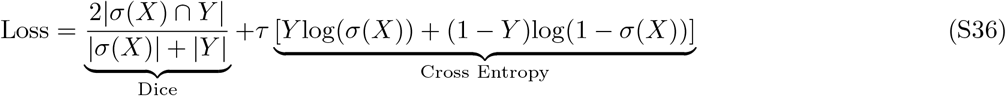

To do this, we used a linear annealing schedule such that at each epoch, *e* the cross-entropy term was multiplied by 1 − min(*e/*(max. epochs), 1).

Networks were trained on batches of 4 512x512 pixel crops, drawn randomly from the training set. During training, the training set was augmented by random horizontal and vertical flips (*p*_flip_ = 0.5), random rotations (± 15^*°*^), and random affine shears (± 15^*°*^along both x- and y-axes).

Separate networks were trained for each species.

### D Tracking and Counting Cell Groups

#### 1 Cell Group Detection

Groups of cells were detected by thresholding the (corrected) height images (see VII A) such that all pixels above a set threshold height were called as part of a group while all pixels below were marked as background. Groups were then determined by labeling connected components in the thresholded image. Connected components smaller than the average area of a single cell (100 pixels for *M. xanthus*) were discarded. Ideally, the height threshold used for this determination would be a very small value (*e*.*g*. 1 nm) above the detected surface, which is by definition set as reference 0 nm. We found that doing so was not robust to experimental noise and surface heterogeneities while also causing groups that split apart to briefly, apparently “re-merge” due to small height variations just after splitting. To balance faithful group segmentation and robustness to such noise, we set the threshold to 1*/*2 the average cell height (for *M. xanthus*, 250 nm).

#### 2 Group Tracking

Groups were tracked over time by on a frame-by-frame basis, linking adjacent pairs of frames until a full time series of tracks had been accumulated. To do so, we used the pixel masks generated by our group detection algorithm (see VII D 1) and for each frame pair, constructed a bipartite graph where vertices represent labeled group masks and edges connect the vertices of masks in adjacent frames that have a non-zero number of overlapping pixels. We chose an optimal frame-to-frame matching of such masks by computing the maximum weight matching of the just-constructed bipartite graph. Sometimes, such groups either split into two, smaller groups or merged with adjacent groups to create a single, larger group. We maintained a “lineage tree,” that kept track of which groups were the product of such events. Merge events are detected by allowing unmatched groups in the forward direction to their best match, regardless of whether or not this matched group has already been matched to another group in the original frame. Splitting events are detected by running the matching in reverse and repeating the procedure done to detect merges. This tree was used when inferring the number of cells in each group (see VII D 3). To detect individual tracks within these tree-structured graphs, we enumerated all straight paths in the graph by depth-first search. Starting from the tree root, a consecutive string of vertices with only a single in- and out-neighbor were labeled as a track. When a node with more than 1 out-neighbors was detected (corresponding to a split event), that node was called as being a part of the current track while a new track was initiated at each out-neighbor. When a node with more than 1 in-neighbors was detected (corresponding to a merge event), the track was terminated at the current node. The search was continued until all vertices were labeled as being part of a track.

#### 3 Cell Counting in Groups

We found that using laser and height images, alone, was not of sufficiently high resolution to accurately segment and track cells over time within groups^3^. To remedy this, we instead opted to simply “count” cells in tracked groups. To do this, predicted cell masks were combined with the detected cell groups, yielding a list of tuples for each group containing the frame number and the number of cells detected within the group at that frame.

We first calibrated the fidelity of these counts by detecting cell groups in our ground truth, manually labeled dataset (see VII D 1). We then segmented the images using our convolutional neural network and compared the detected number of cells in each group to the ground truth (Fig. 25a). We found that, for a given group size *N*, the average predicted size over all predicted groups was a good match to the ground truth (Fig. 25b, top) while the variance of these predictions was underdispersed relative to a Poisson counting model (Fig. 25b, bottom). We found that the distribution of predicted sizes *P* was well-described by a generalized, underdispersed Poisson distribution[23]:

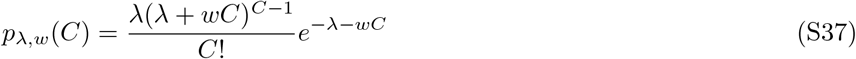

which has mean 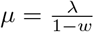 and variance 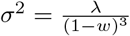. We calibrated the dispersion parameter, *w*, by finding the optimal

*w* that would allow us to accurately predict the ground truth by computing *P* = ┌*µ*┐ of the fitted distribution. We found that *w* = −0.94 allowed us to perfectly predict the number of cells in groups of all sizes *N* ≤ 16 except for *N* = 15 (Fig. 25c).

For cell groups that never merged or split with other groups, the number of cells in that group was inferred by fitting the distribution of number of detected cells in each frame to Eqn. S37 with *w* = −0.94 and computing *P* = ┌*µ*┐ as was done in the calibration procedure, above. For cell groups that merged of split with other groups, we leveraged the dependency structure implied by these events to better inform our estimation. At each merge/split event, we performed joint inference on the larger and two smaller groups by asserting that the count for the largest group (the merged group or the group that splits apart), must be the sum of the two smaller ones. As was shown by Consul and Jain [24], 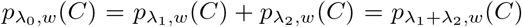 and thus we can obtain all three *λ*_*x*_ parameters for the 3 groups (and then compute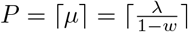) by simultaneous fitting of all the data for all 3 groups.

### E Single cell tracking

#### 1 Candidate cell enumeration

We developed a simple enumeration scheme in which labeled regions found by watershed segmentation of a segmentation neural network’s output probability “image” that were in direct contact with other labeled regions are grouped together from which a list of mutually exclusive candidate solutions is generated per-such-group. Direct contact was established between potentially-adjacent regions by computing each region’s exterior boundary pixels and checking for overlapping pixels between pairs of region borders. Exterior boundary pixels were calculated by morphological dilation of each region followed by subtraction of the region’s original pixels. Groups of continuously touching regions^4^ were labeled as such. For each of these groups, a list of mutually exclusive candidate solutions is generated. To generate the list, we assume that the regions generated by the initial watershed segmentation cannot be further split to yield valid regions. We further assume that each pair of touching regions could have come about through the aberrant splitting of a single cell. We enumerate all such individual cases within a group along with all possible combinations within which such a merge can exist alongside the other cells in said group. The above-described enumeration is then done recursively until the group is merged into a single region. This scheme yields a list of all possible cells that could have arisen from within the group, as well as a list of mutually exclusive combinations of cells that could have arisen from the network output and watershed segmentation.

#### 2 Non-motile pairs of E. coli & F. johnsoniae

To track post-division daughter cells in non-motile *E. coli* and *F. johnsoniae*, we first generated a series of predicted cell probabilities using our convolutional neural network (see VII C). At each frame, these probabilities were used to generate a list of candidate cells using the algorithm described, above (see VII E 1). Simultaneously, candidate cell groups were tracked (see VII D). Cell groups were then linked to candidate cell segmentations by detecting pixel overlaps between the two sets of masks. At each frame and for each group, we checked whether that group contained a candidate segmentation with exactly two cells and if so, included it in a running list of such frames for that group. All such frames were then compiled and cells from adjacent frames in this sequence were linked to one another by maximizing the total intersection over union of cell masks between the masks of these two frames. This maximization was accomplished by treating the adjacent frame pairs as a bipartite graph with nodes representing cell masks and edges representing the (negative) intersection over union value between the two masks and using minimum weight matching.

Calculation of the relative centroid-to-centroid distance for these pairs (shown in Fig. 2i) was done by first extracting the centroid of each cell mask at each frame and computing the Euclidean distance between them. This yields an absolute distance between the two centroids. The sequence of such distances was then smoothed by a 3-frame rolling mean filter for each pair of daughter cells (Fig. 27a). For each pair, the time at which the centroids were maximally separated was found and set as a reference 0 (Fig. 27b). Finally, we divided all the absolute centroid-to-centroid distances by the value of maximum separation to yield a relative centroid-to-centroid distance (Fig. 27c). Note that in Fig. 2i, time was recalculated relative to that shown in Fig. 27c by setting the minimum value to be 0 instead of the reference time to which all of the sequences were aligned.

#### 3 Motile M. xanthus

Motile pairs of *M. xanthus* cells within a phase (as shown in Fig. 3e) were first manually annotated in a single frame at the start of the time series. For each subsequent frame, all potential cell segmentations were enumerated using the algorithm described, above (see VII E 1). These were then linked in time using a novel algorithm that solves a constrained linear sum optimization problem across a series of frames.

### F Wetting Profile Measurement

To measure the capillary length of cells in experiment, we first isolated single cells (or pairs, as in Fig. 2) using our group tracking and counting algorithm (see VII D 2). The contour of the “group mask” found by thresholding of the height map was extracted and the major axis of an ellipse fitted to these contour pixel coordinates was extracted by singular value decomposition of the contour pixel coordinates. To do this, the *n* contour pixel coordinates were mean-centered and concatenated into a 2*xn* matrix and the singular value decomposition of this matrix was computed. From this, we constructed a transformation that would take scalar values in the interval [-1,1] and map them onto the major axis of the cell by taking the dot product of the matrix of left singular vectors, *U*, with the matrix *Is* where *I* is the (appropriately-shaped) identity matrix and *s* is a vector of singular values and scaling the resulting matrix by 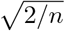 (Eqn. S38).

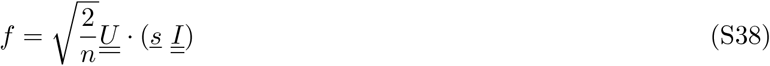

We used the resulting transform to generate points along the middle half — ± 1*/*4 of the major axis length from the contour center, 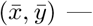 by computing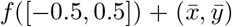. For each generated point along the major axis, we generated a linearly-spaced sampling line along the direction perpendicular to the major axis at that point. We sampled each point of this line by linearly-interpolating from the height map at the specified pixel coordinates. These values were then accumulated and averaged by distance from the major axis to yield a single curve of height vs. distance from major axis. This curve was then fit to a circular arc, as discussed in I C.

We note that fitting of a circular was unstable and often yielded invalid fits on our experimental data. For some analyses (Fig. 22, 12), we fit the measured meniscus profile to an exponential decay (Eqn. S39 and extracted the fitted decay length, *ℓ*_*c*_.

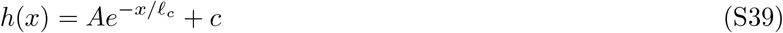

This measurement shows the same qualitative relationship with osmotic pressure as that shown in Fig. 1f (Fig. 23).

## SUPPLEMENTARY TABLES

**Supplementary Table 1:**
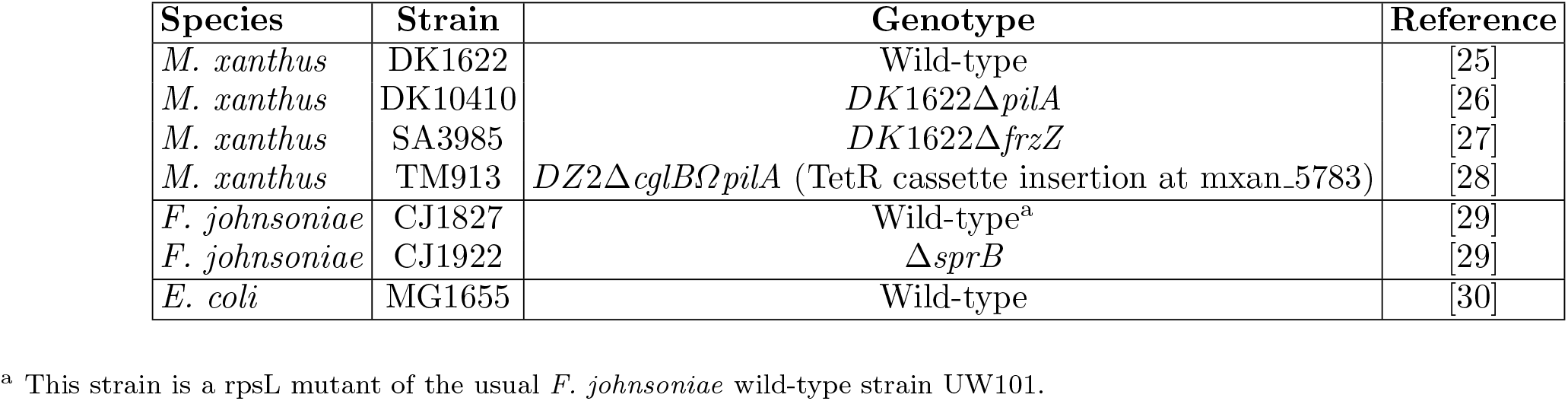
Strains used in this study.

**Supplementary Table 2:**
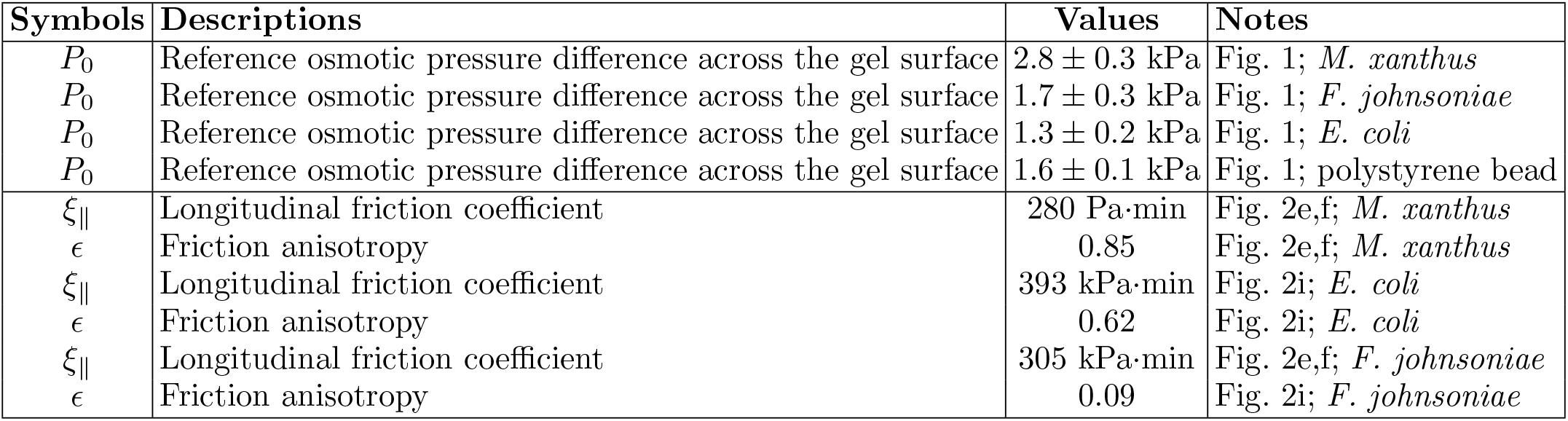
Model parameters obtained by fitting to experimental data.

**Supplementary Table 3:**
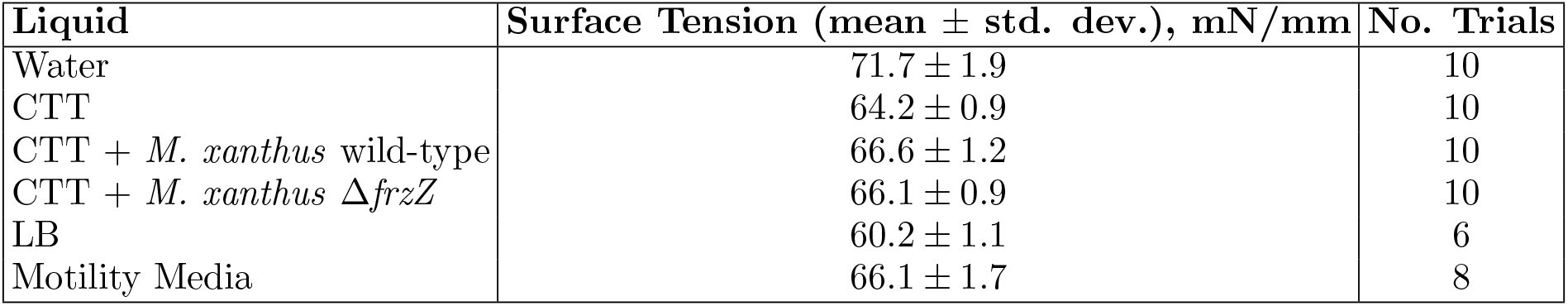
Measured surface tensions for all media used in this study.

## SUPPLEMENTARY FIGURES

**Supplementary Fig. 1.**
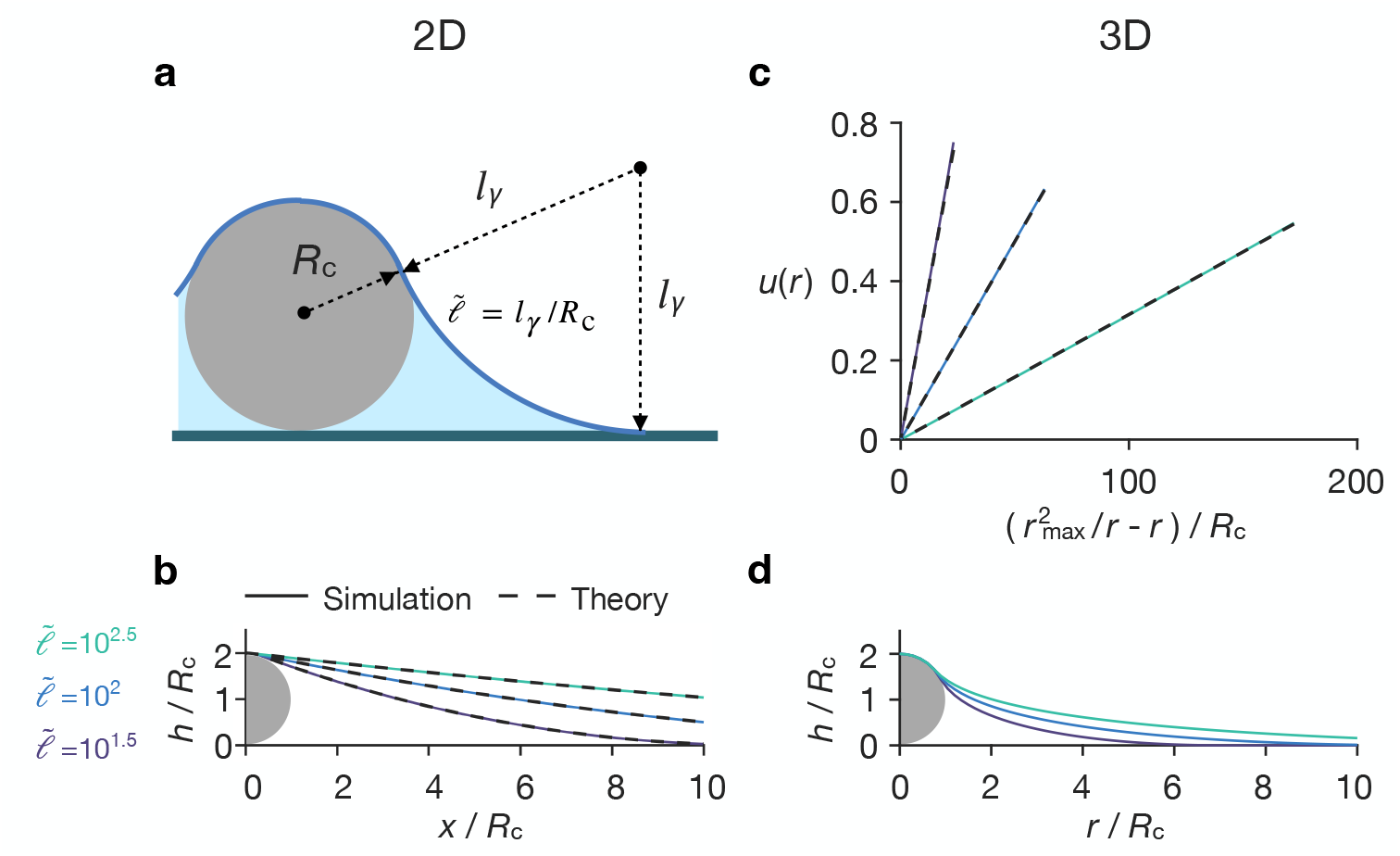
The shape of the water meniscus around a circular or spherical cell. (**a**) and (**b**) show results for a 2D circular “cell”, and (**c**) and (**d**) show results for a 3D spherical “cell”. (**a**) Schematic of the 2D height profile given by Eq. (S5). Color code as in main Fig. 1. *R*_c_ denotes the radius of the 2D circular “cell” and *l*_*γ*_ denotes the capillary length of the meniscus. 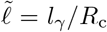 denotes the normalized capillary length. (**b**) Simulation (solid) and analytical (dashed) results for the equilibrium height profile of the water meniscus around a 2D circular “cell”. (**c**) Height profiles obtained from the simulations of a spherical “cell” satisfy Eq. S7. Solid and dashed lines indicate the simulation and analytical results, respectively. See Sec. I B for details. (**d**) Simulation results for the equilibrium height profile of water meniscus around a 3D spherical “cell”. In panels **b** – **d**, colors indicate the designated values of 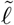.

**Supplementary Fig. 2.**
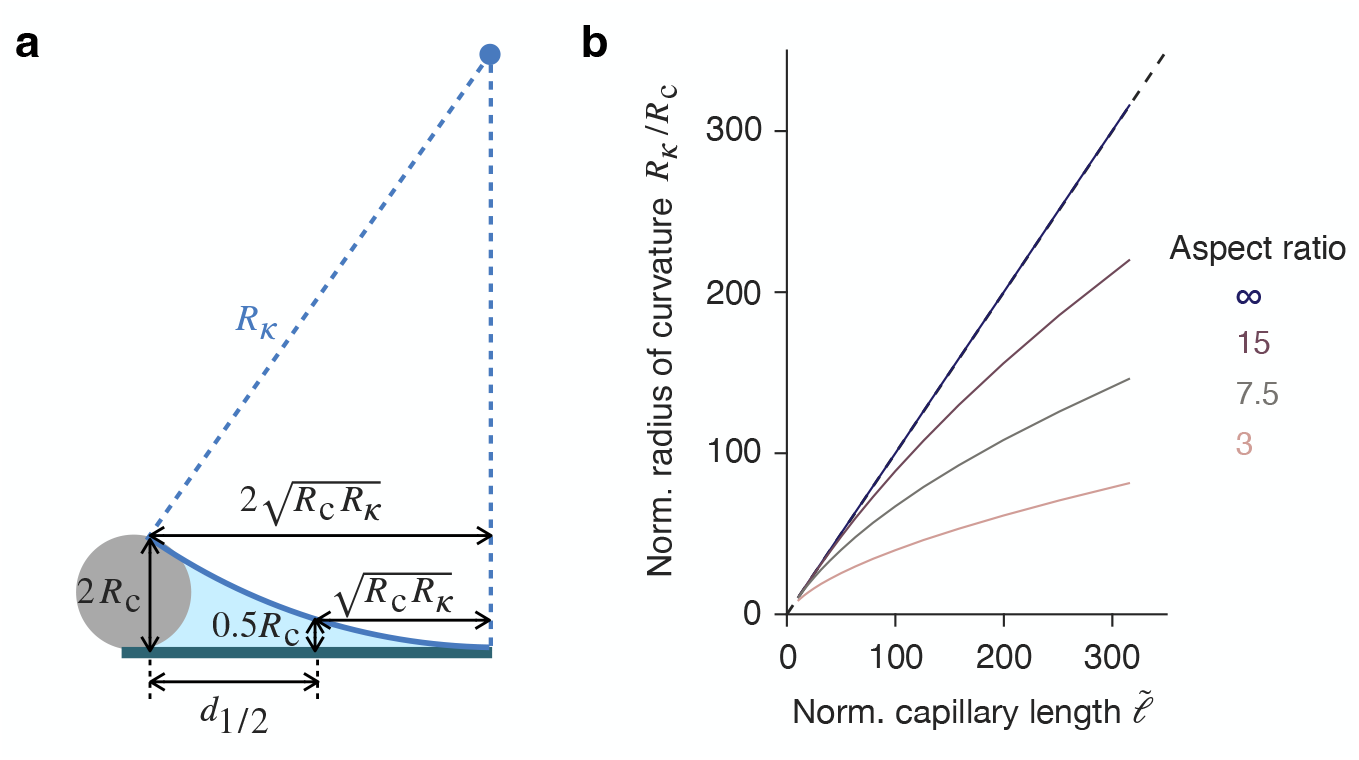
The midplane height profile varies with cell aspect ratio. (**a**) Schematic of the geometry used to estimate the radius of curvature *R*_*κ*_ of the midplane water height profile. Color code as in main Fig. 1. See Sec. I C for details. (**b**) The radius of curvature *R*_*κ*_, normalized by cell radius *R*_c_, plotted against normalized capillary length 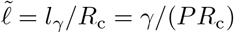 at the designated cell aspect ratios. Black dashed line indicates *R*_*κ*_ = *l*_*γ*_.

**Supplementary Fig. 3.**
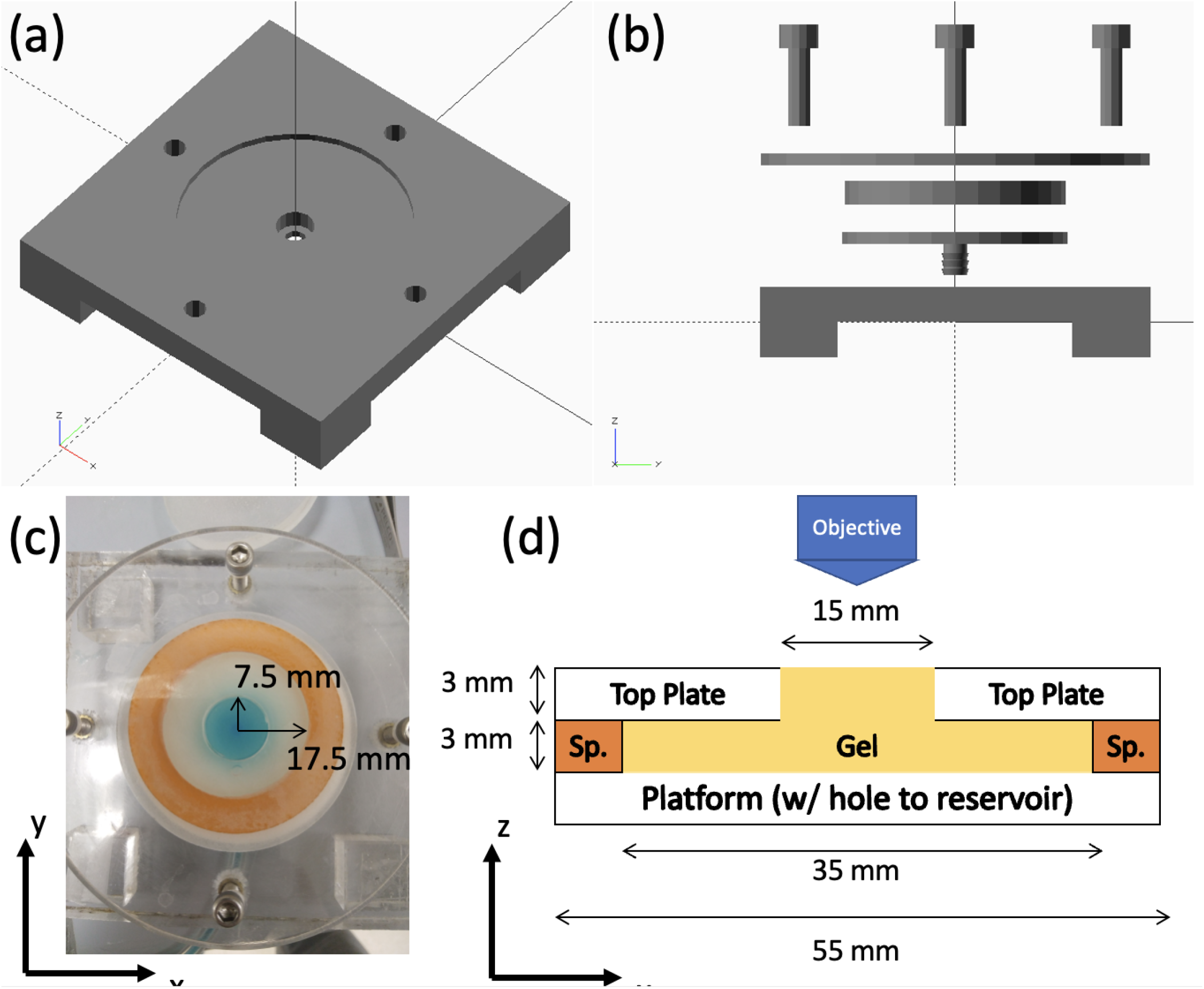
**(a)** 3d view of the base plate. Note the middle cutout has a lower lip that is used to support the outer edge of the Luer lock that will connect to the reservoir. **(b)** Exploded side view of the device, with all pieces shown in the order of their assembly. **(c)** Top view of an assembled device with a gel, with outer radii for the inner through hole and the top plate, itself, shown. **(d)** Side view schematic of the top part of the device, with relevant dimensions for all pieces shown.

**Supplementary Fig. 4.**
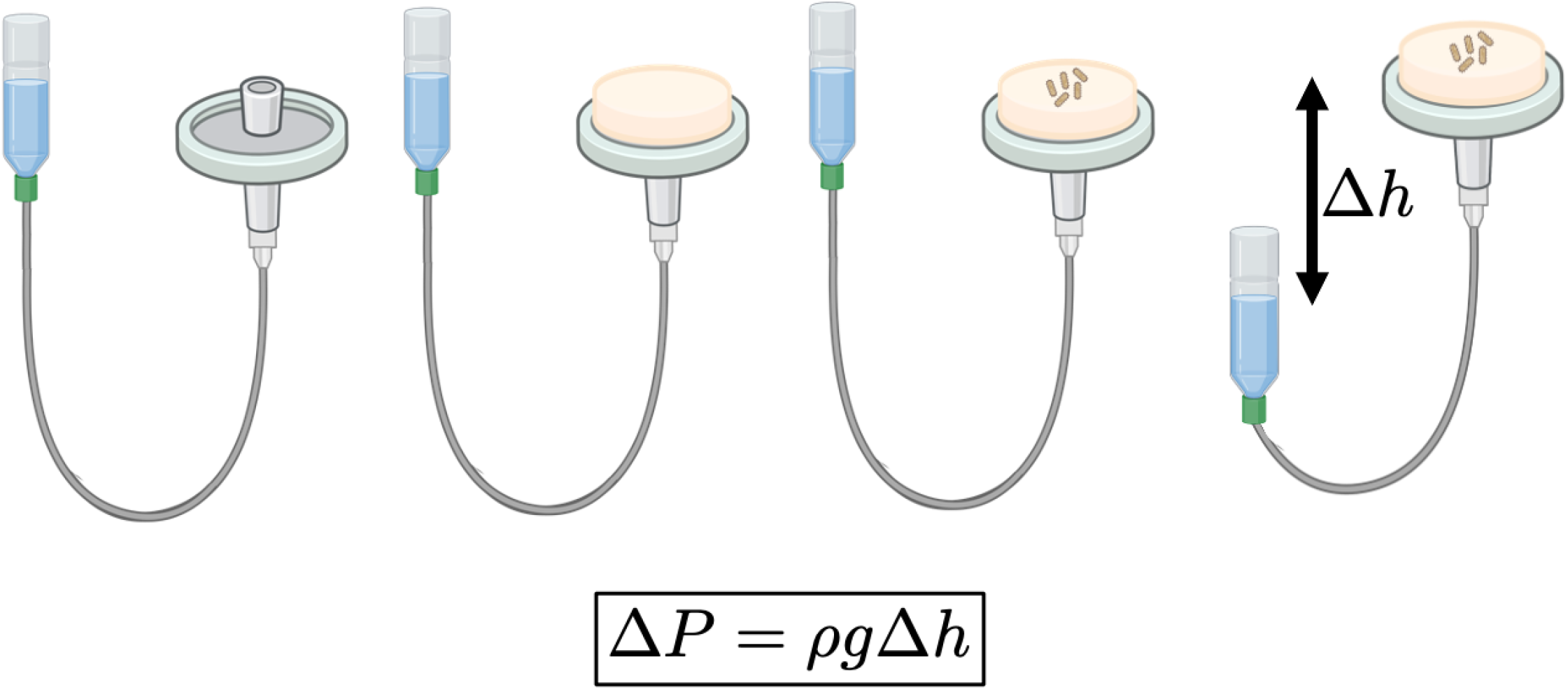
Cartoon showing the hydrostatic pressure control device and its basic operating principle. A plate with a hole is connected to a reservoir of water (or liquid media). The gel is placed on top of the plate, directly coupling it to the reservoir. By moving the reservoir up or down relative to the top of the gel surface, a hydrostatic head is applied to the bottom of the gel. This head then acts on top of the osmotic pressure inherent to the gel, either “pushing liquid into” or “pulling liquid out of” the gel.

**Supplementary Fig. 5.**
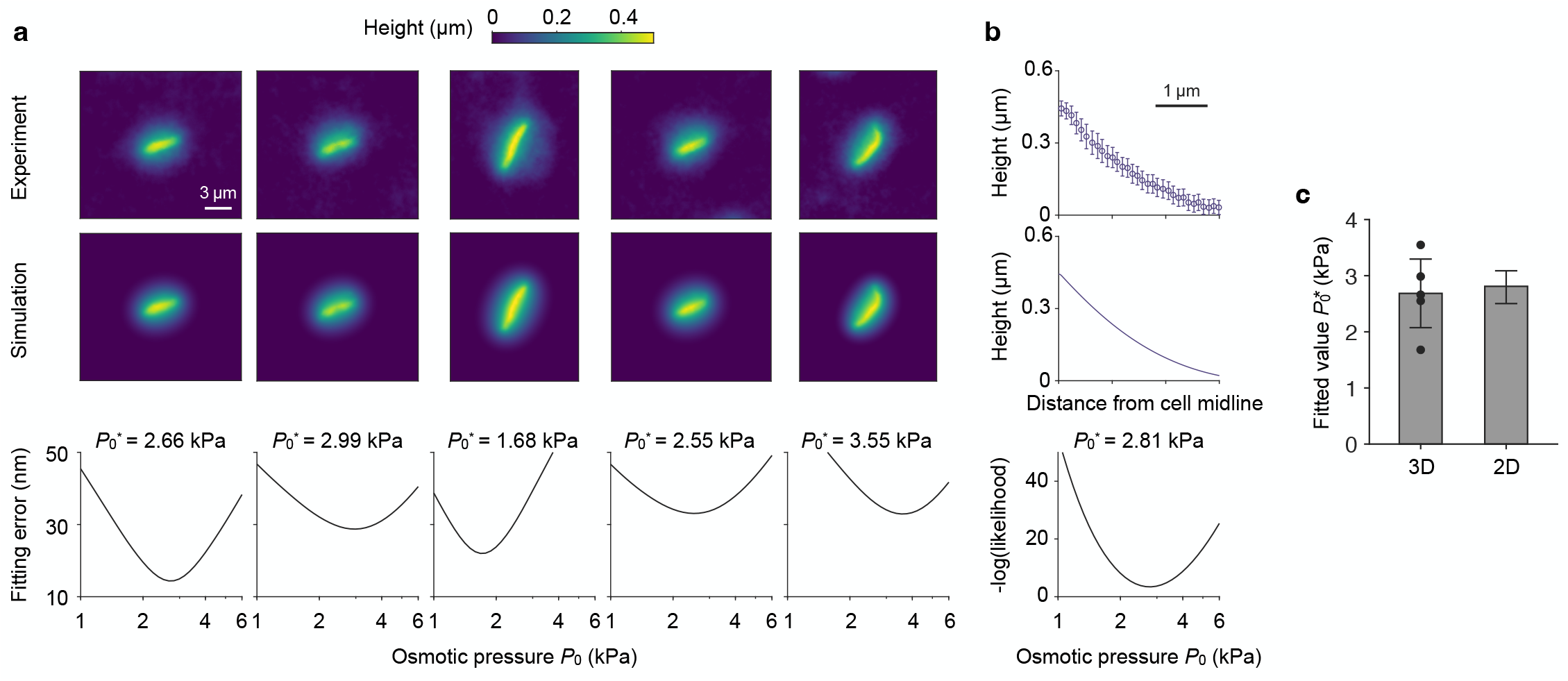
Fitting the 3D and 2D height profiles of the water menisci yield similar results. (**a, b**) The osmotic pressure difference *P*_0_ is determined by fitting the 3D (**a**) or 2D (**b**) height profiles of the modeled water menisci to those measured in experiments. *Top* panels show the experimental measurements and *middle* panels show the simulation results with the optimal fitting parameters 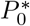. *Bottom* panels show the fitting errors at varying values of *P*_0_. In **a**, each column represents an individual cell and the fitting error is measured by the root-mean-square height difference between the model and the experiment. In **b**, the height profile is measured in the mid-plane perpendicular to the cell body axis, and the fitting error is measured by the negative log-likelihood of the fitting. See Sec. I D for details. (**c**) Bar plot for the fitted values of *P*_0_. Fitting the 3D and 2D height profiles yield *P*_0_ = 2.69 ± 0.61 kPa and *P*_0_ = 2.81 ± 0.29 kPa,respectively.

**Supplementary Fig. 6.**
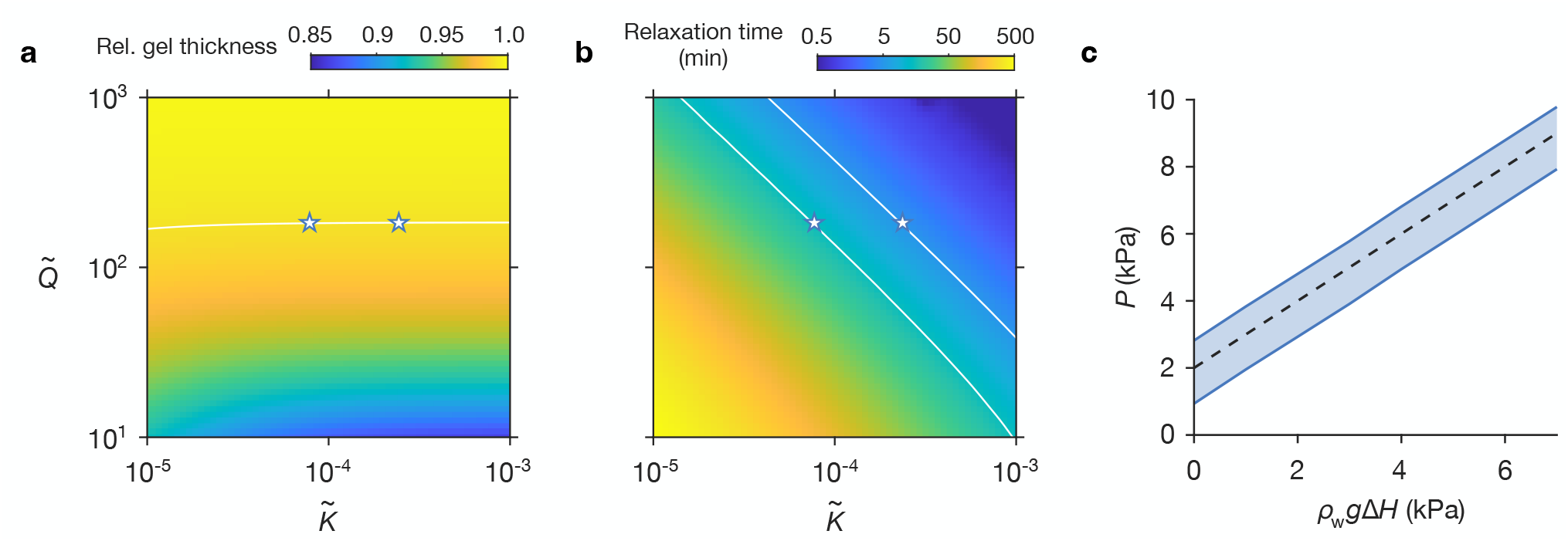
Water evaporation leads to a osmotic pressure gradient across the gel. (**a**) Relative gel thickness *α* and (**b**) the relaxation time *τ*_ss_ of the model (Eq. S10) at varying values of 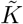 and 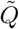. See Sec. I E for details. White curve in **a** denotes *α* = 0.99. White curves in **b** denote *τ*_ss_ = 5 min and *τ*_ss_ = 15 min, respectively. White stars show the intersection points between the curve in **a** and the two curves in **b**, which determine the range of values for 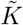 and 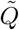 in experiment. (**c**) The osmotic pressure difference *P* across the gel surface at varying height differences Δ*H* between the surface of the gel and the water level of the reservoir. The top and bottom curves represent, respectively, the results for the parameters indicated by the left and right stars in **a** and **b**. Black dashed line indicates a slope of 1.

**Supplementary Fig. 7.**
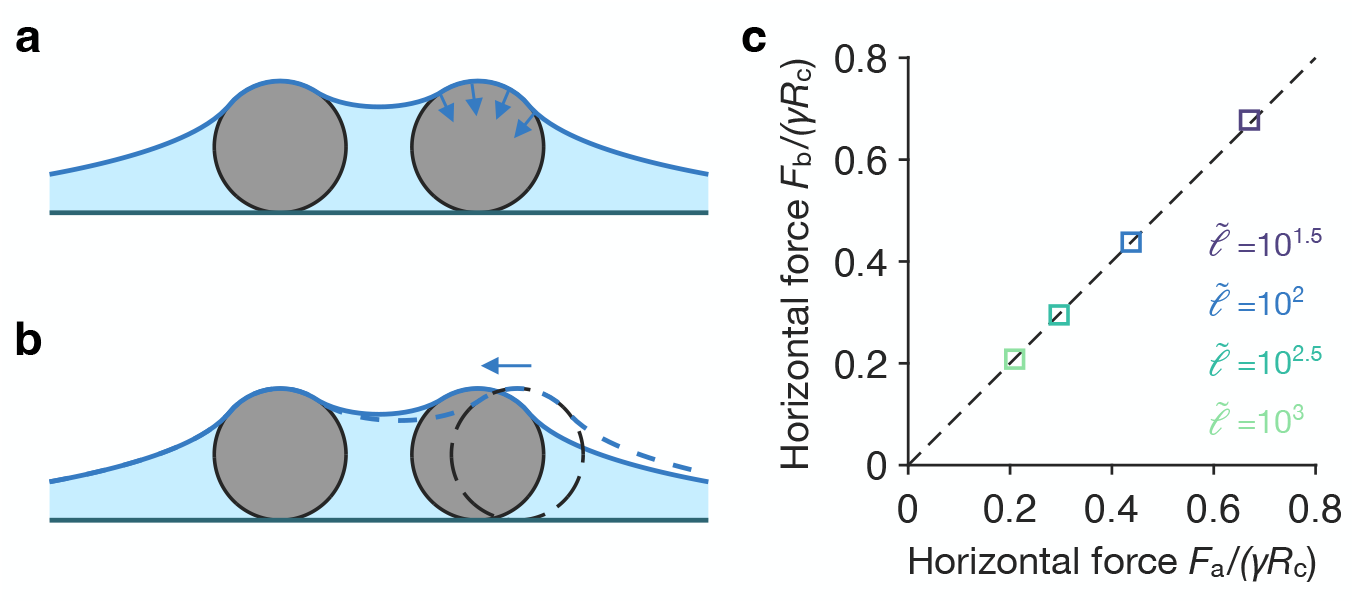
Calculation of the in-plane capillary force. (**a, b**) Schematics of two methods for calculating the capillary force exerted on the cell. The force can be computed from (**a**) a force integral over the region where the water meniscus is in contact with a cell, or from (**b**) the free-energy difference upon displacement of a cell (dashed). See Sec. I F for details. (**c**) Comparison between the force calculated using the method shown in **a** (denoted by *F*_*a*_) and the force calculated using the method shown in **b** (denoted by *F*_*b*_). The forces are normalized by *γR*_c_. For reference, the dashed line shows *F*_*b*_ = *F*_*a*_. Colors indicate the designated values of the normalized capillary length 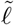.

**Supplementary Fig. 8.**
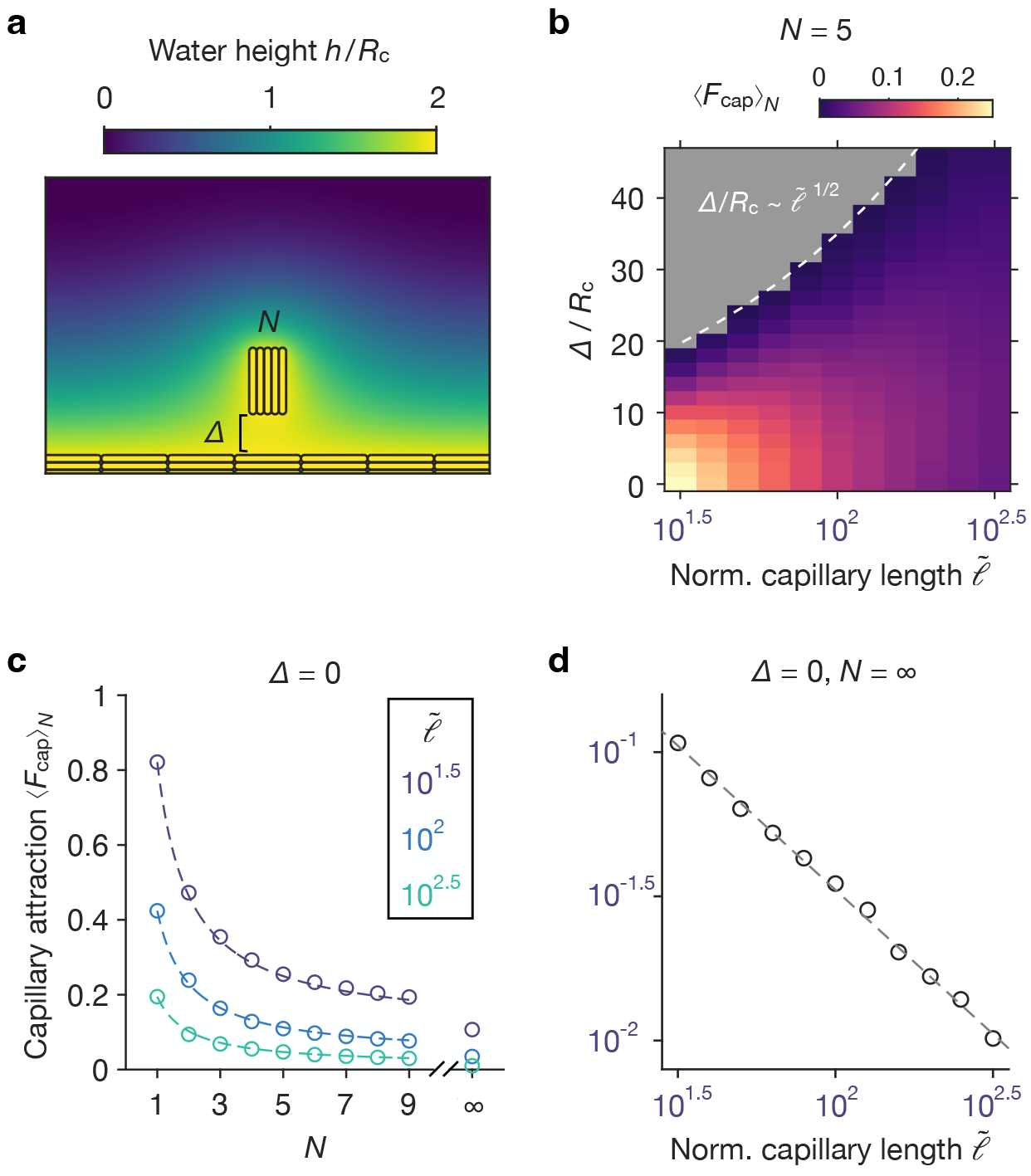
Characterizing the magnitude and range of capillary attraction. (**a**) Illustration of the simulation setup. A group of *N* cells is positioned a distance Δ away from an infinitely large colony. The average capillary force acting on each cell is denoted by ⟨*F*_cap_⟩_*N*_. (**b**) Capillary attraction ⟨*F*_cap_⟩_*N*_, normalized by *γR*_c_, at varying distance Δ and normalized capillary length 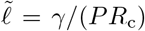 for *N* = 5. Gray denotes that ⟨*F*_cap_⟩_*N*_ = 0. White dashed curve indicates 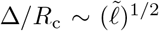. (**c**) Capillary attraction ⟨*F*_cap_⟩_*N*_, normalized by *γR*_c_, at varying cell number *N* and normalized capillary length 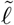 (color) for Δ = 0. Circles indicate simulation data, and dashed curves indicate ⟨*F*_cap_⟩_*N*_ = ⟨*F*_cap_⟩_∞_ + (⟨*F*_cap_⟩_1_ − ⟨*F*_cap_⟩_∞_)*/N*. (**d**) Capillary attraction ⟨*F*_cap_⟩_∞_, normalized by *γR*_c_, at varying normalized capillary length 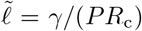 for Δ = 0. Circles indicate simulation data, and the gray dashed line indicates 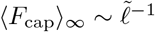.

**Supplementary Fig. 9.**
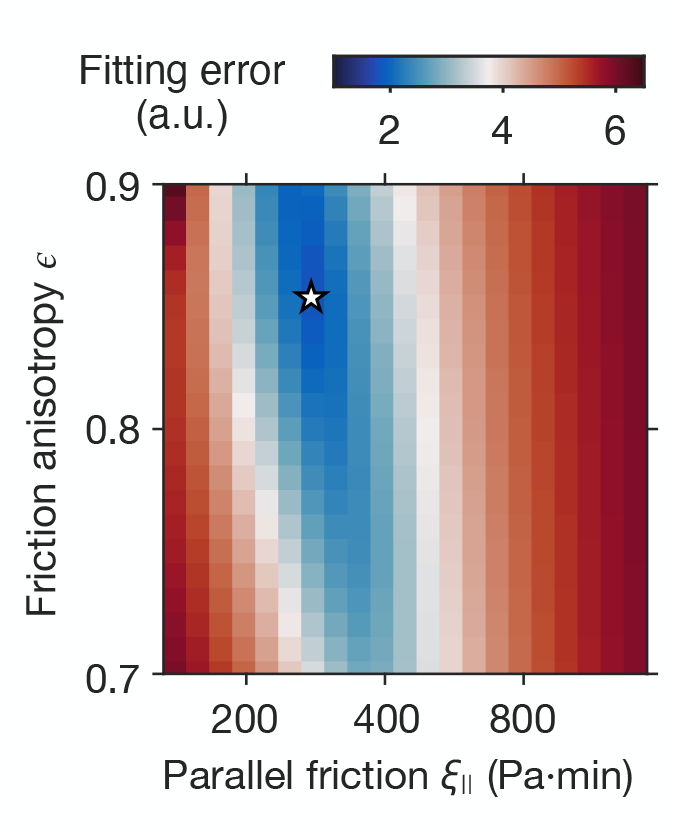
Fitting friction coefficients for *M. xanthus* cells. Fitting error of the merger dynamics between simulation and experiment at varying friction coefficient *ξ*_∥_ and friction anisotropy *ϵ* (see Sec. II B). Star denotes the optimal fitting values.

**Supplementary Fig. 10.**
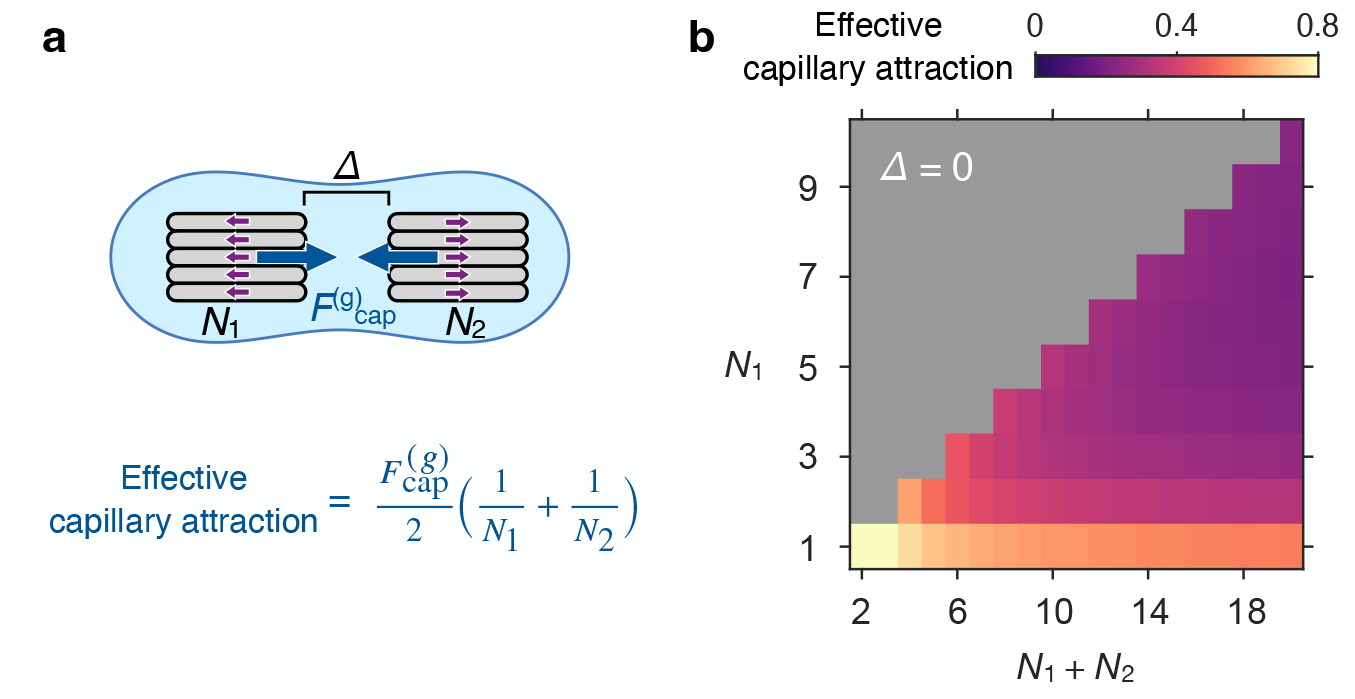
Effective capillary attraction varies with the partition of cell groups. (**a**) Schematic of the capillary attraction 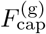 between two subgroups of *N*_1_ and *N*_2_ cells that propel in the opposite directions. Δ denotes the distance between the two subgroups. The effective per-cell capillary attraction 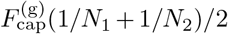 indicates the minimal propelling force for the two subgroups to split up. (**b**) Effective capillary attraction 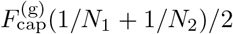, normalized by *γR*_c_, at varying cell numbers *N*_1_ and *N*_2_ for Δ = 0. Gray indicates *N*_1_ *> N*_2_ which is symmetric to *N*_1_ *< N*_2_.

**Supplementary Fig. 11.**
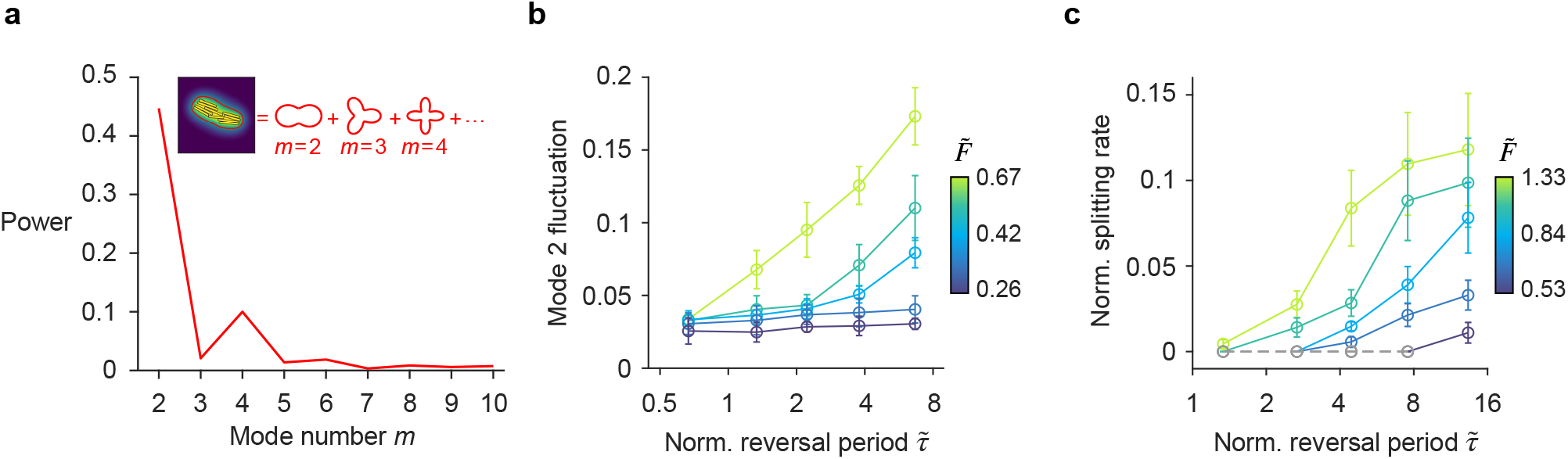
Elongation and splitting of a cell group. (**a**) Power spectrum *c*_*m*_ of contour deformation for the cell group shown in the inset. Deformation modes *m* = 2, 3, and 4 are illustrated in the inset. See Sec. II D for details. (**b**) Mode 2 fluctuation, defined as 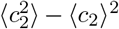, and (**c**) splitting rate of a 10-cell group at varying normalized reversal period 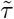 and normalized self-propelling force 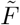. In **b** and **c**, error bars represent standard deviations of 5 replicate simulations. In **c**, the dashed lines and circles indicate that the cell group does not split over the course of the simulation.

**Supplementary Fig. 12.**
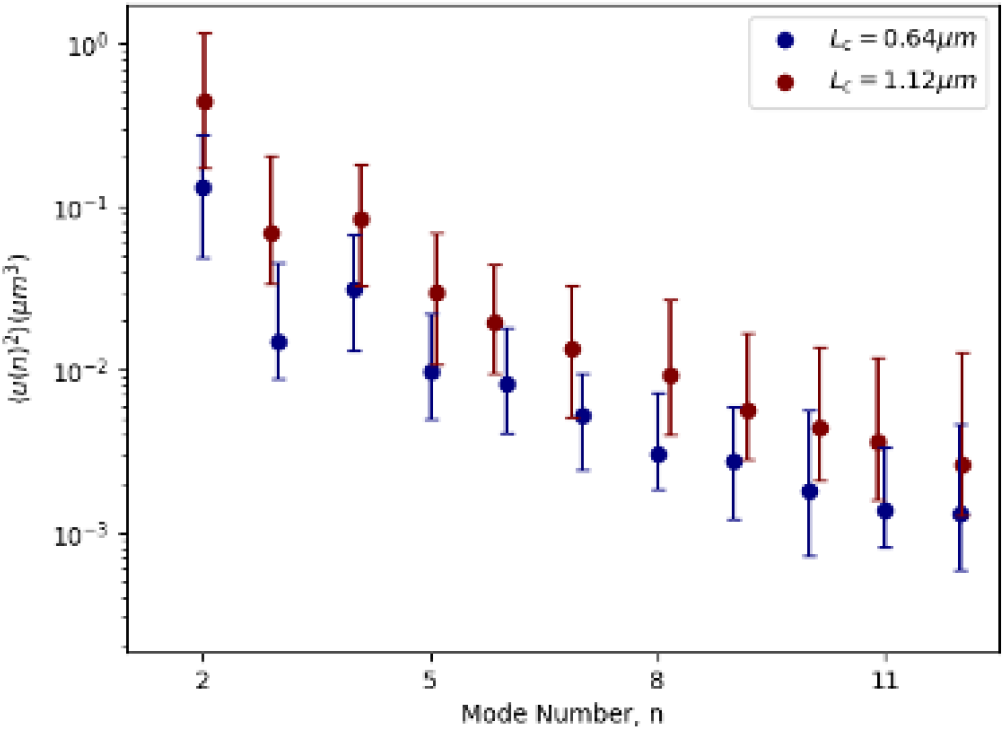
Power spectrum of contour deformations for small groups of cells vary with capillary length. Power spectrum of contour deformations for groups of size *N* = 5 *M. xanthus* cells, measured at different capillary lengths. Decreasing capillary length further confines the group, causing reduced fluctuations.

**Supplementary Fig. 13.**
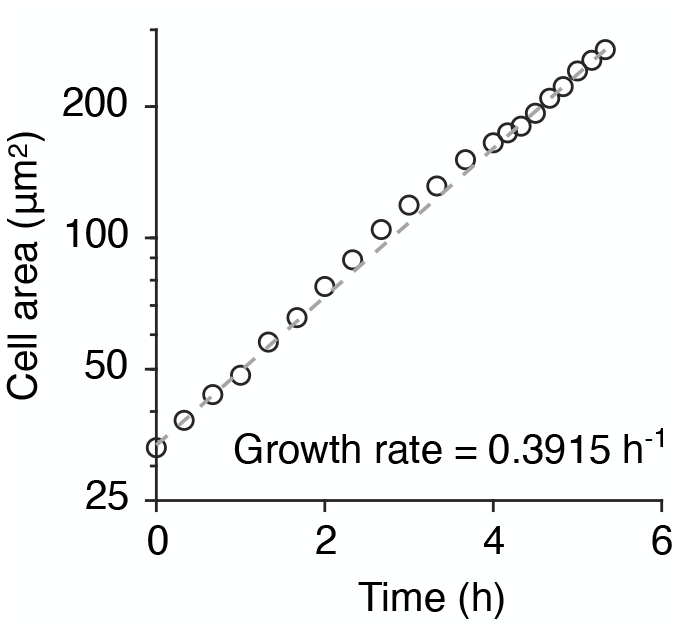
Fitting the areal growth rate of *E. coli* colonies. Area covered by cells as a function of time. Circles indicate experimental measurements and the dashed line indicates an exponential fit with the designated growth rate.

**Supplementary Fig. 14.**
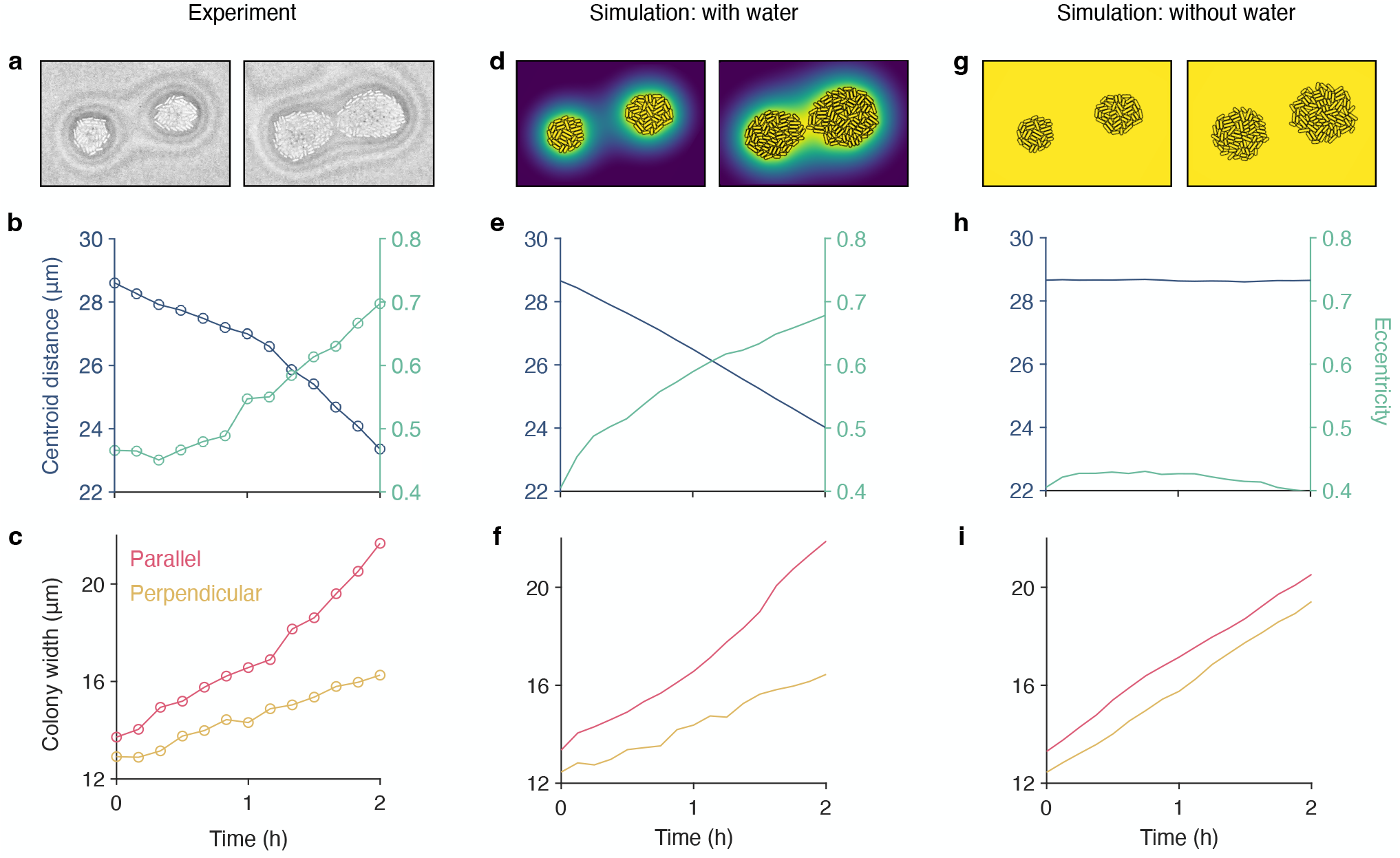
Capillary forces facilitate merger of growing *E. coli* colonies. The growth dynamics of two adjacent colonies are compared between (**a**–**c**) experiment, (**d**–**f**) simulation with water, and (**g**–**i**) simulation without water. (**a**,**d**,**g**) Snapshots of the colonies at *t* = 0 (left) and *t* = 2 h (right). (**b**,**e**,**h**) Time evolution of the centroid distance (blue) and average eccentricity (green) of the colonies. (**c**,**f**,**i**) Time evolution of the average colony widths in the directions parallel (red) and perpendicular (yellow) to the centroid-centroid connecting line.

**Supplementary Fig. 15.**
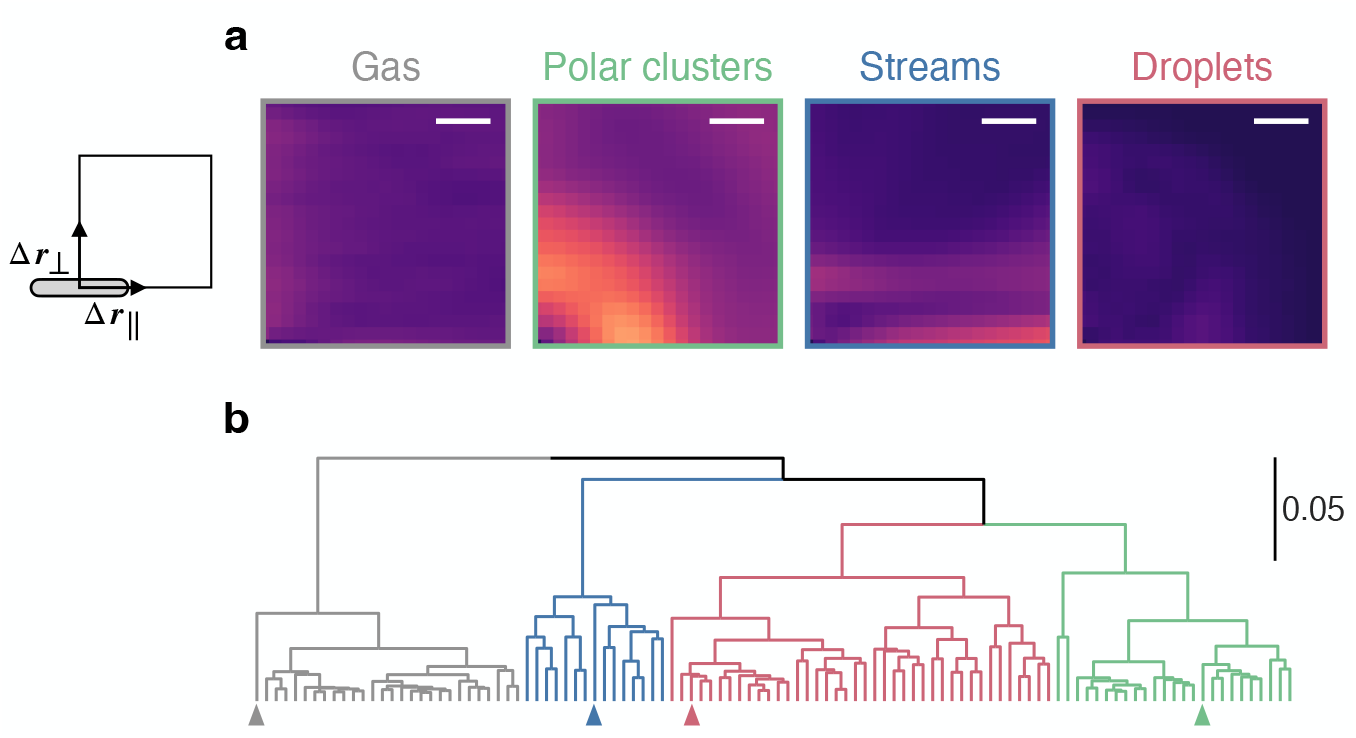
Classification of the simulated collective dynamics. (**a**) Heatmaps showing root-mean-square variation of the normalized density correlation function 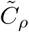 in time for the designated phases of collective dynamics. Scale bars: 3 µm. Plots correspond to main Fig. 3c,d. (**b**) The WPGMA dendrogram of the simulated collective dynamics. See Sec. III A for details. Scale bar indicates a distance of 0.05. Arrowheads indicate the four select examples shown in panel **a**.

**Supplementary Fig. 16.**
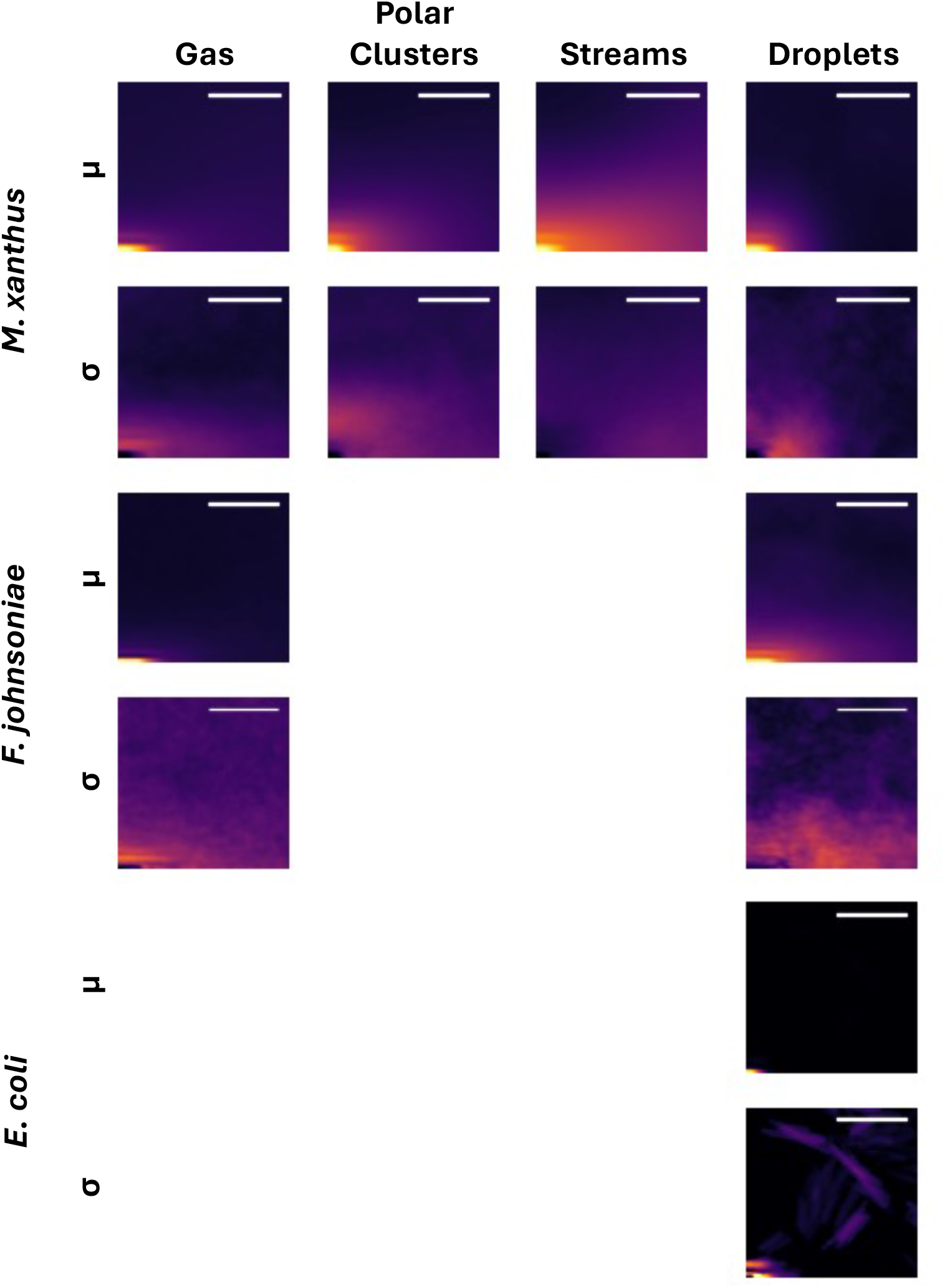
Density correlations for each collective phase found in all species studied. All examples shown were manually selected from representative experiments. Groups of two rows, one corresponding to the mean (*µ*) and the other the standard deviation (*σ*) of the density correlation function described in III B are given for each species. Scale bars: 5 *µ*m.

**Supplementary Fig. 17.**
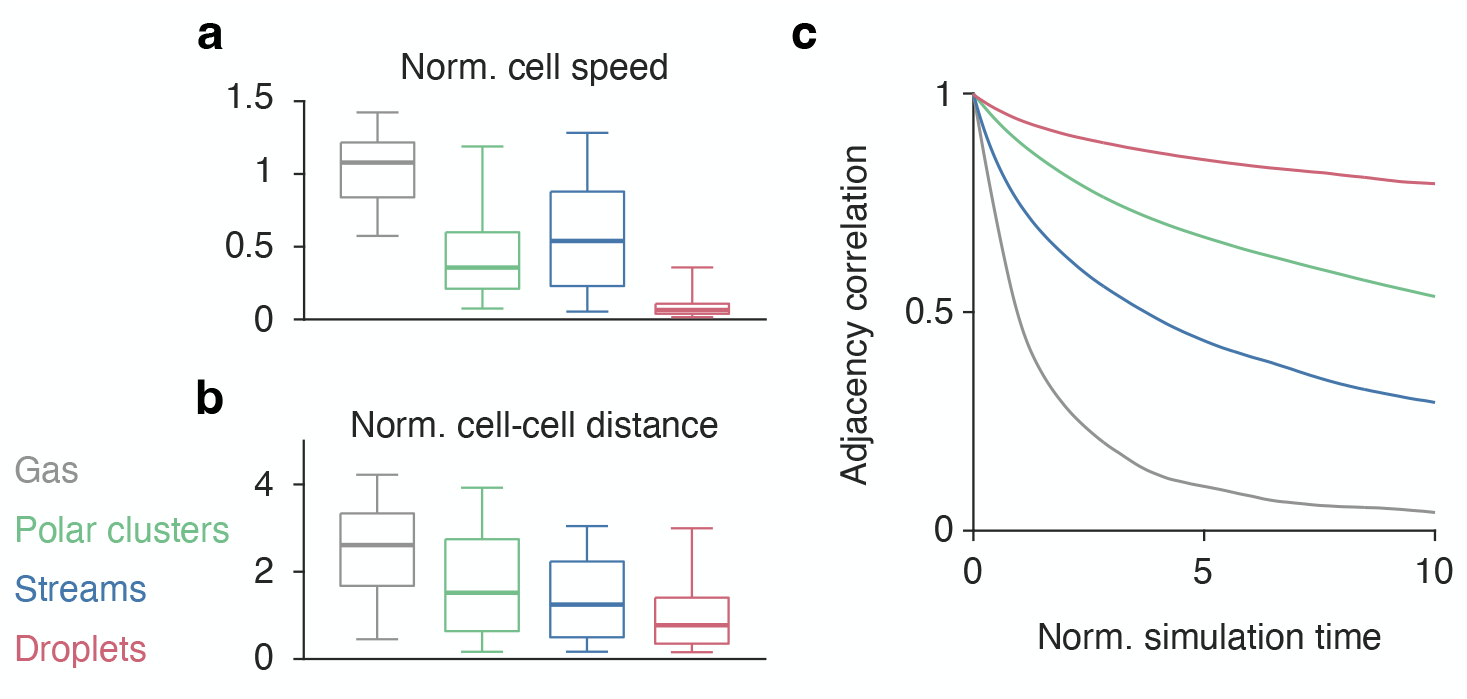
Different phases of collective dynamics display a mobility-adjacency trade-off. Box plots of (**a**) normalized cell speed and (**b**) cell-cell distance for the designated phases. Whiskers range from the 5^th^ percentile to the 95^th^ percentile. Boxes range from the 25^th^ percentile to the 75^th^ percentile. Horizontal bars inside the boxes denote the median values. (**c**) The average autocorrelation function of the cell-cell adjacency matrix (defined in Sec. III B) for the designated phases. The simulation time is normalized by the time 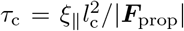 it takes for a non-reversing free-running cell to move over one cell body length. We varied three dimensionless parameters in our simulations: a normalized reversal period 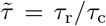, a normalized self-propulsion strength 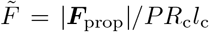 where *PR*_c_*l*_c_ is the scale of the longitudinal capillary force acting on isolated cells, and a normalized capillary length 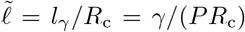. The simulation parameters are: (1) 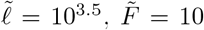, and 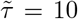 for the gas phase, (2) 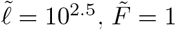, and 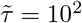 for the polar clusters phase, (3) 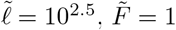, and 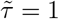 for the streams phase, and 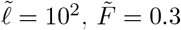 and 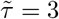 for the droplets phase.

**Supplementary Fig. 18.**
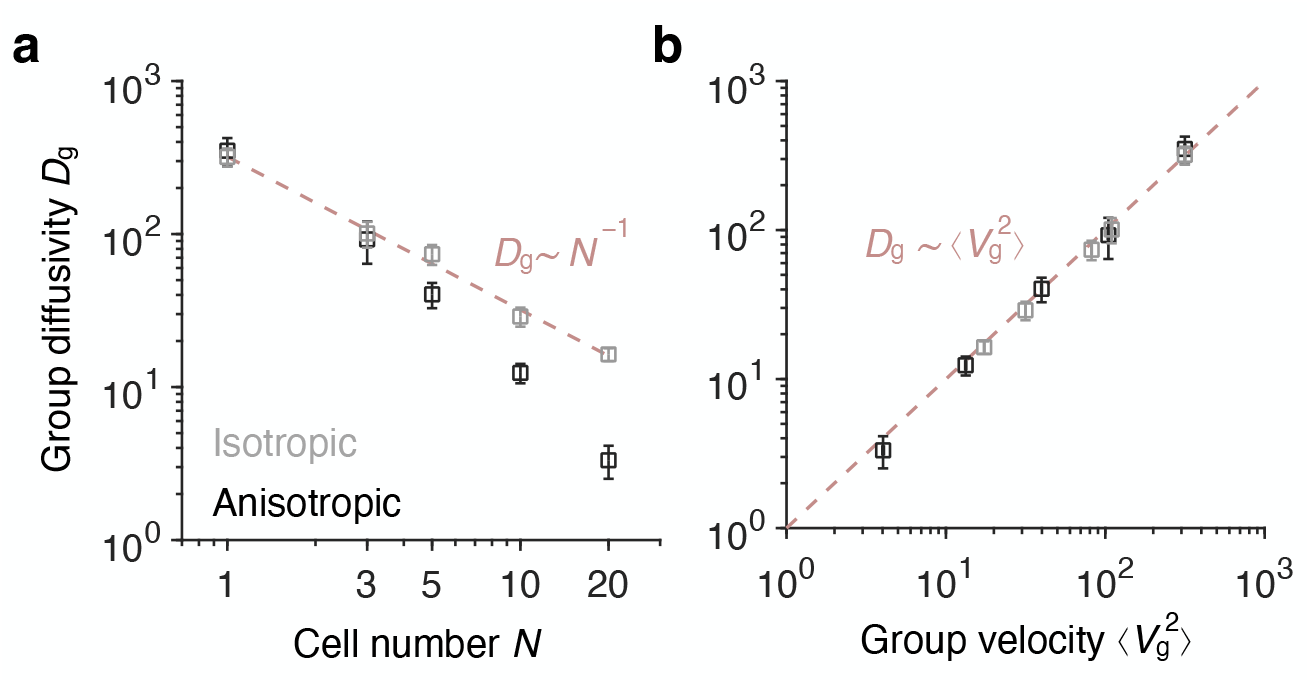
Group diffusivity decreases with increasing group size in the droplet phase. Group diffusivity *D*_g_ versus (**a**) the number *N* of cells in the group and (**b**) the mean squared velocity 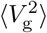 of the group. Gray and black denote simulations with isotropic and anisotropic friction, respectively. Error bars represent standard deviations of 5 replicate simulations. Red dashed lines denote *D*_g_ ∼ *N*^−1^ in **a** and 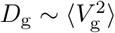 in **b**.

**Supplementary Fig. 19.**
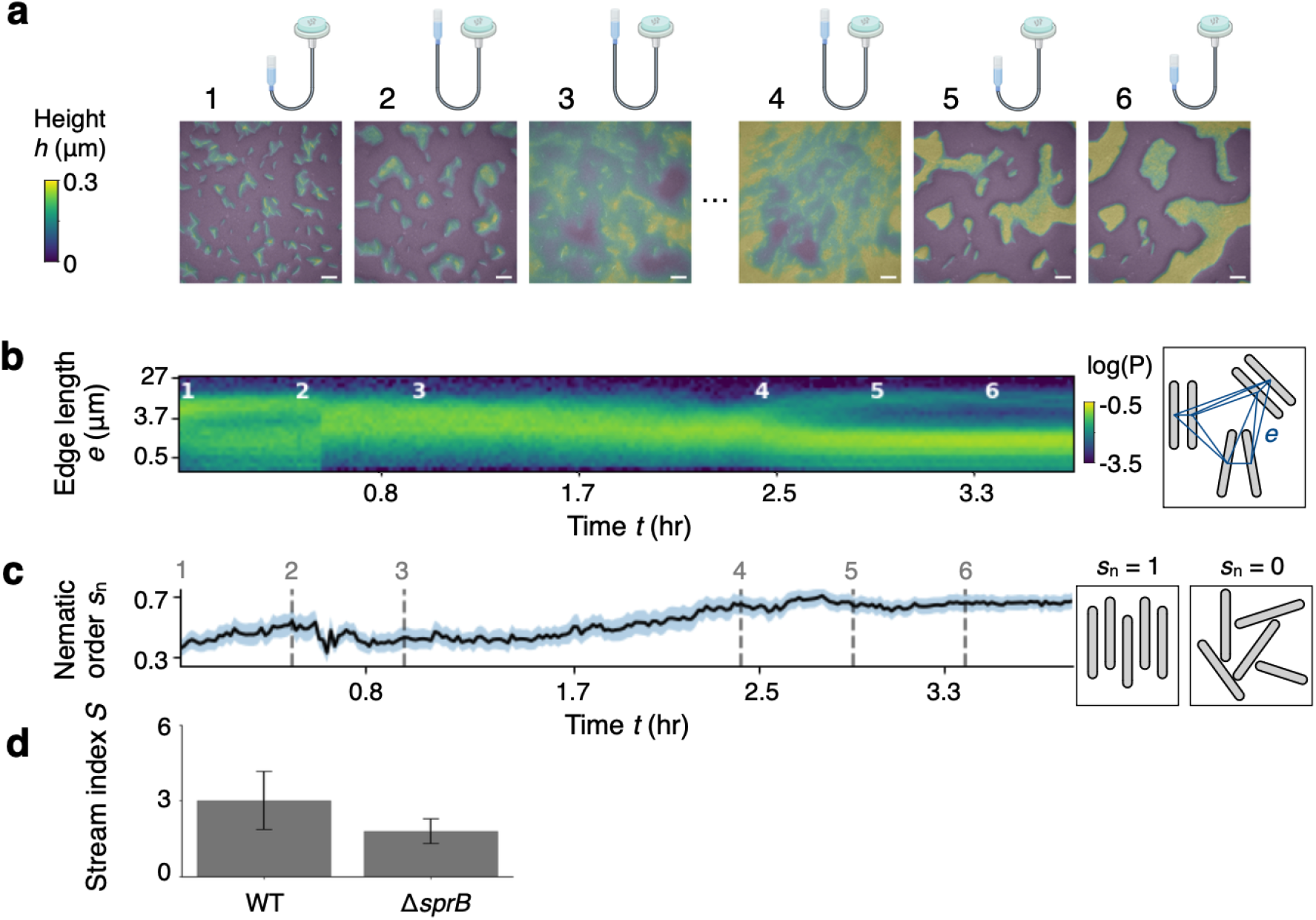
Water availability influences the organization of colonies of *F. johnsoniae*. (**a**) Experimental time series in which cells initially deposited onto an agarose pad are subjected first to increasing water availability until a gas phase is reached (snapshots 1–3). Hours later, water is drained from the substrate, and the colony goes back to forming tightly packed, nematically ordered groups and streams (snapshots 4–6). Time points (1-6) are marked in **b** and **c**. (b) Kymograph of the probability density function (P) of Delaunay edge lengths over time for the time series shown in **a**. Schematic on the right demonstrates how edges (blue lines) are calculated for a small population of cells (gray). At early times and late times in the experiment, low water availability at the surface constrains cells into tightly packed, well-separated groups. At intermediate times, high water availability permits cells to be further spaced apart and homogenizes the population. (**c**) Strength of nematic order *s*_n_, calculated within groups of 8 nearest-neighboring cells and averaged over the population at each time. Shaded regions indicate variance. Right: schematics showing a small population of cells with perfect nematic order (*s*_n_ = 1) or no order (*s*_n_ = 0). (**d**) Stream index values for wild-type *F. johnsoniae* and a non-motile mutant (Δ*sprB*). Error bars are standard deviations for each corresponding mean calculated over N=28 and 5 frames taken from 3 and 1 separate timelapses, respectively. Plots correspond to main Fig. 4.

**Supplementary Fig. 20.**
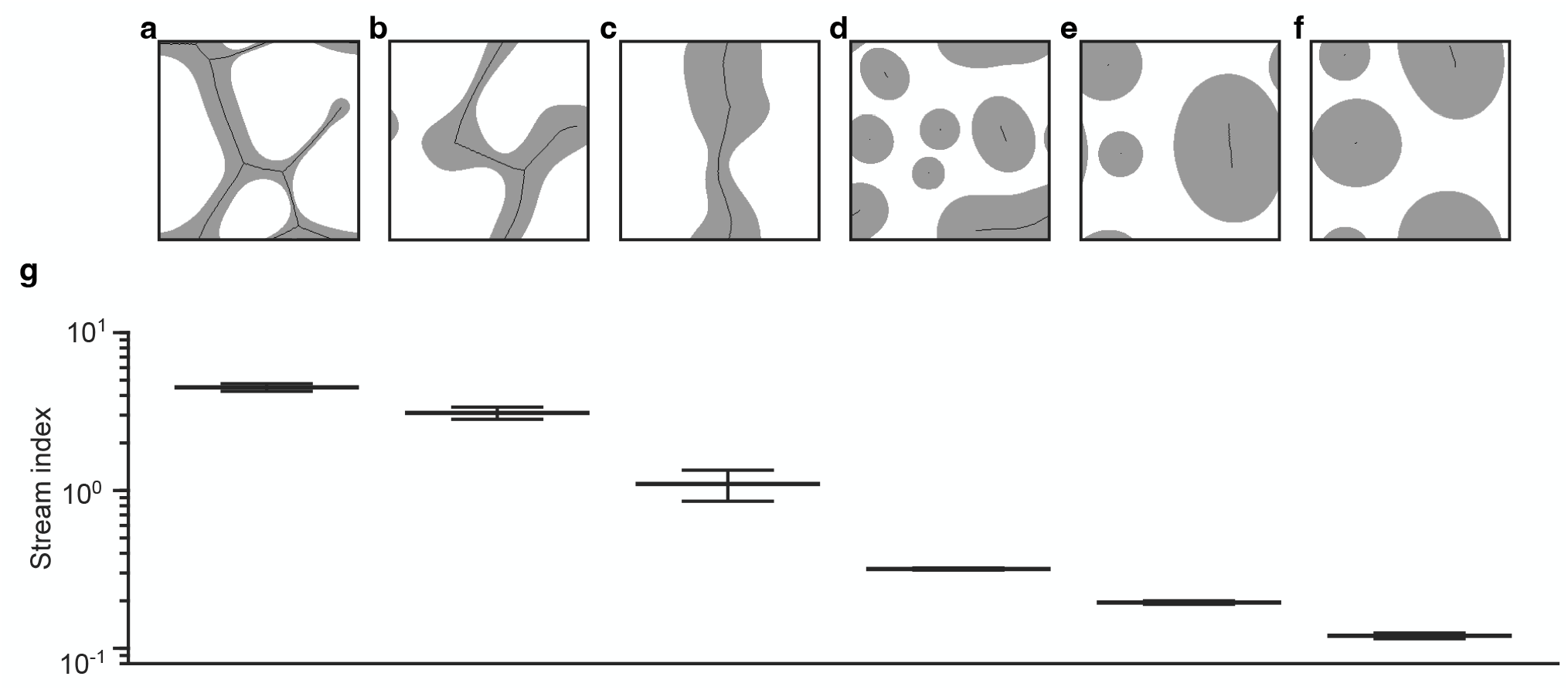
Quantifying the large-scale morphologies of cell populations. (**a**–**f**) Typical snapshots of the cell populations described by the continuum model with varying parameters (Sec. IV). White and gray denote voids and cell regions, respectively. Black curves indicate the skeletons of the cell regions as defined in Sec. IV C. Simulation parameters are: (**a**) *σ* = 1, *τ*_**ρ**_ = 10, (**b**) *σ* = 1, *τ*_**ρ**_ = 10^2^, (**c**) *σ* = 1, *τ*_**ρ**_ = 10^4^, (**d**) *σ* = 4, *τ*_**ρ**_ = 10^1^, (**e**) *σ* = 20, *τ*_**ρ**_ = 10^4^, (**f**) *σ* = 20, *τ*_**ρ**_ = 10^1^. (**g**) Stream indices for the corresponding simulations in panels **a**–**f**. Data are represented as mean ± std with respect to time for each simulation. In panels **a**–**c**, we set *K*_1_ = 1, *K*_2_ = 0.01, and *K*_3_ = 1. In panels **d**–**f**, we set *K*_1_ = *K*_2_ = *K*_3_ = 0. See Sec. IV for details.

**Supplementary Fig. 21.**
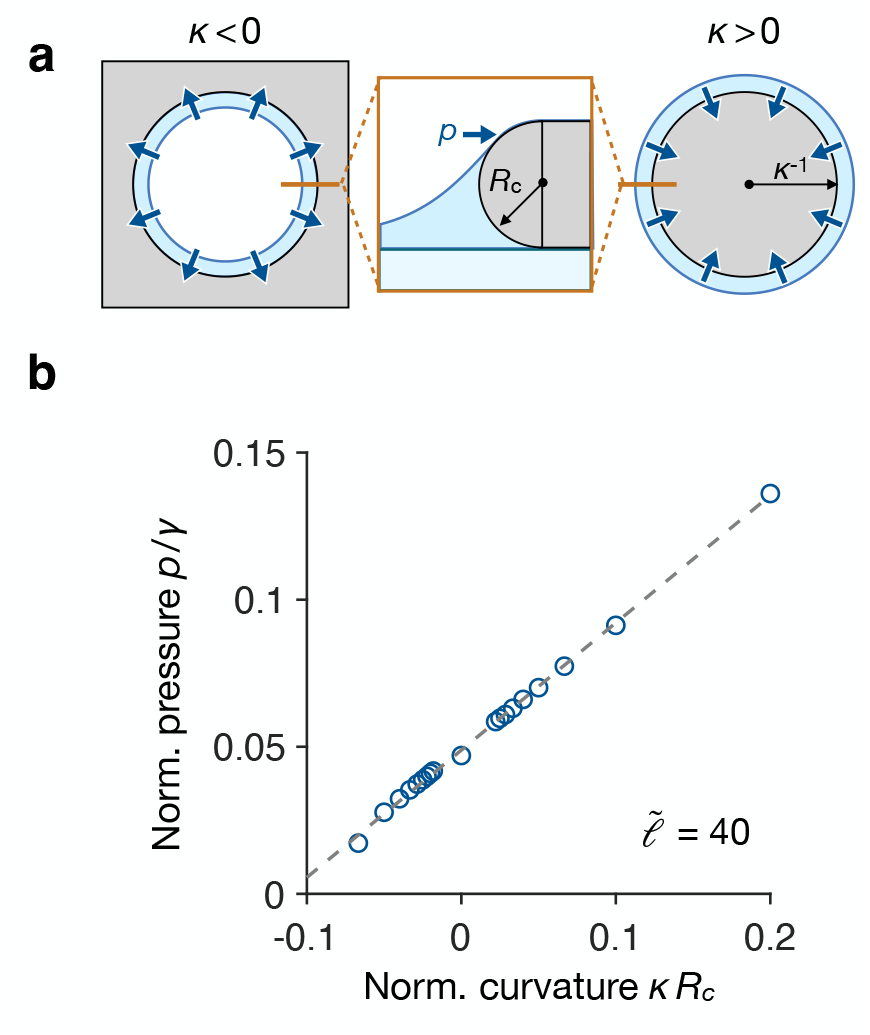
The in-plane capillary force acts as an Laplace pressure at large scales. (**a**) Schematic of the agent-based simulations. Gray denotes cell regions bounded by interfaces of curvature *κ*. Blue denotes the water. Blue arrows denote the in-plane pressure *p* exerted on the cells. *R*_c_ denotes the cap radius of each cell. The left and right panels show the top view schematics for *κ <* 0 and *κ >* 0, respectively. The middle panel shows a close-up cross section. (**b**) Normalized pressure *p/γ* at varying normalized curvatures *κR*_*c*_ for the designated parameter 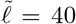. Note that *p* denotes a 2D pressure, defined as force per unit length, and hence has the same unit as *γ*. Circles denote simulation results, and the dashed line indicates a linear fit *p/γ* = *α*_1_*κR*_c_ + *α*_0_ with fitting parameters *α*_1_ = 0.433 and *α*_2_ = 0.05.

**Supplementary Fig. 22.**
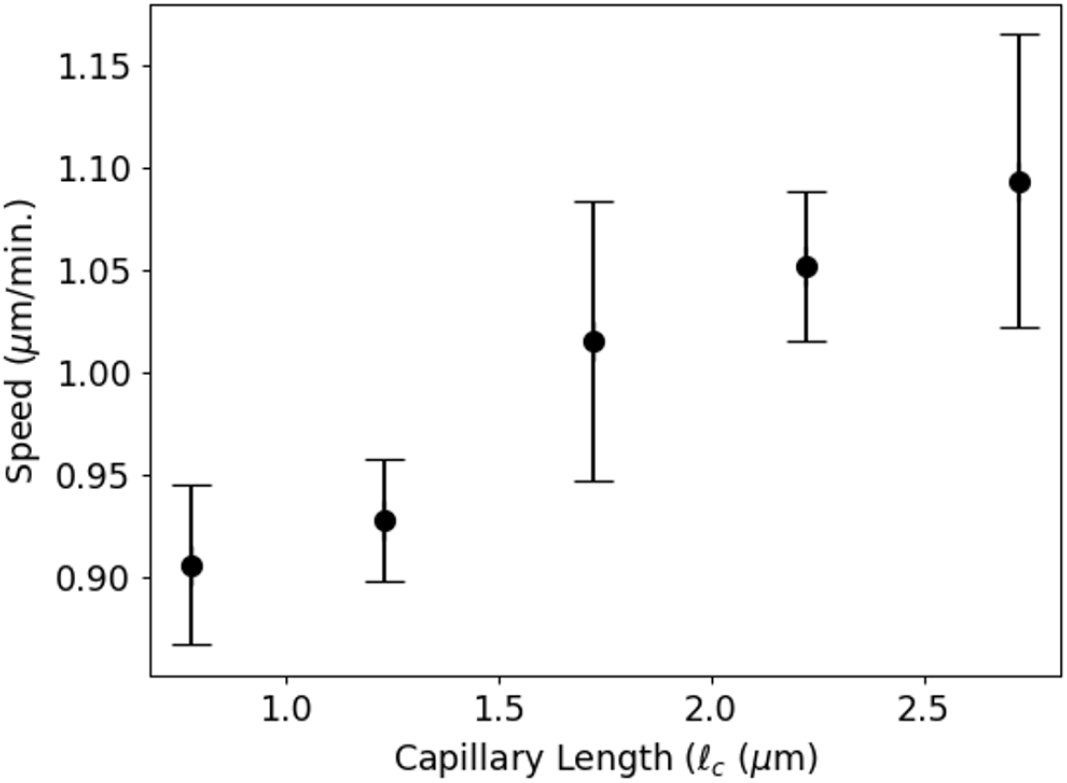
Speed vs. measured capillary length for single, motile *M. xanthus* cells.

**Supplementary Fig. 23.**
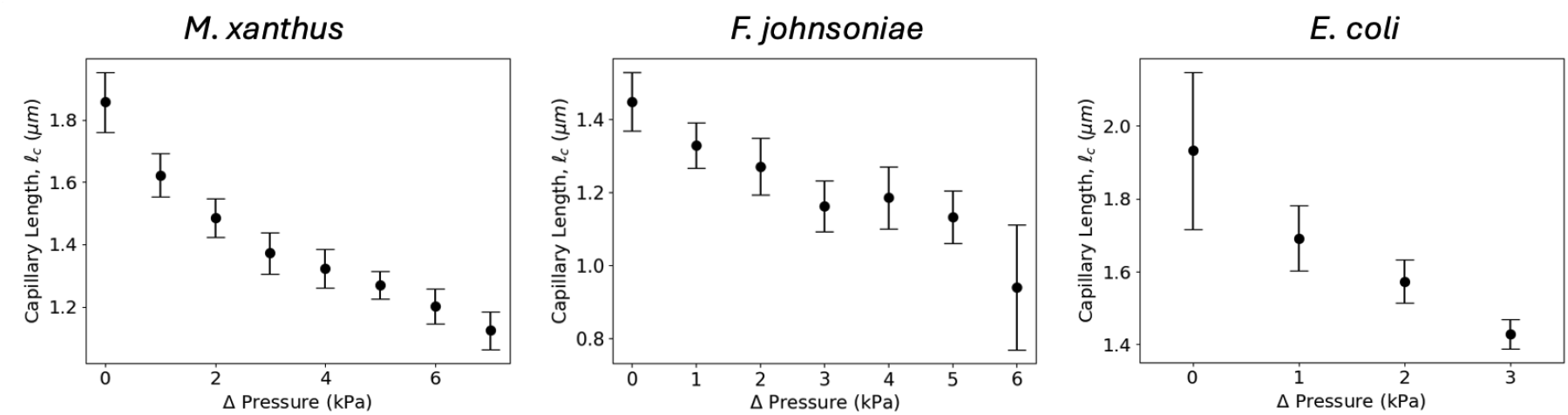
Exponential fits to measured meniscus profiles are anti-correlated with applied hydrostatic pressure. Single cells of the indicated species were deposited onto 1.5% v/v agarose pads and their wetting menisci measured (Capillary Length, *𝓁*_*c*_, y-axis on all graphs) at varying levels of applied hydrostatic pressure (ΔPressure, x-axis on all graphs).

**Supplementary Fig. 24.**
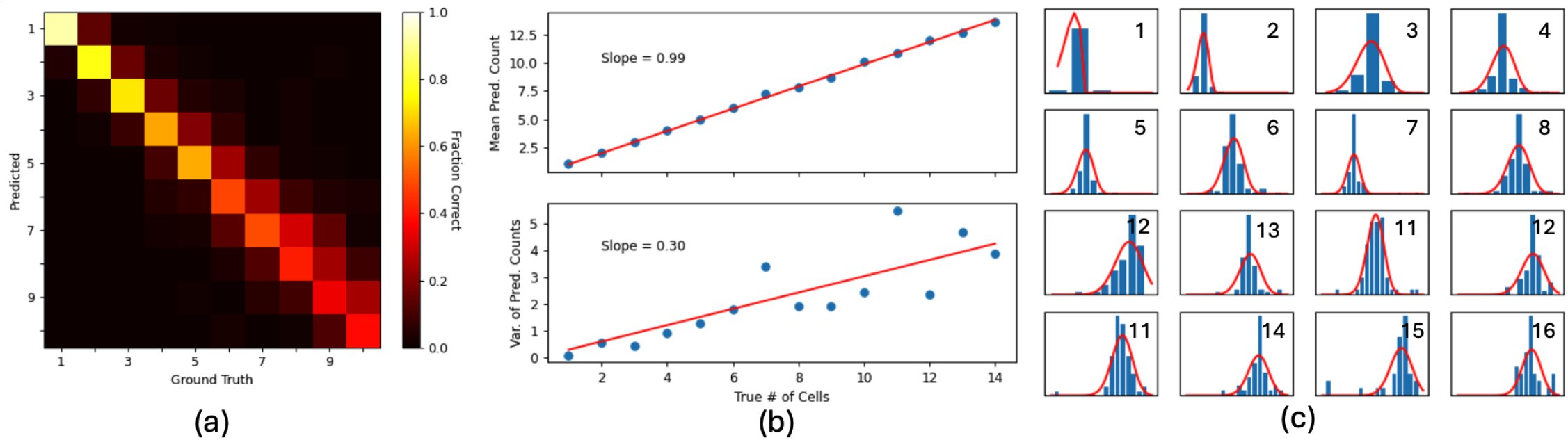
Calibration of the distribution of predicted counts for fixed group size. **(a)** Camera (RGB) image of dense groups of *F. johnsoniae* cells on an agarose pad, overlaid with the corresponding instance segmentation from our neural network. Region shown in (b) and (c) is shown with the white box. **(b**,**c)** Highlighted region of the original image (b) and the corresponding instance segmentation (c) for the region shown, boxed in (a). **(d)** Laser reflectance image of dense groups of *M. xanthus* cells on an agarose pad, overlaid with the corresponding instance segmentation from our neural network. Region shown in (e) and (f) is shown with the white box. **(e, f)** Highlighted region of the original image (e) and the corresponding instance segmentation (f) for the region shown, boxed in (d).

**Supplementary Fig. 25.**
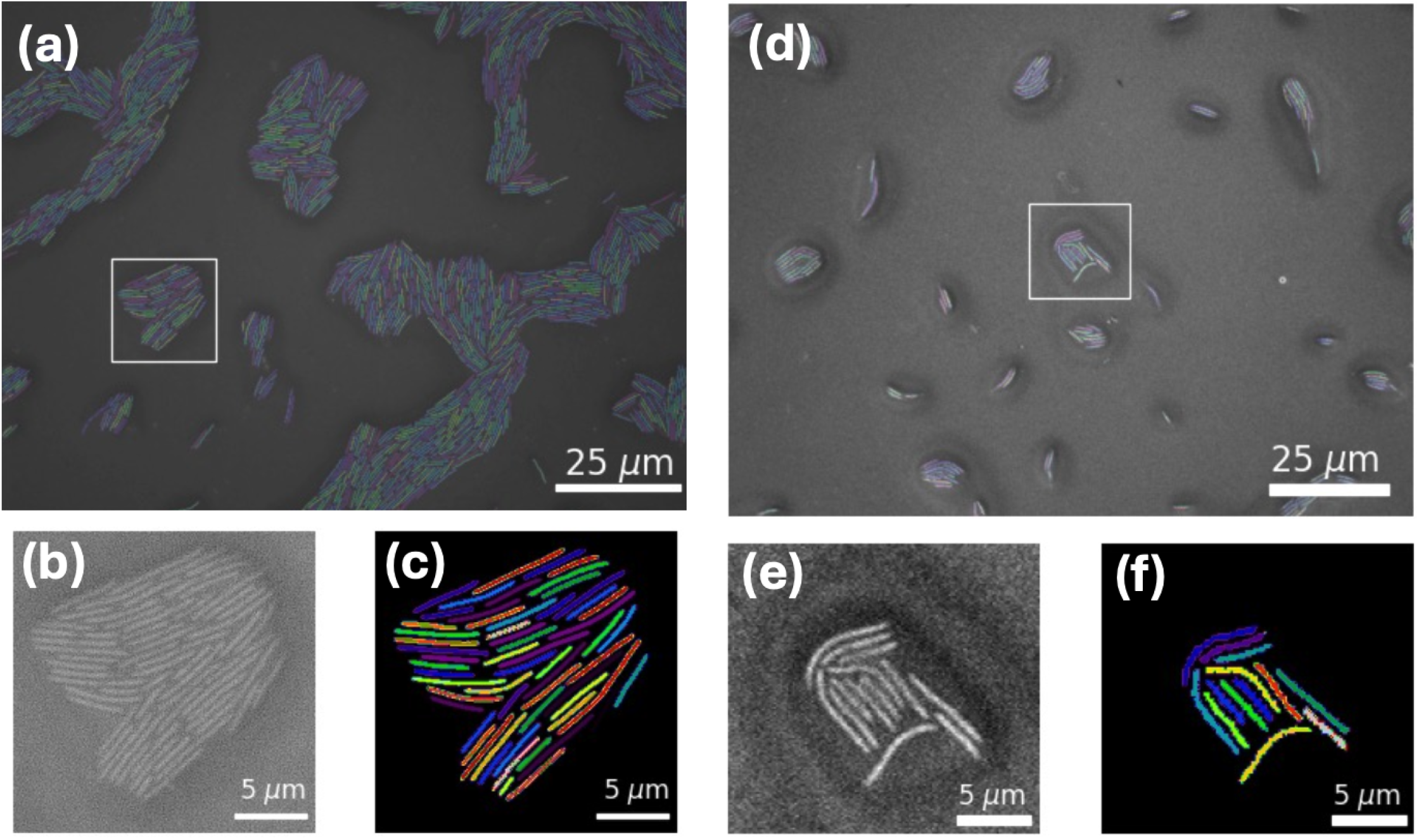
Representative segmentation results for *F. johnsoniae* and *M. xanthus*. **(a)** Heatmap indicated the fraction of all predicted counts for groups of size *N* (x-axis). **(b)** Linear regressions of the mean (top) and variance (bottom) of predicted counts for groups of size *N* (x-axis). Note that if the predicted counts were Poisson-distributed, we would expect the slope of the variance vs. ground truth regression to be 1. Thus, we conclude that counts are underdispersed. **(c)** Histograms of predicted cell counts for each collection of ground truth groups of size *N* (indicated in the top-right corner of each graph) up to groups of size *N* = 16. Red lines are fits to a generalized Poisson distribution [23] with fixed w=-0.94.

**Supplementary Fig. 26.**
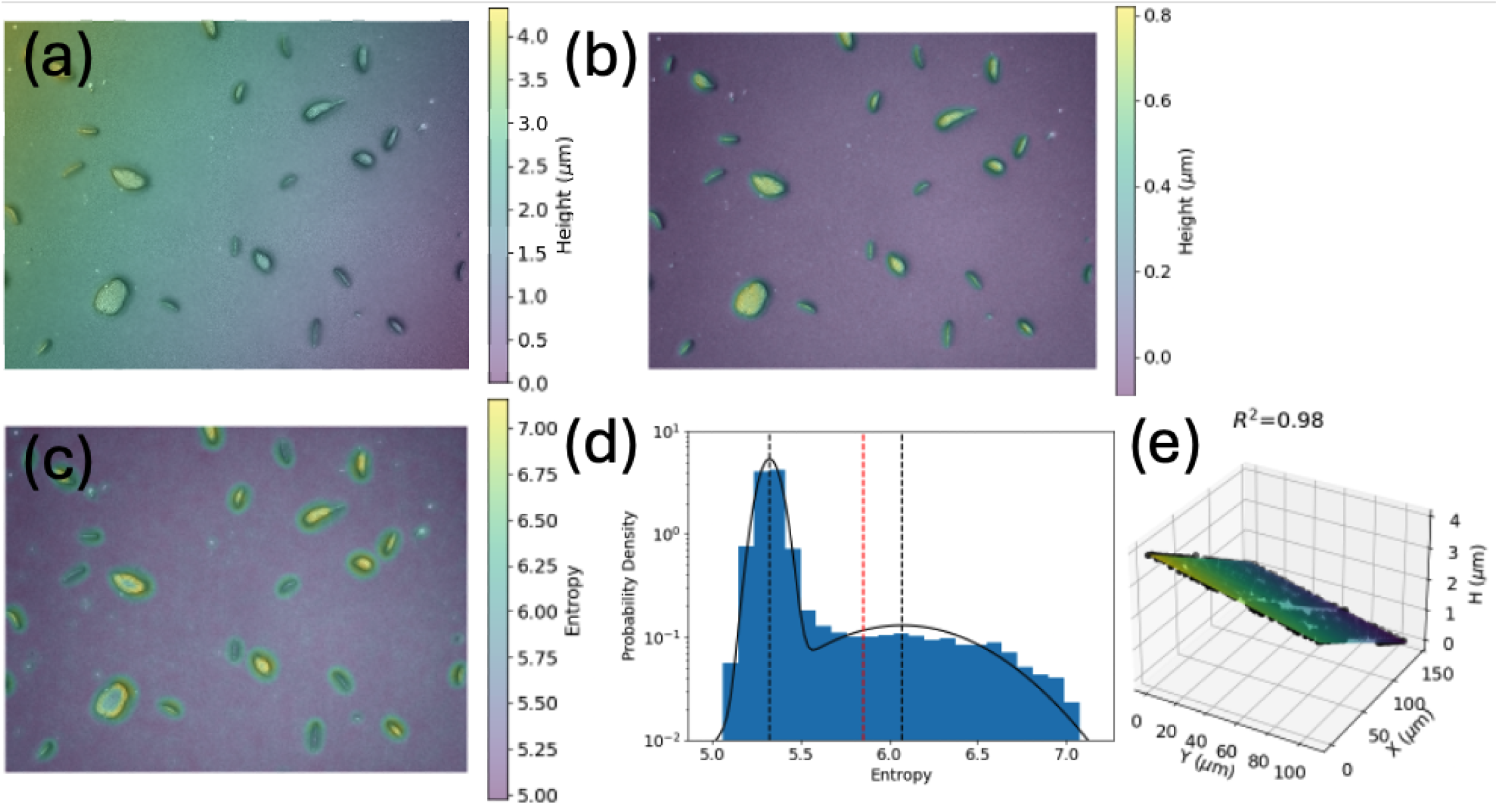
Procedure for correcting Keyence height images for sample tilt and surface unevenness. **(a)** Example uncorrected height height map overlaid on its corresponding reflectance image. Image is of small groups and single cells of *M. xanthus*. **(b)** (a) but with the corrected height map overlaid. **(c)** Output of entropy filtering the reflectance image of (a). **(d)** Histogram of entropy values from (c), with fit of the distribution of entropy values to a 2 component Gaussian mixture model (black, solid line). Black, dashed lines are the fitted means of the 2 components. Dashed red line is the threshold value, below which corresponding pixels are called as part of the “surface.” **(e)** Fit (solid surface) of raw height values (black points, subsampled 1:1000 from original data).

**Supplementary Fig. 27.**
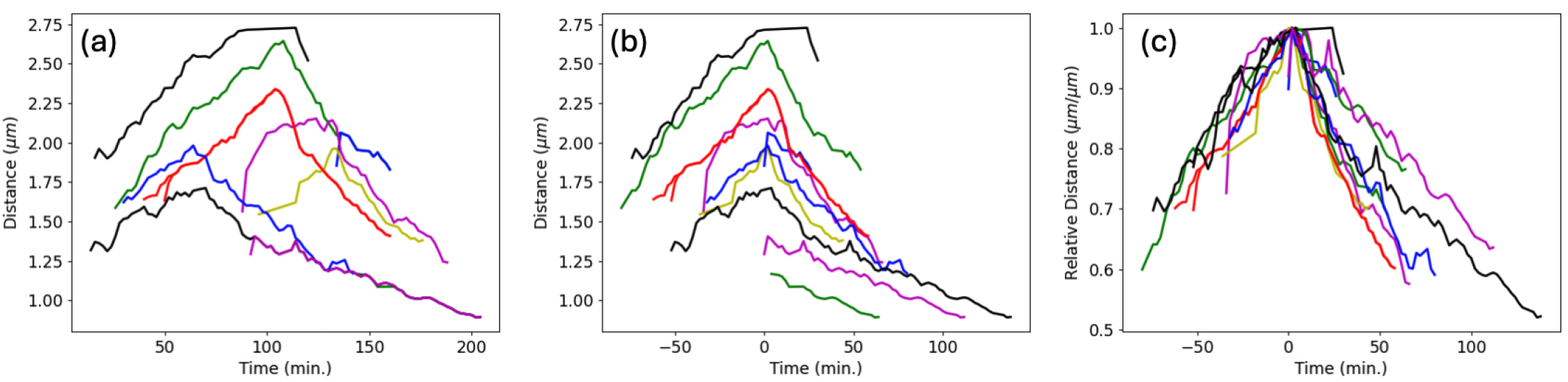
Calculation of relative centroid-to-centroid distance for non-motile cell pairs. **(a)** Smoothed traces of absolute centroid-centroid distance between pairs of cells. Each line represents a single pair of daughter cells. **(b)** Traces are aligned such that time *t* = 0 min. corresponds to the time at which the centroids are maximally-separated. **(c)** Relative centroid-centroid distance is calculated by dividing the distances in (b) by their value at *t* = 0 min.. The curve shown in Fig. 2i was computed by averaging, at each time, the data shown here.

## SUPPLEMENTARY VIDEOS

**Supplementary Video 1**. Movie for the time series shown in Fig. 2d, showing the merger of non-motile *M. xanthus* cells due to capillary attraction.

**Supplementary Video 2**. Movie showing merging and splitting of motile *M. xanthus* groups. Colored regions are groups of cells marked as a single groups. Overlaid numbers are the detected number of cells in that group.

**Supplementary Video 3**. Movie showing an example of simulated cells in the “gas” phase. Simulation parameters are 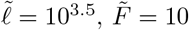 and 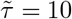. Color bar is the same as in main Fig. 3.

**Supplementary Video 4**. Movie showing an example of simulated cells in the “droplets” phase. Simulation parameters are 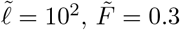 and 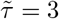. Color bar is the same as in main Fig. 3.

**Supplementary Video 5**. Movie showing an example of simulated cells in the “polar clusters” phase. Simulation parameters are 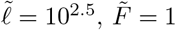 and 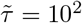. Color bar is the same as in main Fig. 3.

**Supplementary Video 6**. Movie showing an example of simulated cells in the “streams” phase. Simulation parameters are 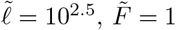 and 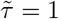. Color bar is the same as in main Fig. 3.

**Supplementary Video 7**. Movie showing an example of *M. xanthus* cells in the “gas” phase.

**Supplementary Video 8**. Movie showing an example of *M. xanthus* cells in the “droplets” phase.

**Supplementary Video 9**. Movie showing an example of *M. xanthus* cells in the “polar clusters” phase.

**Supplementary Video 10**. Movie showing an example of *M. xanthus* cells in the “streams” phase.

**Supplementary Video 11**. Movie for the time series in Fig. 3f, showing neighbor exchange in the “stream” phase.

**Supplementary Video 12**. Movie for the time series shown in Fig. 4a, showing the large-scale morphologies of *M. xanthus* cells upon changes in water availability.

**Supplementary Video 13**. Movie showing the large-scale morphologies of *F. johnsoniae* cells upon changes in water availability.

## Footnotes

1 This expression only works for *l*_*γ*_ ⩾ *R*_c_. When *l*_*γ*_ *< R*_c_, i.e., the central angle *θ*^∗^ of the two circular arcs in Eq. (S5) become larger than *π/*2, and both *±* signs should be kept in front of the square-root terms.

2 Following previous work [4], we define cell-cell distance as the smallest distance between the centerlines of the cell cylinders.

3 Note that the data shown in Fig. 3e was taken using a high resolution scan as well as an accompanying RGB image (see VI A, VII)

4 Note that touching is not necessarily an all-to-all relation. If region **A** touches **B** and **C** but **B** and **C** do not touch, the resultant group would include **A, B**, and **C**.

